# DNA-directed nanometer-scale integration of single colloidal quantum emitters with on-chip photonics

**DOI:** 10.1101/2025.01.23.634416

**Authors:** Xin Luo, Shelly Ben-David, Luigi Ranno, Tara Sverko, Natalie Ngoh, Jae Young Lee, Peter F. Satterthwaite, Nicholas Sbalbi, Annette Jones, Chi Chen, Moungi G. Bawendi, Juejun Hu, Farnaz Niroui, Robert J. Macfarlane, Mark Bathe

**Affiliations:** Department of Biological Engineering, Massachusetts Institute of Technology, Cambridge, MA, 02139, USA; Department of Materials Science and Engineering, Massachusetts Institute of Technology, Cambridge, MA, 02139, USA; Department of Electrical Engineering and Computer Science, Massachusetts Institute of Technology, Cambridge, MA, 02139, USA; Department of Chemistry, Massachusetts Institute of Technology, Cambridge, MA, 02139, USA

## Abstract

Incorporating colloidal quantum emitters into integrated photonics would greatly advance quantum optics. However, placing individual sub-10 nm colloidal particles onto micron-sized photonic structures with nanometer-scale precision remains challenging. We introduce topography-modulated origami placement, which uses DNA origami to place colloidal nanomaterials within lithographically defined, shape-matched resist wells. The method patterns origami templates onto photonic chips with high yield and nanometer-scale precision. A soft- silicification-passivation step stabilizes deposited origamis while preserving DNA binding sites that selectively attach and align colloidal quantum rods, controlling their position and emission polarization. Placed quantum rods demonstrated room-temperature single-photon emission. We integrated individual quantum rods with waveguides, micro-ring resonators, and bullseye microcavities, and achieved cavity-coupled single-photon emission. This approach offers a general platform for integrating colloidal quantum materials into photonic systems.

---

Advances in nanophotonics, particularly quantum photonic integrated circuits (PICs), demand precise and selective nanoscale spatial control over the fabrication of optically functional materials^1,2^. While conventional top-down methods often face resolution and yield limitations in fabricating quantum emitters^2,3^, bottom-up molecular and colloidal synthesis can scalably produce diverse optical nanomaterials with nanometer- to atomic-scale features and tunable properties^4,5^. However, colloidal materials are randomly dispersed in solution, whereas applications in quantum sensing, communication, and computing require deterministic, single-particle incorporation of colloidal quantum emitters into optical and electronic structures on PICs^5,6^. Current methods predominantly rely on inefficient processes such as post-deposition fabrication^7,8^ or lithography-guided deposition with stochastic site occupancy^9,10^, while capillary-force-assisted placement is typically limited to large nanoparticles (NPs) on flat substrates with size-matching traps and lacks programmable selectivity^11,12^. Thus, achieving precise, selective, and scalable integration of sub-10 nm colloidal quantum emitters into topographically complex photonic structures with nanoscale control remains an unsolved challenge.

DNA nanotechnology, particularly DNA origami, has emerged as a scalable bottom-up approach to organize nanoscale optical materials, including molecular emitters^13,14^, metallic NPs^15^, upconverting NPs^16^, quantum dots (QDs)^17,18^ and nanodiamonds^19^ into complex hierarchical structures with nanometer-scale accuracy. The DNA origami placement (DOP) method further demonstrated positioning of 2D^20–22^ and 3D^23^ DNA origamis onto landing pads defined by electron beam lithography (EBL), successfully organizing dye molecules^20,21^ and gold NPs^23,24^. However, quantum emitters such as nanodiamonds and quantum rods (QRs) are unstable in the divalent-cation-rich conditions required for DOP^17,19^. In addition, while alternative lithographic methods have been explored for origami placement on various flat substrates^25–27^, limited progress has been made with practical PIC structures, as their micron-scale topographical features can hinder precise placement and lead to random trapping of colloidal nanomaterials.

Here, we introduce Topography-Modulated Origami Placement (TMOP), which combines lithographically defined confinement with DNA origami templating to deterministically integrate individual colloidal quantum materials into on-chip photonic devices (Fig. 1). TMOP uses EBL to pattern origami-shape-matched wells in a resist layer, where topographical confinement, origami design, and ionic conditions control the origami placement position and orientation at the nanometer scale. Hybridization overhangs on the deposited origamis then capture DNA-functionalized nanomaterials. We first use gold NPs, which tolerate the divalent-cation-rich placement conditions, to show that TMOP controls the number and position of gold nanospheres (AuNSs) and the orientation of anisotropic gold nanorods (AuNRs). A soft-silicification-passivation strategy stabilizes TMOP origamis while preserving their binding overhang accessibility, enabling assembly of polarization-aligned QRs under the divalent-cation-free conditions required for their colloidal stability. We deterministically place individual QRs on silicon nitride (SiN_x_) waveguides, micro-ring resonators, and bullseye microcavities, and demonstrate cavity-coupled single-photon emission from a single QR in a titanium dioxide (TiO_2_) bullseye cavity. TMOP thus provides a general platform for colloidal quantum emitters in integrated quantum photonics.

**Fig. 1.**
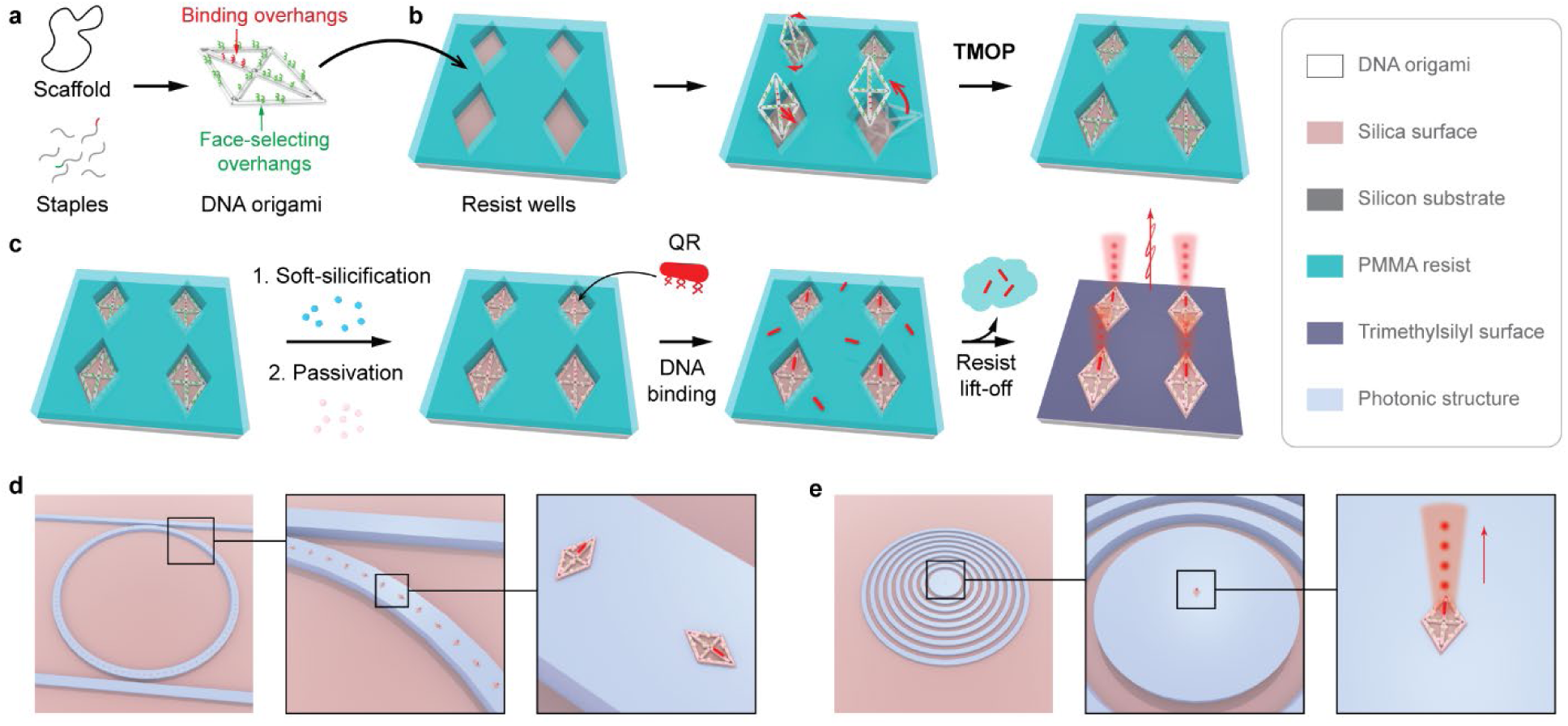
Overview of TMOP for deterministic integration of colloidal quantum emitters with on-chip photonics. **a**, Design and assembly of a planar DNA origami with binding overhangs (red) and face-selecting overhangs (green). **b**, Topography-modulated origami placement at the base of shape-matched wells in a PMMA resist layer with origami binding overhangs facing up. **c**, Soft-silicification and passivation of the placed origamis for Mg^2+^-free, DNA-directed binding and orientation of colloidal QRs. Resist lift-off removes background and yields clean QR arrays with controlled position and emission polarization. **d**, Deterministic integration of oriented QRs on a micro-ring resonator. **e**, Placement of a single QR at the center of a bullseye cavity for device-coupled single-photon emission.

## Topography-modulated origami placement

TMOP’s shape-matched resist wells provide two key advantages over previous approaches^20,21^. First, the resist layer acts as a physical barrier against random attachment of nanomaterials to the background, and any accumulated material is removed by resist lift-off. Second, topographical confinement further regulates origami orientation and minimizes origami aggregation. We fabricated origami-shaped wells in a poly(methyl methacrylate) (PMMA) resist layer by standard EBL (see Methods; Supplementary Fig. 1) and used 2D six-helix bundle (6HB) wireframe origamis (Supplementary Fig. 2 and Supplementary Note 1) as the model platform because of their planarity and rigidity^28,29^. One side of the 2D origami carried single-stranded DNA overhangs of two types (Supplementary Table 1): the NP-binding sequence (Fig. 1a, red) and the 20-nucleotide polythymidine sequence (Fig. 1a, green), which act as an entropic brush that prevents this side from binding the substrate^17,21^ and keeps the binding overhangs accessible.

We first performed TMOP with 35 mM MgCl_2_ using a rhombic 6HB wireframe origami with diagonal axes of approximately 75 nm and 125 nm. Notably, TMOP worked with crude, unpurified origami directly from folding, avoiding the material loss and aggregation associated with origami purification^30^. In a typical TMOP process, a droplet of crude origami solution was incubated on a chip bearing EBL-fabricated rhombic wells (Fig. 2a, left; Supplementary Fig. 3) and washed to remove excess DNA (see Methods). Atomic force microscopy (AFM) confirmed origami positioning at the well bases (Fig. 2a, middle; Supplementary Fig. 4), and subsequent PMMA lift-off removed the resist and any background residue, leaving clean origami arrays (Fig. 2a, right; Supplementary Fig. 5). Silicification before lift-off rendered the origamis robust and visible by scanning electron microscopy (SEM). We fabricated a 55 µm × 50 µm silicified TMOP array, comprising subarrays of origami with customized orientations and spacings (Fig. 2b), showing no background contamination after lift-off (Supplementary Figs. 6 and 7). We also applied TMOP to a ladder-shaped 6HB wireframe origami and a single-layer triangular origami, producing large-scale arrays with high yields (Supplementary Figs. 8–12). TMOP’s bath-based workflow could be readily adapted for wafer-scale processing (Supplementary Fig. 13 and Supplementary Note 2).

**Fig. 2.**
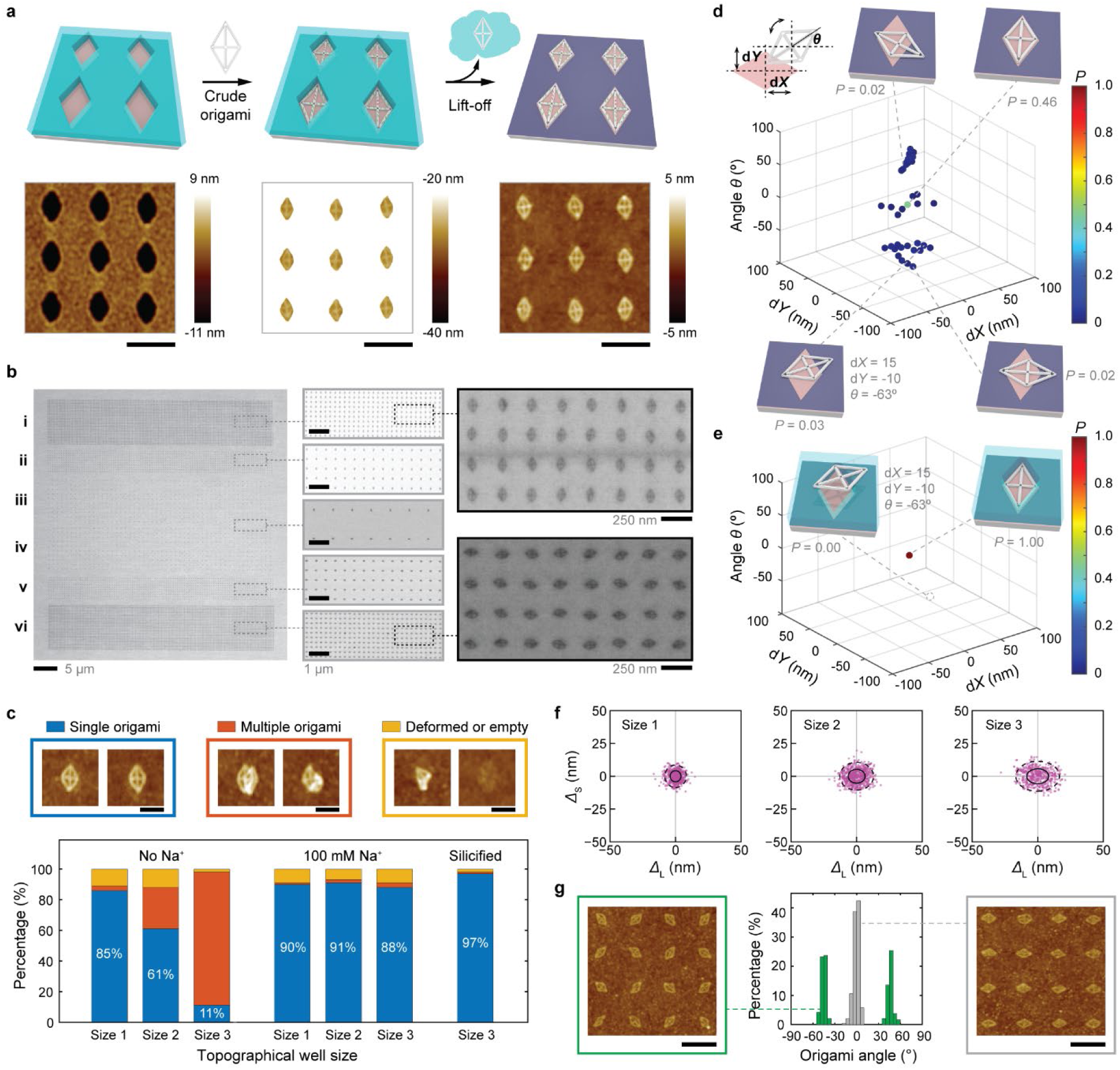
Topography-modulated placement of 6HB wireframe origami. **a**, Schematics (top) and AFM images (bottom) of TMOP with a rhombic origami. Left: well; middle: origami at the well base; right: array after lift-off. **b**, SEM images of silicified rhombic origamis oriented vertically (i, ii, iii) and horizontally (iv, v, vi) and spaced 250 nm (i, vi), 500 nm (ii, v) or 1 µm (iii, iv) apart. **c**, Placement statistics for well sizes 1 to 3 (Methods) with no Na^+^ (left), 100 mM Na^+^ (middle), or 100 mM Na^+^ and silicification (right). Insets: single origamis (blue), multiple origamis per well (orange), and deformed origamis or empty wells (yellow). **d,e**, Simulated probabilities (*P*) of origami conformations (d*X*, d*Y*, *θ*) without (d) or with (e) wells; insets show the corresponding conformations. **f**, Displacement of single origamis from their lattice sites along the long (*Δ*L) and short (*Δ*S) axes, for well sizes 1 to 3 (*N* = 514, 652 and 487); solid and dashed ellipses mark 1σ and 2σ, respectively. **g**, Origami orientation distributions and AFM images for wells oriented at 0° (right, gray) or ±45° (left, green). Scale bars, 250 nm (a,g) and 100 nm (c).

To examine how topographical confinement regulates origami placement, we prepared rhombic wells of increasing size (sizes 1–3; AFM measured openings of 97 × 135, 106 × 150 and 114 × 161 nm, respectively; see Methods; Supplementary Fig. 3). As well size increased from size 1 to 3, the yield of correctly placed single origamis decreased from 85% to 11% (*N* = 288 each), while the occurrence of wells containing multiple origamis increased from 3% to 86% (Fig. 2c; Supplementary Fig. 14), indicating that origami–substrate binding was strong enough for multiple origamis to occupy the same wells under 35 mM Mg^2+^. To modulate origami binding kinetics, we added 100 mM Na^+^ to the placement solution to attenuate Mg^2+^-mediated origami–substrate interactions^17,31,32^, which suppressed multiple origami binding and increased single-origami placement yields to ≈90% across all well sizes (*N* = 576 each; Supplementary Fig. 5). Silicification before resist lift-off further improved the yield to 97% (size 3, *N* = 576; Fig. 2c; Supplementary Fig. 15).

To understand the topographical confinement mechanism, we used an energy-landscape model and kinetic simulations (see Methods; Supplementary Note 3). Without wells, numerous local energy minima away from the correct conformation limited the probability of correct placement to ≈0.46 (Fig. 2d; Supplementary Figs. 16 and 17), whereas well confinement eliminated these local minima, leaving a single aligned conformation (*P* = 1.00; Fig. 2e; Supplementary Fig. 18). The simulations further suggested that origami rigidity is important in reducing kinetically trapped conformations (Supplementary Note 4). Similar effects were observed in simulations of ladder-shaped and triangular origamis (Supplementary Figs. 19 and 20). Tighter confinement also improved origami placement precision (Fig. 2f), as site-to-site displacements from the fitted lithographic lattice showed standard deviations of 4.1 ± 0.2 nm and 4.3 ± 0.1 nm along the origami long and short axes, respectively, for well size 1 (*N* = 514), and 8.3 ± 0.2 nm and 5.7 ± 0.1 nm for well size 3 (*N* = 487) (see Methods; Supplementary Fig. 21). Well confinement also directed the rhombic origami orientation, defined as the angle of the rhombus long diagonal axis between −90° and +90°. In a horizontal rhombic well array, 81% of origami monomers oriented horizontally within 0 ± 5° following TMOP (standard deviation 4°, *N* = 245; Fig. 2g, gray; Supplementary Fig. 14a). In contrast, an array of alternating +45° and −45° wells led to 86% of origamis oriented within ±5° of their target orientation (standard deviations 4° and 3°, *N* = 236; Fig. 2g, green; Supplementary Fig. 22).

## Controlling colloidal particle number and orientation

Deterministic integration requires control over not only where materials land, but also how many occupy each site and how anisotropic particles are oriented. We first established these capabilities with gold NPs, which remain stable under origami placement conditions. While lithography-guided direct deposition can place gold NPs^33,34^, DNA origami can program their number, type and relative positions with nanometer-scale precision^35^, enabling precisely engineered functionalities^15^. Previous origami-directed placement of gold NPs on chips, however, has been limited to AuNSs on flat surfaces, which can suffer from aggregation and non-specific adsorption^23,24,36^. Here, we demonstrate the clean arrangement of both AuNSs and orientation-controlled AuNRs using TMOP.

We designed and assembled ladder-shaped 6HB wireframe origamis with either one (Fig. 3a, bottom left) or two (Fig. 3b, top) binding sites bearing overhangs complementary to the DNA sequence on functionalized AuNSs. After origami placement and AuNS incubation (see Methods), AFM showed primarily single or paired AuNSs at the well bases, respectively (Fig. 3a). Random AuNS deposition on the resist background (Supplementary Figs. 23 and 24) was removed by lift-off, leaving clean arrays of origami-directed single AuNSs or AuNS dimers (Fig. 3a,b; Supplementary Figs. 25 and 26), with placement yields of 86% and 75%, respectively (*N* = 432 each; Fig. 3c), surpassing previously reported yields^24^.

**Fig. 3.**
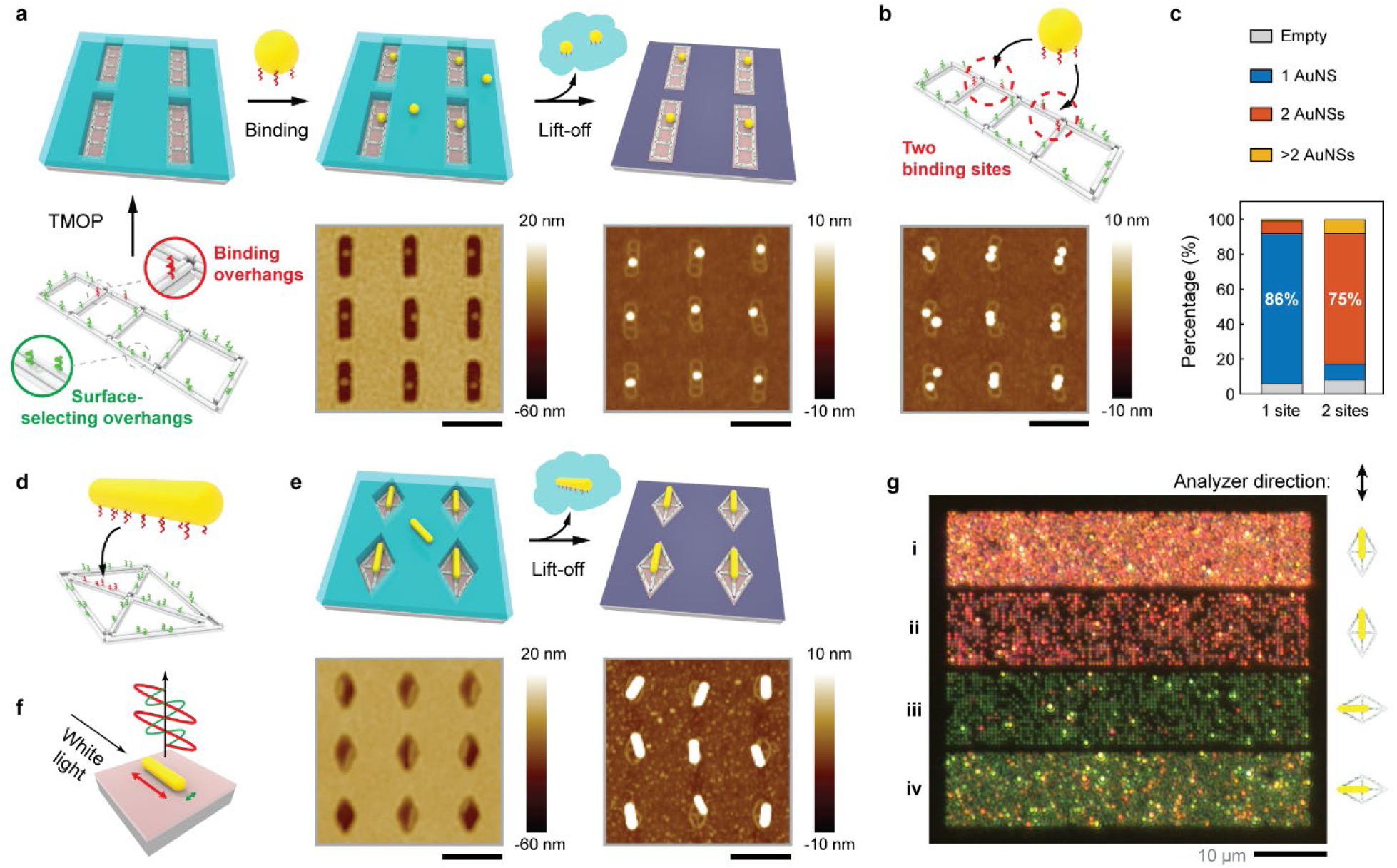
TMOP-directed array of plasmonic materials. **a**, Schematic of AuNS arrangement templated by ladder-shaped origami bearing one binding site (three overhangs), and AFM images of AuNSs within wells (left) and after lift-off (right). **b**, Ladder-shaped origami with two binding sites and AFM image of the resulting AuNS dimer array. **c**, Statistics of AuNS monomers and dimers placed with origamis bearing one or two binding sites. **d**, Linearly arranged binding overhangs on rhombic origami for AuNR binding and alignment. **e**, Schematic and AFM images of TMOP-directed AuNR arrangement within the wells (left) and after lift-off (right). **f**, Polarization- and wavelength-dependent light scattering from an AuNR. **g**, Dark-field microscopy image of TMOP-directed AuNR arrays with vertical (i, ii) or horizontal (iii, iv) orientations and 500 nm (i, iv) or 250 nm (ii, iii) spacings, observed through a vertically aligned analyzer under unpolarized illumination. AFM scale bars, 250 nm.

Customized overhang arrangement further allows TMOP to orient anisotropic nanomaterials such as AuNRs, which are versatile plasmonic structures with strong, tunable, and anisotropic optical properties^37^. Here we demonstrated TMOP-directed arrangement of AuNRs (75 nm × 18 nm) into arrays with orthogonal orientations using rhombic origamis with five binding overhangs along the long axis (Fig. 3d). AFM confirmed AuNRs on the origamis before and after lift-off, with an individual AuNR placement yield of 68% (*N* = 800; Fig. 3e; Supplementary Figs. 27 and 28). Deviations from ideal alignment are attributed to capillary-force-driven reconfiguration during drying (Supplementary Fig. 29 and Supplementary Note 5).

Because the longitudinal and transverse plasmon resonances of AuNRs are excited by light polarized along different rod axes at corresponding wavelengths, aligned AuNR arrays produce orientation-dependent scattering^38^ (Fig. 3f). We fabricated a 50 µm × 45 µm array comprising four strips of vertically or horizontally oriented AuNRs (Fig. 3g; Supplementary Figs. 28 and 30). Under unpolarized incident light and polarized collection, vertically aligned AuNRs (i and ii) primarily scattered red light along the vertical direction (i.e., the analyzer direction) arising from longitudinal plasmon resonance, while horizontally aligned AuNRs scattered primarily green light from transverse plasmon resonance (Fig. 3g). Under polarized incident light without an analyzer, AuNR arrays appeared red, green, or mixed depending on their orientation relative to the incident light polarization (Supplementary Fig. 31). This enables patterned, polarization-controlled coloration (Supplementary Fig. 32), with potential applications in metasurfaces^39^ and optical data storage^40^.

## TMOP-directed arrangement of single-photon emitters

Colloidal quantum emitters such as QDs and QRs combine scalable synthesis with tunable optical and electronic properties, making them attractive for electronics^41^, light-emitting devices^42^ and quantum information science^43^. QRs in particular offer morphology-dependent emission polarization (along the rod axis) for efficient coupling with photonic devices^44,45^. However, unlike gold NPs, DNA-functionalized QRs typically aggregate at the high Mg^2+^ concentrations required for origami placement. Here, we developed a soft-silicification-passivation method that stabilizes TMOP origamis under Mg^2+^-free conditions, enabling the arrangement of Mg^2+^-sensitive colloidal quantum emitters such as QRs (Fig. 4a).

**Fig. 4.**
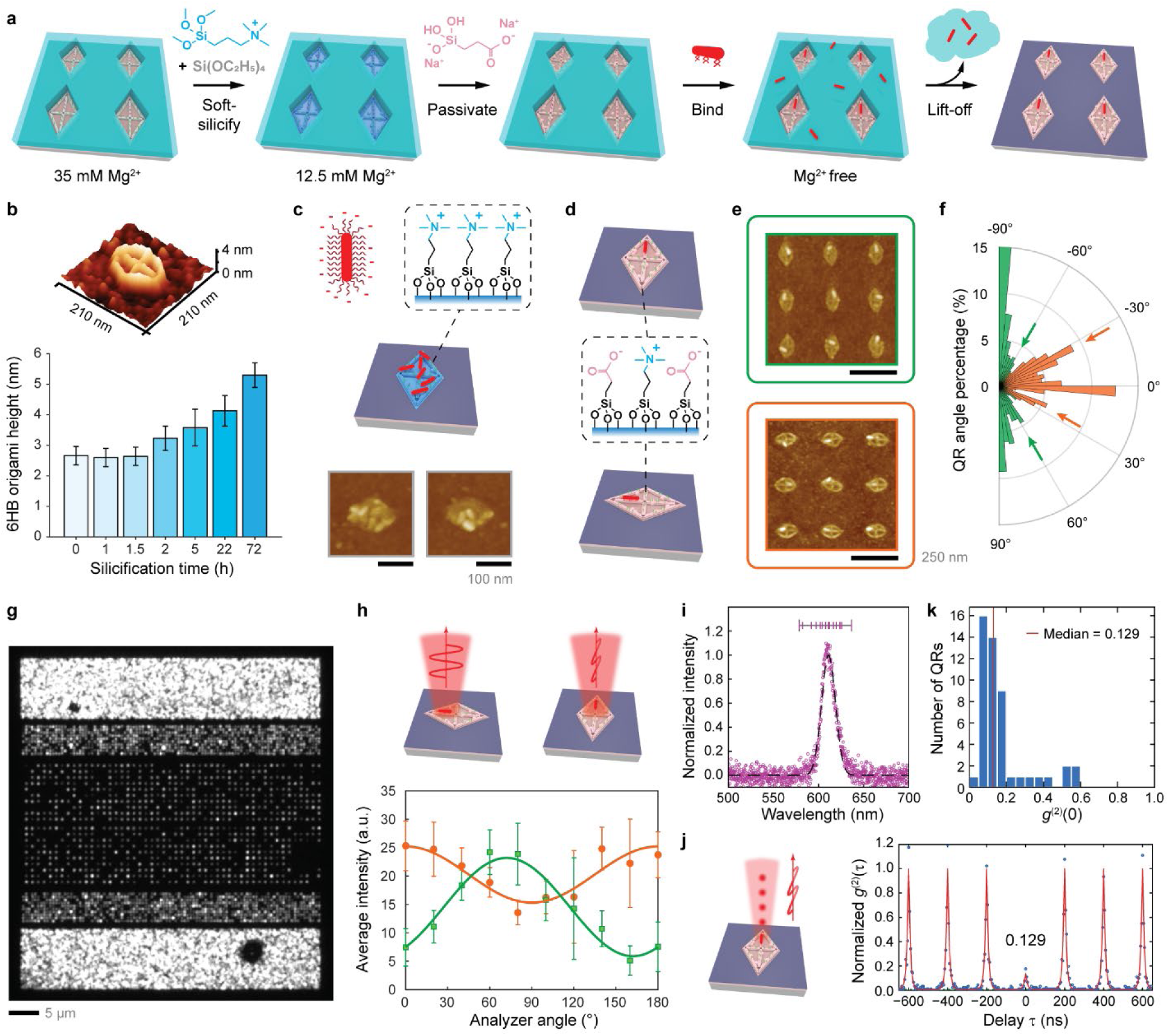
TMOP-directed array of single-photon emitters. **a**, Soft-silicification-passivation workflow: soft-silicification, passivation, QR binding, and lift-off. **b**, Dry AFM origami heights versus silicification time (mean ± s.d.; *N* = 39, 46, 43, 40, 30, 39 and 22 origamis measured at 0, 1, 1.5, 2, 5, 22 and 72 h, respectively). Inset: 3D height profile after 1.5 h. **c**, Schematic and AFM images of random, electrostatic QR attachment to silicified origami. **d**, Illustration of origami surface charge and chemistry after soft-silicification-passivation. **e**, AFM images of single QRs bound to vertically (top, green) and horizontally (bottom, orange) aligned origamis. **f**, QR orientation distributions on vertical (green) and horizontal (orange) origamis; arrows mark major deviations from design. **g**, Widefield PL image of a TMOP-directed QR array (maximum projection of 100 frames, 100 ms exposure each). **h**, Average PL intensity versus analyzer angle for vertically (green) and horizontally (orange) aligned QR arrays. Points, mean ± s.d. of six repeated confocal scans of the same array at each angle; lines, sinusoidal fits. **i**, Representative PL spectrum of a single QR (purple, raw data; black, Gaussian fit with *λ*0 = 611.1 nm and full width at half maximum (FWHM) = 18.1 nm). Horizontal bar: fitted peak positions of all emitters measured (*N* = 18). **j**, Schematic and second-order correlation trace, *g*(2)(*τ*), of the QR with the median *g*(2)(0) value (0.129), under pulsed excitation at room temperature. Blue points, data; red line, fit. **k**, Distribution of the *g*(2)(0) of QRs measured across the array (*N* = 49).

We silicified TMOP-directed origamis within the resist wells to secure their attachment while preserving access to the NP binding overhangs (see Methods). AFM showed that origami height remained around 2.6 nm for up to 1.5 h of silicification and then increased to 5.3 ± 0.4 nm over 72 h, indicating rigidification and silica deposition (Fig. 4b; Supplementary Fig. 33). We found that silicification as short as 1.5 h, termed soft-silicification (compared with days of silicification in previous work^23,46,47^), could ensure stable origami attachment under Mg^2+^-free conditions while preserving binding-strand accessibility. However, DNA-functionalized QRs directly incubated with silicified origamis attached randomly, likely through non-specific electrostatic attraction to the positively charged silicified surface (Fig. 4c; Supplementary Fig. 34). Passivating the positively charged surface with a negatively charged silane molecule restricted QR binding to DNA hybridization (Fig. 4d), giving individual QRs at each TMOP origami site with a 79% single-QR placement yield (*N* = 720; Fig. 4e; Supplementary Figs. 35 and 36). Orientational analysis showed that of single QRs, 73% oriented within 90 ± 30° with vertical templates (*N* = 346; Fig. 4f, green; Supplementary Fig. 35) and 67% within 0 ± 30° with horizontal templates (*N* = 326; Fig. 4f, orange; Supplementary Fig. 36). Subpopulations of QRs deviating by around ±30° from the designed orientations (Fig. 4f, arrows) are consistent with capillary-force-driven reconfiguration within the origami topography (Supplementary Fig. 37).

Widefield fluorescence microscopy revealed photoluminescence (PL) from individual QRs in the arrays (Fig. 4g). To demonstrate deterministic emission polarization control, we measured PL intensity versus analyzer angle for two orthogonally oriented QR arrays (Supplementary Fig. 38). Results showed sinusoidal relationships with anti-correlated PL intensities (Fig. 4h), indicating ensemble QR alignment along roughly orthogonal directions. These results show that TMOP can set both the position and the orientation of individual colloidal emitters on a device substrate.

Individual QRs within the arrays showed single-peaked PL spectra with peak wavelengths spanning 578.6 to 636.6 nm and linewidths of 15.3 ± 2.2 nm (*N* = 18), as expected for isolated QRs (Fig. 4i and Supplementary Fig. 39). To assess the quantum optical properties of the placed emitters, we performed Hanbury Brown–Twiss measurements on individual TMOP-directed QRs under pulsed excitation at room temperature (see Methods, Supplementary Note 6, and Supplementary Fig. 40). Second-order correlation results showed pronounced antibunching (Fig. 4j and Supplementary Fig. 41). The *g*(2)(0) values, obtained from background-corrected raw coincidence peak areas (Supplementary Note 6), fell below the 0.5 single-photon threshold for 45 of the 49 randomly selected emitters measured, with a median value of 0.129 (Fig. 4k). TMOP-directed QRs thus retained the room-temperature single-photon emission reported for isolated, randomly deposited QRs^45,48^.

## Deterministic emitter integration with on-chip photonics

Integrating colloidal quantum emitters into practical devices typically requires their deterministic placement on non-uniform topographies^5,6^. We therefore tested TMOP on two photonic platforms: SiN_x_, a widely used PIC material, to assess placement on device topographies, and TiO_2_, whose high refractive index and negligible visible autofluorescence^49^ suit single-emitter optical measurements. We fabricated 310-nm-thick SiN_x_ waveguides, micro-ring resonators, and bullseye cavities on silicon chips (Fig. 5; Supplementary Fig. 42). SiN_x_ structures were first treated with O_2_ plasma to create surface silanol groups. After PMMA coating, wells orthogonal or parallel to the waveguides were patterned on their top surfaces by EBL. Because the resist partially planarizes over the 310-nm-high structures, it is thinner on narrower waveguides, giving well depths of 46 nm, 62 nm and 86 nm on 0.5 µm, 1 µm and 2 µm wide waveguides, respectively (Supplementary Fig. 43). Despite wells deeper than those on flat substrates (≈30 nm), origami placement yield on 2 µm waveguides reached 94% (*N* = 336; Supplementary Fig. 44), and subsequent QR attachment generated deterministically arranged individual QRs with a 71% placement yield (*N* = 329; Supplementary Fig. 45). We also demonstrated TMOP-directed QR placement on SiN_x_ micro-ring structures with potential applications in lasers^50^ and optical sensors^51^ (Fig. 5a; Supplementary Fig. 46). QR orientation can be set parallel or perpendicular to the local tangential direction of the ring (Fig. 5b), which could be used to tune emitter–device

**Fig. 5.**
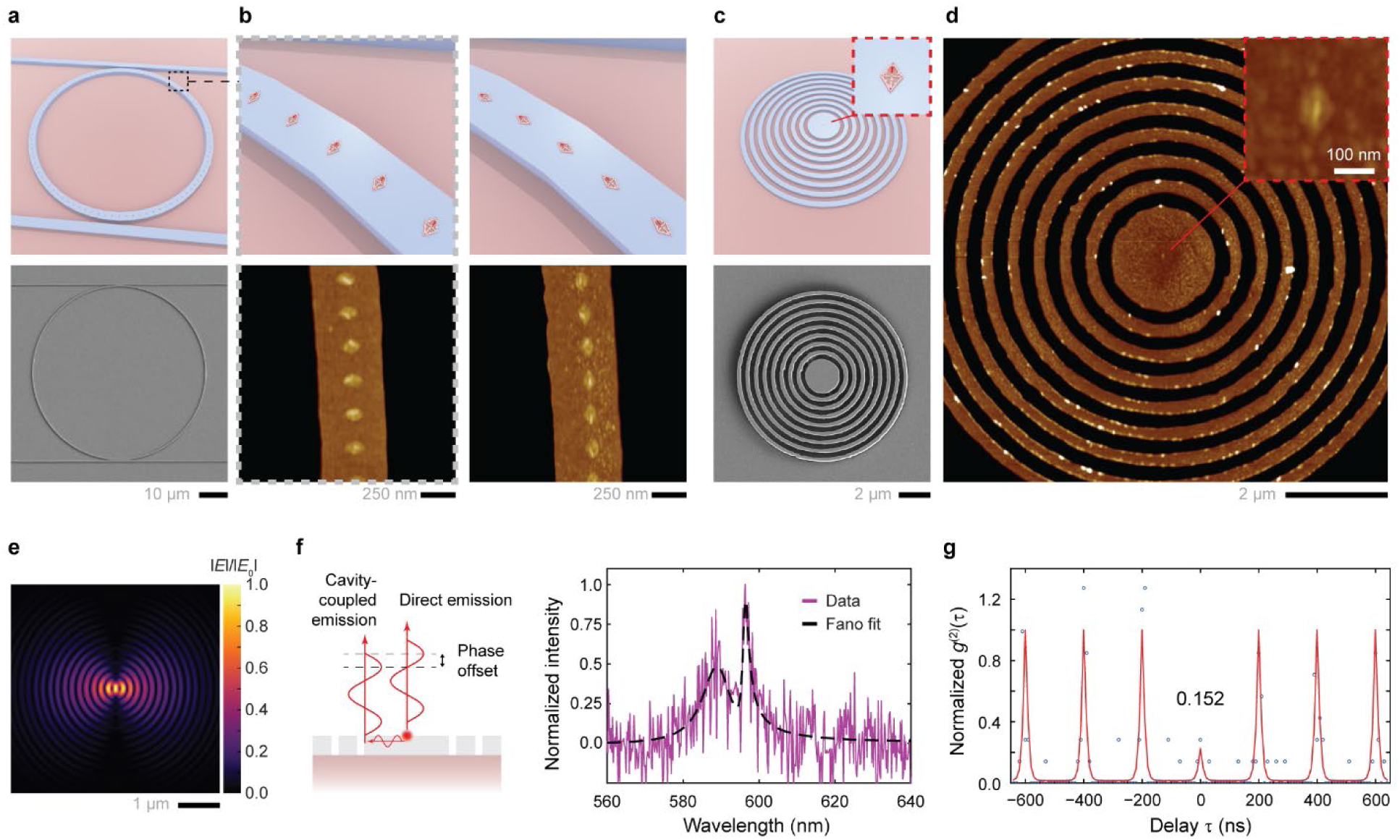
Deterministic QR integration with photonic structures. **a**, Schematic and SEM image of a micro-ring resonator with QRs. **b**, AFM images of orthogonal (left) and parallel (right) QRs on micro-rings. **c**, Schematic of a single QR at a bullseye cavity center and SEM image of a SiNx bullseye (310 nm thick). **d**, AFM images of a single QR at the cavity center. **e**, FDTD-simulated in-plane field distribution of the target TiO2 bullseye cavity (see Methods). **f**, Schematic and PL spectrum (purple) of a single QR at the center of a TiO2 bullseye cavity, showing a Fano line shape (fit, black). **g**, Second-order correlation trace, *g*(2)(*τ*), of the cavity-coupled QR under pulsed excitation. Blue points, data; red line, fit.

coupling given the polarized emission of QRs. Bullseye cavities are promising resonators for quantum optics^52^, but a key challenge is positioning individual emitters at the cavity centers to maximize coupling and avoid emission artifacts^53^. Using TMOP, we demonstrated the deterministic placement of single QRs at the centers of SiN_x_ bullseye cavities with a 68% yield (23 of 34 imaged; Fig. 5c,d; Supplementary Fig. 47).

For device-coupled single-photon emission, we fabricated TiO_2_ bullseye cavities from a sputtered TiO_2_ film on Si/SiO_2_ substrates, sweeping their eye radius, grating period, and duty cycle to tune the resonance into the QR emission band (see Methods; Supplementary Fig. 48). For these devices, the TMOP method was adapted to the sputtered TiO_2_ surface by priming the well bases with the silicification precursor to promote origami binding and pre-loading QRs onto origamis (see Methods). Finite-difference time-domain (FDTD) simulations of the target bullseye geometry predicted a resonance within the QR emission band (Supplementary Fig. 48c) and strong field confinement at the cavity center (Fig. 5e). Test cavities loaded with a bulk spin-coated QR emitter film showed a well-defined resonance with a quality factor of ≈284 (Supplementary Fig. 48d and Supplementary Note 7), confirming that the fabricated devices resonate with QR emission. For single emitters positioned at the field maximum of a bullseye cavity, the collected light comprises both direct emission from the QR and cavity-mediated emission, which can interfere coherently^54^ (Fig. 5f). Because the phase of the cavity-mediated pathway shifts by approximately π across the resonance, the two contributions interfere constructively on one side of the cavity mode and destructively on the other. Since the QR emission is spectrally broad compared with the cavity resonance linewidth, this interference imprints an asymmetric Fano line shape on the spectrum, even at the single-photon level (Supplementary Note 8). The measured PL spectrum showed this asymmetry and was well described by the model with a moderate coupling parameter of 0.51 (Fig. 5f and Supplementary Table 2), consistent with coherent interference between the direct and cavity-mediated emission pathways. Critically, the same device gave *g*(2)(0) = 0.152 (Supplementary Note 6) under pulsed excitation at room temperature, indicating single-photon emission (Fig. 5g). In contrast, single QRs on flat substrates showed symmetric spectra with antibunching (Fig. 4i,j), whereas QR clusters at a cavity center produced asymmetric Fano line shapes without antibunching (Supplementary Fig. 49). TMOP therefore delivered a deterministically positioned single-photon emitter at the field maximum of an on-chip optical cavity.

### Conclusions

The TMOP method bridges bottom-up molecular and colloidal synthesis with top-down fabrication of photonic devices. By combining DNA origami with EBL-defined topographical confinement, we controlled the number and orientation of plasmonic NPs, and our soft-silicification-passivation strategy extended this control to divalent-cation-sensitive colloidal quantum emitters. This enabled deterministic single-QR placement on waveguides, micro-ring resonators and bullseye cavities, and cavity-coupled single-photon emission from a TiO_2_ bullseye cavity. Resist lift-off leaves clean device surfaces, and the bath-based workflow is compatible with standard nanofabrication and wafer-scale processing. The placement and orientation accuracy of colloidal nanomaterials could be further improved through rational design of origami geometry and binding-site positions that better resist capillary-force-driven reconfiguration during drying^15,33,35^. Because DNA origami placement is also compatible with diamond-like carbon^22^, glass^27^, and 2D materials^55,56^, the approach could extend to other photonic platforms, bringing the synthetic tunability of colloidal emitters to integrated quantum photonic circuits for communication, sensing, and computing.

## Methods

### DNA origami design and assembly

6HB wireframe origamis were designed using ATHENA^57^. Design details and DNA sequences are provided in Supplementary Note 1. Origamis were folded in a 12.5 mM MgCl_2_ buffer containing 40 mM Tris (tris(hydroxymethyl)aminomethane) and 12.5 mM MgCl_2_ (pH = 8.0 ± 0.2)^58^. Specifically, 2 nM DNA scaffold (M13mp18/p7249, Tilibit nanosystems) was mixed with 20 equivalent unmodified staple strands and 60 equivalent modified staple strands (standard desalting, Integrated DNA Technologies (IDT)) in 12.5 mM MgCl_2_ buffer. The mixed solution was annealed in a thermocycler (Bio-Rad T100) using the following program: 95 °C for 5 min, 85 °C to 76 °C at 5 min/°C, 75 °C to 30 °C at 13.75 min/0.5 °C, 29 °C to 25 °C at 10 min/°C, then held at room temperature (*ca.* 23 °C). The folded origami sample was used directly for TMOP without further purification.

### DNA functionalization of NPs

All NPs were functionalized with a thiolated single-stranded DNA (ssDNA) purchased from IDT with standard desalting (sequence: /5ThioMC6-D/TTTTTTTTTCCCAGGTTCTCT) as a disulfide-modified strand and further reduced by 100 equivalent tris(2-carboxyethyl)phosphine hydrochloride (TCEP) at a final concentration of 1 mM.

AuNSs were functionalized using a freezing-assisted method^59^. Citrate-stabilized AuNS (17 nm diameter) was synthesized following a standard seeded growth protocol^60^. 1 mL crude AuNS (≈2 nM) was mixed with 6,000 equivalent thiolated DNA and kept in a −80 °C freezer for 15 min, followed by defrosting at room temperature. Thawed AuNS solution was washed six times with 10 mM NaCl buffer (10 mM Tris, 10 mM NaCl, pH = 8.0 ± 0.2) by centrifugation at 16,000 rcf for 3 min and quantified by UV-Vis absorption at 450 nm (extinction coefficient: 3.24 × 10^8^ M^−1^ cm^−1^).

AuNRs were functionalized using a dehydration-assisted method^61^. 2 mL 0.1 nM citrate-stabilized AuNR (nanoComposix, catalog number: GRCN800) was pelleted by centrifugation at 6,000 rcf for 5 min and mixed with 30,000 equivalent thiolated DNA, with a final volume of around 20 µL. 180 µL 1-butanol was added to the mixture, followed by brief vortex mixing. 200 µL 10 mM NaCl buffer was then added followed by another quick vortex mixing and brief centrifugation to facilitate liquid phase separation, with DNA-functionalized AuNRs dispersed in the bottom aqueous phase. The AuNR solution was collected and washed six times with 10 mM NaCl buffer by centrifugation at 4,000 rcf for 4 min and quantified by UV-Vis absorption at 830 nm (extinction coefficient: 5.9 × 10^9^ M^−1^ cm^−1^).

QRs were functionalized using an ultrasonication-dehydration-assisted method^17^. 10 µL 5 mg/mL CdSe/CdS core-shell type QRs in hexane (Sigma-Aldrich, catalog number: 900514) was mixed with 500 equivalent thiolated ssDNA in the presence of 100 mM NaOH for a final volume around 50 µL. The mixture was ultrasonicated at 37 kHz for 5 min, immediately followed by adding 600 µL 1-butanol and brief vortex mixing. 200 µL 10 mM NaCl buffer was then added to the mixture followed by another quick vortex and brief centrifugation to facilitate liquid phase separation, with DNA-functionalized QRs dispersed in the bottom aqueous phase. The QR solution was collected and washed using an ultracentrifugal filter (Amicon 100 kDa) six times (8,000 rcf, 3 min) and quantified by UV-Vis absorption at 350 nm (extinction coefficient: 6.485 × 10^7^ M^−1^ cm^−1^).

### SiN_x_ photonic structure chip fabrication

The fabrication started with a 100 mm Si (100) wafer (p-type boron-doped) with 3 µm thermally grown wet oxide (prime grade, University Wafer Inc.). Upon receipt, the wafer was thoroughly cleaned following the RCA clean protocol: RCA-1 (deionized (DI) water/ammonium hydroxide/hydrogen peroxide, 7:2:2) followed by RCA-2 (DI water/hydrochloric acid/hydrogen peroxide, 7:2:2). For each step, the solution was heated to 75 °C and the wafer was cleaned for 10 min, followed by DI water rinse. After the complete RCA clean, the wafer was dried using a spin rinse dryer. Then, the wafer was loaded in a low-pressure chemical vapor deposition chamber (Tystar Mini Tytan 4600) to deposit the SiN_x_ layer at 770 °C, with a pressure of 250 mTorr, 25 sccm of dichlorosilane and 75 sccm of ammonia, resulting in a growth rate of *ca.* 3.41 nm/min. A thickness of 310 nm was targeted for the SiN_x_ film. After growth, the refractive index and thickness were characterized with reflectometry (Filmetrics F50) and ellipsometry (J.A. Woollam model RC2), followed by singulation of the wafer into individual pieces through cleaving.

For waveguide patterning, chips were first coated with hexamethyldisilazane (HMDS) at 150 °C in a YES-310TA vapor prime oven for 1 min (Yield Engineering Systems Inc.). A thin UV-sensitive negative photoresist (AZ-nLOF 2020, Microchemicals GmbH) was coated using a CEE Apogee spin coater at 2,000 rpm for 60 s, followed by baking at 110 °C for 1 min. Before spinning, the photoresist was first diluted in an equal amount by volume of methyl isobutyl ketone (MIBK) to achieve a lower thickness, more suitable for high lithographic resolution. The chips were subsequently coated with a thin conductive layer (ESpacer 300Z from Resonac) by spin coating at 4,000 rpm for 45 s. Exposure was performed using an ELS-BODEN EBL system (Elionix HS-50), with 10 nA current and 50 kV beam voltage. After exposure, chips were rinsed in water to remove the conductive layer, post-exposure baked at 115 °C for 1 min, developed in a 2.38% tetramethylammonium hydroxide (TMAH)-based developer (AZ 726) for 2 min, and thoroughly rinsed with deionized water. The patterned chips were dry etched using an inductively coupled plasma (ICP) reactive-ion etcher (SAMCO RIE-230iP) at 1 Pa pressure, 300 W ICP power, 100 W bias power, 20 sccm of CF_4_ and 60 sccm of Ar. After etching, the photoresist mask was stripped using an O_2_ plasma (ESI 3511V-001).

### FDTD simulation of TiO_2_ cavities

Finite-difference time-domain simulation software Ansys Lumerical FDTD was used to optimize the bullseye geometry and to evaluate the local field distribution and the cavity resonance. The modeled target structure comprised a 100-nm-thick TiO_2_ bullseye cavity with an eye radius of 570 nm surrounded by seven concentric grooves of 100 nm depth and 343 nm period with a duty cycle of 0.79, on a 1,010 nm (measured) SiO_2_ layer on a Si substrate. The refractive indices of TiO_2_ and SiO_2_ were taken as 2.36 and 1.45 (measured), respectively. Emission was modeled as a broadband in-plane electric dipole at the cavity center, with the structure in its final, lifted-off state. The mesh size was 10 nm in *x*, *y* and 20 nm in *z*.

### TiO_2_ bullseye cavity fabrication

TiO_2_ bullseye cavities were fabricated on Si/SiO_2_ substrates. A TiO_2_ film of 100 nm was deposited on the 1 µm wet thermal oxide layer, by reactive sputtering in an AJA ATC-1800, with a Ti target, 300 W power, 3 mTorr pressure, 27 sccm Ar flow, and 14 sccm O_2_ flow, with a deposition rate of approximately 0.16 Å/s. A negative-tone electron-beam resist, ma-N 2403, was spin-coated at 4,000 rpm, subsequently baked at 90 °C for 2 min, and exposed by EBL in an Elionix HS-50 with 50 kV acceleration voltage, 1 nA beam current, and a dose of 700 µC/cm^2^, then developed in AZ300 MIF for 60 s at room temperature. The patterned resist served as the mask for reactive-ion etching in a SAMCO 230iP tool at 100 °C with 100 W bias, 400 W ICP, 6 sccm CH_4_, 50 sccm CF_4_, 45 sccm Ar for 94 s, after which the mask was stripped with an O_2_-based reactive-ion etching at room temperature with 50 W bias, 100 W ICP, and 50 sccm O_2_ for 300 s. Arrays with eye radius, grating period, and duty cycle varied over 540–600 nm, 293–393 nm, and 0.69– 0.81, respectively, were fabricated to accommodate the natural distribution of single-QR emission wavelengths and fabrication deviations from the simulated optimum for resonance match. The representative design from Supplementary Fig. 48d had 7 rings, 570 nm eye radius, 363 nm period, 0.71 duty cycle, and etch depth of around 120 nm. Cavity dimensions were measured with SEM.

### Fabrication of resist wells on silicon chips

Resist layer wells were fabricated via standard EBL. Silicon chips (1 cm × 1 cm) cleaved from a 100 mm Si (100) wafer (p-type boron-doped) with 90 nm dry thermal oxide (prime grade, University Wafer Inc.) were first cleaned by ultrasonication in isopropanol (IPA) for 10 min and dried with pressurized N_2_, followed by O_2_ plasma treatment at 500 W, 255 duty ratio with 10 sccm O_2_ for 2 min on a Tergeo Plus Plasma Cleaner (PIE Scientific LLC). Additional H_2_O plasma treatment under the same conditions with 10 sccm H_2_O instead of O_2_ can further increase surface hydroxyl group coverage. After priming with HMDS at 150 °C in a YES-310TA vapor prime oven (Yield Engineering Systems Inc.) for 1 min, PMMA resist (950 A2 in anisole, MicroChem) was spin coated at 4,500 rpm for 1 min and baked at 180 °C for 5 min, yielding around 60-nm-thick resist layers. Next, EBL exposure was performed with various doses using an Elionix HS-50 with 2 nA operating current and 50 kV beam voltage. The target well dimensions were 150 nm × 80 nm (long and short axes length) for the rhombic structure, 150 nm × 40 nm (length and width) for the ladder-shaped structure, and 145 nm (edge length) for the equilateral triangular structure. Proximity effect correction was applied for 250 nm spacing wells with the BEAMER software. Patterns were developed with MIBK/IPA 1:3 (MicroChem) at 4 °C for 50 s, rinsed with IPA, dried with pressurized N_2_, and then exposed to directional O_2_ plasma on an RIE-230iPC system (SAMCO) for 18 s at 20 °C, 2.7 Pa with 50 W bias and 20 sccm O_2_. Post-RIE well depth was measured to be around 30 nm. Increasing EBL doses from 1,020 µC/cm^2^ to 1,320 µC/cm^2^ and 1,722 µC/cm^2^ resulted in 97 × 135 nm (size 1), 106 × 150 nm (size 2), and 114 × 161 nm (size 3) as the short and long axes length of the rhombic well openings measured by AFM (*N* = 144 each), respectively. Note that AFM measurements were convoluted with probe size and imaging parameters, and O_2_ plasma etching expanded the well opening, both with potential bias on sharp well vertices.

For typical experiments, identical patterns were fabricated on both halves of each chip, which were then cleaved into two 0.5 cm × 1 cm pieces using a PELCO LatticeAx 420 system (Ted Pella Inc.). For SiN_x_ photonic structure chips (1 cm × 1 cm), the process remained unchanged except that 950 PMMA A4 was employed, followed by spin coating of DisCharge H_2_O (4×, DisChem Inc.) to improve electron conductivity. After aligned EBL exposure with global fiducial markers, the chips were first rinsed with water to remove the DisCharge coating, then developed using the same setup for 90 s.

### Topography-modulated origami placement

Crude DNA origami directly from folding at 2 nM in 12.5 mM MgCl_2_ buffer was adjusted to 1 nM with a final MgCl_2_ concentration of 35 mM and NaCl concentration of 100 mM. 20 µL crude origami solution was pipetted onto a 0.5 cm × 1 cm silicon chip bearing EBL-fabricated well patterns and incubated at 23 °C for 2 h in a humid petri dish with wetted Kimwipes (Kimtech) to reduce evaporation, with 300 rpm shaking on a BioShake iQ shaker. The chip was then transferred into a plate well filled with 0.9 mL 40 mM MgCl_2_ buffer (containing 10 mM Tris and 40 mM MgCl_2_, pH = 8.0 ± 0.2) on a 48-well culture plate (10861-560, VWR), with the patterned side up. The chip was washed by shaking the plate at 700 rpm for 5 min and then transferred into another plate well with 0.9 mL fresh buffer for the next wash. In total, the chip was washed 4 times in 0.9 mL 40 mM MgCl_2_ buffer and 2 times in 0.9 mL 60 mM MgCl_2_ buffer (containing 10 mM Tris and 60 mM MgCl_2_, pH = 9.0 ± 0.2) to remove excess DNA while enhancing substrate binding with higher Mg^2+^ concentration and higher buffer pH. The chip was then dried by quickly dipping into water twice to remove bulk residual buffer and blow-drying with pressurized air. For PMMA lift-off, the dried chip was ultrasonicated in an *N*-methylpyrrolidone (NMP) bath at the lowest intensity setting on the instrument for 6 min using a sweepSONIK (40 kHz dualSWEEP) ultrasonic generator (Blackstone-NEY Ultrasonics, Cleaning Technologies Group, LLC). The chip was then immersed and stirred for 30 s in an acetone bath and IPA bath consecutively and dried with pressurized N_2_.

For TMOP on SiN_x_ photonic structures (1 cm × 1 cm chip), 100 µL origami solution was applied and incubated with the chip for 3 h. The chip was washed in a 24-well culture plate (10861-558, VWR) with 2 mL wash buffers each time to adapt for the larger chip size.

To silicify origami for SEM characterization (not a soft-silicification), a silicification precursor solution was prepared freshly by adding 75 µL (3-aminopropyl)triethoxysilane (APTES, 99%, 919-30-2, Sigma-Aldrich) and 150 µL tetraethyl orthosilicate (TEOS, ≥99% GC, 78-10-4, Sigma-Aldrich) to 4.775 mL 12.5 mM MgCl_2_ buffer sequentially with vortex mixing in between. The mixture was incubated at room temperature with 900 rpm shaking for 30 min and left undisturbed for 20 min, followed by filtration through a 0.22 µm centrifuge tube filter (Corning Costar Spin-X). After TMOP, prior to drying, the origami-placed chip was transferred to a new plate well containing 1 mL as-prepared silicification precursor, and incubated for 1.5 h with 500 rpm shaking. The chip was then washed in 0.9 mL 12.5 mM MgCl_2_ buffer 4 times in the plate wells. Drying and lift-off were the same as described above.

### TMOP-directed gold NP arrangement

DNA origami with binding overhangs was first placed through TMOP as described above until completion of 40 mM MgCl_2_ buffer washes. The chip was removed from the buffer and its back side dried with a Kimwipe, without removing the liquid on the top side of the chip. The chip was placed back into the humid petri dish and 20 µL 10 nM DNA-functionalized AuNSs (or 1 nM DNA-functionalized AuNRs) in 40 mM MgCl_2_ buffer was added onto the chip. After pipetting the liquid in and out 10 times for mixing, the chip was incubated at room temperature for 4 h (2.5 h for AuNRs). The chip was then washed 4 times in 0.9 mL 40 mM MgCl_2_ buffer and 2 times in 0.9 mL 60 mM MgCl_2_ buffer in the plate, followed by drying and lift-off as described above.

### TMOP-directed QR arrangement

DNA origami with binding overhangs was first placed through TMOP and then soft-silicified to accommodate Mg^2+^-free conditions. Soft-silicification precursor solution was prepared freshly by sequentially adding 150 µL N-trimethoxysilylpropyl-N,N,N-trimethylammonium chloride (TMAPS, 50% in methanol, 35141-36-7, Fisher Scientific) and 150 µL TEOS to 4.7 mL 12.5 mM MgCl_2_ buffer with vortex mixing in between. This mixture was incubated at room temperature with 900 rpm shaking for 30 min and left undisturbed for 20 min to allow large silica particles to float to the top of the solution. 1 mL solution was taken from the bottom of the mixture and added to a plate well on the 48-well plate. After origami placement in TMOP, the chip was washed 3 times in 0.9 mL 40 mM MgCl_2_ buffer and once in 0.9 mL 60 mM MgCl_2_ buffer in the plate, before transferring to the plate well containing the soft-silicification precursor and incubating for 1.5 h. The chip was then washed 4 times with 0.9 mL 12.5 mM MgCl_2_ buffer and transferred to another plate well containing 1 mL passivation solution, which was freshly prepared by mixing 200 µL carboxyethylsilanetriol disodium salt (CTES, 25% in water, 18191-40-7, Oakwood) with 4.8 mL 12.5 mM MgCl_2_ buffer and 30 µL concentrated hydrochloric acid (37%, 7647-01-0, Sigma-Aldrich) and filtering through an Amicon Ultra Centrifugal Filter (10 kDa, Sigma-Aldrich). After 15 min incubation, the chip was washed 2 times in 0.9 mL 12.5 mM MgCl_2_ buffer and 3 times in 0.9 mL phosphate-buffered saline (PBS) buffer (without calcium and magnesium, pH = 7.4 ± 0.1, 21-040, Corning).

The chip was removed from the buffer solution and its back side dried with a Kimwipe, without removing the liquid on the top side of the chip. The chip was then placed back into the humid petri dish and 20 µL 20 nM DNA-functionalized QRs in PBS buffer was added onto the chip. After pipetting the liquid in and out 10 times for mixing, the chip was incubated at room temperature for 4 h. The chip was then washed 4 times in 0.9 mL PBS buffer and 2 times in 0.9 mL 60 mM MgCl_2_ buffer in the plate, followed by drying and lift-off as described above. Alternatively, chips were dried through an ethanol series (dipping in 50% ethanol for 20 s and in 75% ethanol for 40 s) followed by a gentle air blow, which improved on-chip QR PL.

For QR placement on SiN_x_ and TiO_2_ photonic structure chips (1 cm × 1 cm), all steps were performed in a 24-well culture plate (10861-558, VWR) with 2 mL corresponding solutions to adjust for the larger chip size. 100 µL QR solution was used for QR binding.

### Energy-landscape model and kinetic simulation

Two-dimensional projected DNA structures and corresponding deposition shapes were modeled with their nanoscale dimensions: 125 nm × 75 nm (rhombic structure), 140 nm × 40 nm (ladder-shaped structure), and 140 nm × 120 nm (triangular structure). Thermodynamic energy landscapes were calculated by establishing a conformational grid on *X*, *Y* coordinates and angle *θ* (structure rotation relative to deposition shape). Using each deposition location as the origin, grid points were defined as: −80 nm ≤ *X, Y* ≤ 80 nm with 5 nm intervals, and −90° ≤ *θ* ≤ 87° for rhombic and ladder-shaped structures and −60° ≤ *θ* ≤ 57° for triangular structure with 3° intervals. For each grid point, we calculated the overlapped area (*A*_o_) between a structure and its deposition shape by generating alpha shapes and summing triangulated overlapped regions. The binding free energy for both no-well and well cases was determined as *E*_b_ = *u*_b_*A*_o_, where *u*_b_ is the assumed binding free energy per area (−0.001, arbitrary units). The free energy penalty, calculated only for the well case, was set as *E*_r_ = *u*_r_*A*_r_, where *u*_r_ is the assumed free energy penalty per area (0.001*α*, where *α* is an adjustable penalty coefficient, see Supplementary Note 4; *α* = 5 for Fig. 2e) and *A*_r_ represents area outside the well. Total interaction free energy was computed as *E*_t_ = *E*_b_ + *E*_r_.

Kinetic simulations were deterministic steepest-descent searches on this energy landscape. Each grid point served as an initial conformation with DNA structures allowed to move toward energetically favorable conformations. The perturbing intervals for searching neighboring conformations were set to ±5 nm for *X* and *Y* positions, and ±3° for rotation angle *θ*. From each initial state, structures were translated or rotated to the neighboring conformation with the lowest *E*_t_ of its 6 neighbors until reaching a local minimum. For TMOP simulations, initial conformations with a positive *E*_t_ were eliminated due to the lack of origami interaction with the well base (see Supplementary Notes 3 and 4). *P* was the fraction of initial conformations reaching each final conformation.

### AFM and SEM characterization

AFM measurements were performed in ambient (air) using an Asylum Research Jupiter XR AFM (Oxford Instruments) in tapping mode with 55AC-NG gold-coated ultra-high frequency AFM probes (tip radius < 7 nm, *f*_0_ = 1,200 kHz; 85 N/m, NanoAndMore). SEM imaging was conducted on a Zeiss Sigma HD VP Scanning Electron Microscope at 2 kV, using an Inlens Secondary Electron Detector for silicified TMOP origami arrays (without coating) and an Everhart–Thornley Secondary Electron Detector for SiN_x_ photonic structures (after 5-second gold coating with a PELCO SC-7 Sputter Coater).

### Optical microscopy

Dark-field microscopy of TMOP-directed AuNR arrays was conducted using a Nikon Eclipse LV150NA microscope equipped with a DS-Ri2 camera and 100× air objective (Nikon MUE61900 EPI, NA 0.8), employing a rotatable polarizer for incident light and an analyzer with vertical polarization.

Confocal fluorescence imaging of QR arrays was conducted on a home-built setup using a Nikon Eclipse Ti microscope body with a piezo-controlled 3D movable stage and a 100× air objective (Nikon MUE42900, NA 0.9). Arrays were raster-scanned using a piezoelectric stage under 488 nm laser excitation with circularly polarized light achieved using a wire grid polarizer (Thorlabs WP25M-VIS) and a quarter-wave plate (Thorlabs AQWP05M-600). Emission was filtered through a 506 nm single-edge dichroic (Semrock FF506-Di03) and 500 nm long-pass filter (Thorlabs FELH0500), followed by a rotatable wire grid polarizer (Thorlabs WP25M-VIS) rotated in 20° increments between scans. The filtered emission was directed through a telescopic 1:1 pinhole system (20 cm lenses, 50 µm pinhole) and collected on a silicon avalanche photodiode. Widefield PL imaging was performed on a Nikon Ti-E inverted microscope equipped with a 100× air objective (Nikon CFI TU Plan ELWD, MUE21900; NA 0.80). Samples were excited by a white-light LED (CoolLED pE-300lite) spectrally filtered by a bandpass filter (450 nm center wavelength, 40 nm full width at half maximum (FWHM); Thorlabs FBH450-40). Excitation and emission were separated by a 505 nm long-pass dichroic mirror (Thorlabs DMLP505R), and PL was further filtered by a 500 nm long-pass filter (Thorlabs FELH0500) before detection on an electron-multiplying charge-coupled device (EMCCD) camera (Andor iXon Ultra 897).

### Photon correlation and spectrum measurements

Second-order photon correlation and PL spectrum measurements of individual TMOP-placed QRs on flat substrates were performed at room temperature on a home-built confocal micro-PL setup. The sample was mounted on a piezo stage (Physik Instrumente) on top of a translation stage and excited through a 100× objective (NA 0.85) by a pulsed 532 nm laser at a repetition rate of 5 MHz with an average power of 1 µW. Excitation and emission were separated with a dichroic filter, and the collected emission was routed by a flip mirror either to a spectrometer for PL spectra or to a Hanbury Brown–Twiss detection arm, in which photons were split by a 50:50 beamsplitter onto two avalanche photodiodes (Micro Photon Devices, PD-050-CTE) and their arrival times registered by a time-correlated single-photon counting (TCSPC) module (PicoQuant, MultiHarp). PL spectra were collected with a spectrometer (Princeton Instruments, SP-2300) using a 300 grooves/mm grating and a thermoelectrically cooled charge-coupled device (CCD) camera (Princeton Instruments, PIXIS 100). Coincidence histograms were constructed and *g*(2)(0) calculated from raw peak areas (not fitted) as described in Supplementary Note 6.

### Cavity-coupled PL on TiO_2_ bullseyes

TiO_2_ cavity resonances and cavity-coupled QR emission were characterized on the confocal micro-PL setup described above. To test cavity resonances, a 20 µL solution of 82 nM DNA-functionalized QRs was spin-coated onto the TiO_2_ bullseye chip at 3,000 rpm for 1 min. For TMOP on TiO_2_ bullseye cavities, chips were treated with O_2_ plasma at 150 W, 255 duty ratio with 5 sccm O_2_ for 5 min on a Tergeo Plus Plasma Cleaner, then primed with HMDS at 150 °C for 1 min. PMMA resist (950 A2 in anisole) was spin coated at 2,500 rpm for 1 min and baked at 160 °C for 5 min, followed by spin coating of DisCharge H_2_O (4×, DisChem Inc.) at 2,500 rpm for 1 min without baking. EBL exposure was performed as described above. The chips were rinsed with water to remove the DisCharge coating, developed in MIBK/IPA 1:3 at 4 °C for 60 s with gentle stirring, rinsed with IPA and dried with pressurized N_2_, then exposed to directional O_2_ plasma on an RIE-230iPC system (SAMCO) for 14 s at 20 °C, 2.7 Pa with 50 W bias and 20 sccm O_2_. Because DNA origami binds sputtered TiO_2_ weakly, the well base surface was modified before origami placement by incubating the empty patterned chip in the soft-silicification precursor described above for 30 min. QRs were pre-loaded onto origami in solution by mixing purified rhombic origami (2 nM final concentration) with DNA-functionalized QRs at a 6:1 origami:QR ratio in 1 mM MgCl_2_ buffer (containing 10 mM Tris and 1 mM MgCl_2_, pH = 8.0 ± 0.2) in a total volume of 100 µL, followed by incubation at room temperature overnight. The silicified chip was washed four times in 2 mL 1 mM MgCl_2_ buffer (5 min, 500 rpm), after which the 100 µL origami-QR solution was applied to the chip in a humid petri dish, and incubated at room temperature for 3 h with 350 rpm shaking. The chip was then washed three times in 2 mL 1 mM MgCl_2_ buffer and twice in 2 mL 60 mM MgCl_2_ buffer, 5 min each at 500 rpm. The chip was dried through the ethanol series described above, and kept under vacuum overnight. For lift-off, the chip was immersed in NMP at 70 °C for 3 h, without ultrasonication, then stirred consecutively in acetone and IPA baths and dried with pressurized N_2_. Fitting functions for the cavity resonances and for the cavity-coupled single-QR emission are given in Supplementary Notes 7 and 8, respectively.

## Data processing

Detailed photon-correlation analysis is described in Supplementary Note 6, and cavity-coupled emission analysis in Supplementary Notes 7 and 8. Detailed image processing methods to acquire sizes, yields, distributions, and placement precisions are described in Supplementary Note 9. Measured values are reported as mean ± s.d. unless stated otherwise. For placement precision, the uncertainty of each standard deviation is the standard error across AFM images (N = 4, 5 and 4 images for well sizes 1, 2 and 3).

## Large language model disclosure

The large language models, Claude Opus 5 and 5.5 (Anthropic), were used to edit and condense text to improve readability. The Python code used to quantify TMOP precision and analyze photon correlation was developed with the assistance of Claude Opus 5 (Anthropic). The authors defined the analysis, guided code development, and verified the code and its outputs, both of which are available in the Dryad repository^62^.

### Data availability

All data supporting the findings of this study are available within the paper and its Supplementary Information. DNA origami designs and staple sequences are provided in the Supplementary Information and in the Dryad repository^62^. Raw data and additional images are openly available in the Dryad repository^62^.

### Code availability

The image processing code, photon-correlation analysis code and simulation code used in this study are openly available in the Dryad repository^62^.

## Acknowledgements

M.B., X.L. and C.C. acknowledge funding from the U.S. National Science Foundation (CCF-1956054) and the U.S. Office of Naval Research (N000142114013). M.B., X.L. and P.F.S. acknowledge funding from the U.S. National Science Foundation (FET-2403100) and the U.S. Office of Naval Research (N000142512217). M.B., R.J.M., X.L., P.F.S., F.N., S.B.D. and N.N. acknowledge funding from the U.S. National Science Foundation Future Manufacturing Seed Grant program (FMSG-2428308). J.H. and L.R. acknowledge funding from the U.S. National Science Foundation ITE Convergence Accelerator Track I (ITE-2236093). R.J.M., X.L. and N.S. acknowledge funding from the U.S. Air Force Office of Scientific Research Natural Materials and Systems Program (FA9550-23-1-0210 and FA9550-25-1-0202). M.G.B., T.S. and A.J. acknowledge funding from the U.S. Department of Energy (DE-SC0021650). T.S. thanks the Natural Sciences and Engineering Research Council of Canada for a Postgraduate Scholarship. J.Y.L. thanks the National Research Foundation of Korea for a Sejong Science Fellowship (NRF-2021R1C1C2003554). S.B.D. acknowledges support from the U.S. National Science Foundation Graduate Research Fellowship Program (2141064). N.N. is supported by a National Science Scholarship from the Agency for Science, Technology and Research, Singapore. This work was performed in part through the use of the facilities of MIT.nano and the Harvard University Center for Nanoscale Systems (CNS).

## Author contributions

X.L., F.N., R.J.M. and M.B. conceived the research. X.L. developed the TMOP method and the integration of NPs and QRs on chips, and performed DNA origami design and assembly, materials synthesis, resist well fabrication, nanomaterial placement, dark-field microscopy and AFM characterization. J.H. and L.R. designed and fabricated chips with SiN_x_ photonic structures. M.G.B., T.S. and A.J. designed and conducted the fluorescence and emission polarization experiments. F.N. and S.B.D. designed the TiO_2_ cavity and quantum optical work. S.B.D. simulated and fabricated the TiO_2_ bullseye cavities and performed optical and single-photon characterization. N.N. continued the optical characterization and acquired the cavity-coupled emission data. J.Y.L. developed the computational code and performed the kinetic simulations. P.F.S. contributed to TMOP optimization and analyzed the cavity-coupled emission data. N.S. conducted SEM characterization. C.C. optimized the QR–DNA functionalization. X.L., S.B.D., L.R., T.S., N.N., J.Y.L., P.F.S. and N.S. curated the data, and X.L., S.B.D. and P.F.S. prepared the figures. M.G.B., J.H., F.N., R.J.M. and M.B. supervised the work and acquired funding, and F.N., R.J.M. and M.B. administered the project. X.L. wrote the original draft, and all authors reviewed and edited the manuscript.

## Competing interests

M.B., R.J.M., C.C. and X.L. are co-inventors on a patent application (WO2024186458A2) involving the TMOP method. The patent application (PCT/US2024/016144) was filed by Massachusetts Institute of Technology on Feb. 16, 2024. The authors declare no other competing interests.

## Additional information

Supplementary Information is available for this paper. Correspondence and requests for materials should be addressed to Farnaz Niroui, Robert J. Macfarlane or Mark Bathe.

## Notes

### Summary of Updates

Method renamed from Cavity-Shape Modulated Origami Placement (CSMOP) to Topography-Modulated Origami Placement (TMOP) to avoid confusion between the lithographic resist wells and the optical cavities now used in this work. New results: cavity-coupled single-photon emission from a single TMOP-placed quantum rod in a TiO2 bullseye cavity; room-temperature single-photon emission from quantum rods across arrays. Main text, figures, Methods and Supplementary Information updated accordingly. Authors added: Shelly Ben-David, Natalie Ngoh, Peter F. Satterthwaite and Farnaz Niroui.

