## Supplementary Information for "DNA-directed nanometer-scale integration of single colloidal quantum emitters with on-chip photonics"

#### Contents

Supplementary Notes 1–9
Supplementary Figs. 1–49
Supplementary Tables 1 and 2
Supplementary References

### Supplementary Notes

#### Supplementary Note 1. Design of DNA structures and DNA sequences

Two-dimensional 6HB wireframe origami structures with defined target geometries were designed using ATHENA[1] with M13mp18 (7,249 nt, Tilibit nanosystems) as the scaffold. Sequence routing designs, PDB files and oxView files of the origami structures were included in the online data repository. Models and AFM images of the 6HB rhombic origami and the ladder-shaped origami are shown in Supplementary Fig. 2. The design and sequences of the single-layer triangular DNA origami were adopted from previous research by Gopinath et al.[2]. A full list of the staple strand sequences of all DNA origamis is available in the online data repository.

Suitable staple strands in the 6HB wireframe origami structures were identified using oxView[3,4] for extension modifications at specific locations as binding overhangs or face-selecting overhangs. The selected staple strands have 3' ends oriented toward the same side of the 2D origami structure where sequence extensions are added. The rhombic origami incorporates 36 modified staples (31 face-selecting overhangs and 5 binding overhangs), while the ladder-shaped origami contains 37 modified staples (34 or 31 face-selecting overhangs and 3 or 6 binding overhangs for single or double binding site designs, respectively). Each face-selecting overhang consists of 20 thymidines (20T), while binding overhangs comprise 16 thymidines (16T) followed by the complementary sequence of the binding sequence on the thiolated DNA strand used to functionalize nanoparticles. The sequences of staple strands modified with binding overhangs are detailed in Supplementary Table 1.

#### Supplementary Note 2. TMOP setup and workflow

The TMOP methodology introduces significant operational improvements over existing DNA origami placement techniques, establishing a robust framework for integration with standard fabrication protocols. As shown in Supplementary Fig. 13a, the existing DOP workflow[5] requires three separate washing steps each demanding 5–8 repetitions, where in each repetition a small volume of buffer (20 to 60 μL) is applied to a droplet on the chip using a micropipette. The buffer droplet is then pipetted in and out of the micropipette tip a few times to wash the chip surface, during which time special caution is needed to not touch the chip patterns with the micropipette tip. This manual process presents inherent challenges, particularly when Tween 20 surfactant induces increased droplet spreading, making precise liquid handling more demanding. We observed notable inconsistencies when performing DOP of the rhombic origami, likely stemming from these manual intervention requirements.

In contrast, the TMOP workflow eliminates these operational complexities through strategic innovation in the washing protocol (Supplementary Fig. 13b). The well-patterned PMMA resist layer eliminates the need for surfactant washing, enabling excess origami removal through straightforward buffer bath incubation. In the current practice, chips are washed in standard 24-well or 48-well plates containing wash buffer, with plate shaking for 5-min intervals followed by sequentially transferring the chip through fresh buffer-filled plate wells. This streamlined approach requires only 4–6 standardized wash cycles to remove excess materials in solution, reducing procedural variability. The resist well architecture also enables simple drying using pressurized air flow, eliminating the need for ethanol dehydration sequences. We found that the DNA origami incubation (binding) step of TMOP was also compatible with buffer bath incubation in a well plate, which could further streamline the workflow. However, to reduce the cost of materials (DNA, gold NPs, QRs), most experiments in this research were performed by incubating a droplet of colloidal materials on top of the patterns on the chip in a humid petri dish (smaller volume needed).

The TMOP workflow could in principle be scaled to whole-wafer processing, in which DNA origami and subsequent colloidal nanomaterials would be placed onto many devices on the wafer simultaneously, through incubation and washes in larger buffer baths. The capability to operate with crude DNA origami from folding (with excess staple strands) further suggests that TMOP could be integrated into standard foundry processes to fabricate colloidal-material enabled hybrid nanophotonic devices.

#### Supplementary Note 3. Energy-landscape model and kinetic simulation

To further elucidate the role of shape-matched well confinement, we performed thermodynamic calculations and kinetic simulations to gain insight into the placement process via quantitative modeling. The rhombic origami placement was modeled both with and without resist wells, assuming a positive linear relationship between the binding free energy and the overlap in binding area between the origami and the target shape (landing pad or well) (Supplementary Fig. 16). We calculated the binding free energy of an ensemble of origami-target conformations at distinct translational displacements (d$X$ and d$Y$) between −80 nm and +80 nm using 5 nm increments and rotational displacements ($\theta$) between −90° and +87° in 3° increments, accounting for the rhombic shape's two-fold rotational symmetry (Supplementary Fig. 17). Without wells, an origami in solution initially diffuses to the flat surface and binds to a silica landing pad with a randomly positioned and oriented conformation, followed by largely 2D diffusion to maximize its electrostatic surface-binding free energy[5]. After applying a steepest-descent energy minimization mechanism to the set of initial bound conformations, kinetic simulations predicted numerous metastable equilibria or "trapped" end-states away from the correctly positioned and oriented conformation (d$X$ = d$Y$ = 0, $\theta$ = 0). Consequently, the probability of correctly placed origami was only ≈0.46. In simulating TMOP, we introduced a free energy penalty associated with bound origami in areas outside of the target shape in order to account for the well confinement effect, which was assumed to require a mechanical free energy associated with origami deformation in order to fit in the well-confined space while maintaining a specific non-matching conformation (d$X$, d$Y$, $\theta$) in the ensemble (Supplementary Fig. 16c). Conformations with a positive total interaction free energy (binding free energy and free energy penalty combined) were eliminated from the kinetic simulation due to their lack of sufficient binding interaction of origami with the corresponding well base to retain them there (Supplementary Fig. 16d). Consequently, TMOP simulations demonstrated a single aligned origami conformation ($P$ = 1.00), owing to the elimination of local free energy minima through topographic well confinement. This is consistent with the high placement yield of TMOP observed experimentally (97%). When a more flexible origami structure was employed, which corresponded to a lower free energy penalty for origami deformation, remaining local free energy minima could still lead to decreased placement yield (Supplementary Fig. 18 and Supplementary Note 4). The free energy penalty correlated with the mechanical rigidity and deformation response of the origami placed, highlighting the importance of employing mechanically stiff 6HB origami designs. Simulations of the ladder-shaped 6HB wireframe origami (Supplementary Fig. 19) and the triangular single-layer origami (Supplementary Fig. 20) were also qualitatively consistent with simulation results of the rhombic origami.

Specifically, the kinetic simulation used a three-dimensional grid to analyze DNA origami placement conformation, with spatial coordinates (−80 nm ≤ d$X$, d$Y$ ≤ 80 nm, 5 nm intervals) and rotation angles (−90° ≤ $\theta$ ≤ 87° for rhombus/ladder structures, −60° ≤ $\theta$ ≤ 57° for triangles, 3° intervals). For each grid point (d$X$, d$Y$, $\theta$) in the conformation ensemble, the binding free energy was calculated as $E_b = u_b A_o$, where $u_b$ is the binding free energy per area (−0.001) and $A_o$ is the overlapped area between the wireframe origami structure and the deposition shape (Supplementary Fig. 16a). For well simulations, an additional free energy penalty $E_r = u_r A_r$ was included, where $u_r$ is 0.001$\alpha$ ($\alpha$ being the free energy penalty coefficient) and $A_r$ is the non-overlapping area. The total interaction free energy was $E_t = E_b + E_r$ (Supplementary Fig. 16b). In the kinetic simulation, initial structures were allowed to move toward energetically favorable conformations with ±5 nm positional and ±3° rotational perturbations, continuing until reaching a local free energy minimum.

In well simulations, conformations with positive total interaction free energy ($E_t$) were excluded from the placement conformation ensemble due to the lack of origami interaction with the well base. Specifically, if a certain conformation (d$X$, d$Y$, $\theta$) presents a negative total interaction free energy despite the calculated free energy penalty ($E_r$), the origami can potentially bind the base of the well through some degree of origami deformation (Supplementary Fig. 16c). However, if the calculated total interaction free energy is positive (when $A_o < \alpha A_r$), the origami cannot overcome the energy barrier to interact with the well base through electrostatic interaction at this specific conformation (d$X$, d$Y$, $\theta$) (Supplementary Fig. 16d). Hence, such conformations (d$X$, d$Y$, $\theta$) were removed from the placement conformation ensemble for the subsequent kinetic simulations.

**Supplementary Note 4. Simulation implication on origami rigidity and shape**

The validity of a certain conformation in the well simulation (total interaction free energy $E_t > 0$ or $E_t < 0$) depends on the overlapping areas ($A_o$ and $A_r$) and the free energy penalty coefficient ($\alpha$). When $A_o > \alpha A_r$ and $E_t <$

0, the model indicated that origami with this specific conformation (d$X$, d$Y$, $\theta$) can overcome the free energy penalty to bind to the well base. In real scenarios, well sidewalls are solid physical barriers and the free energy penalty can be associated with the free energy cost to deform the origami structure to fit into the well with the specific conformation (d$X$, d$Y$, $\theta$). Hence, the free energy penalty coefficient ($\alpha$) correlates to the free energy cost of origami structure deformation, which is a function of origami rigidity, structural defects and mechanical response. In general, greater $\alpha$ (higher free energy penalty) corresponds to more rigid and robust origami structures like the 6HB wireframe structures employed in this study. Consequently, more conformations will present a positive total interaction free energy and be removed from the kinetic simulation, among which are some local free energy minimum conformations (Supplementary Fig. 17b). With more flexible origami structures or origami structures with defects (smaller $\alpha$, lower free energy penalty), conformations with local free energy minima can remain in the free energy landscape (Supplementary Fig. 18) and reduce the yield of aligned origami placement.

We also observed strong dependency of the kinetic simulation results on origami geometry. Triangular origami with three-fold rotational symmetry presented drastically fewer local free energy minimum conformations compared to the rhombic origami and the ladder-shaped origami (Supplementary Figs. 19 and 20). Kinetic simulation predicted 99.4% aligned triangular origami placement without a well ($\alpha = 0$), while rhombic origami and ladder-shaped origami (two-fold rotation symmetry) were predicted with 46.4% and 66.3% aligned placement, respectively. This result suggests that origami shapes with higher symmetry exhibit fewer local free energy minima, consistent with high placement yields achieved in literature using the equilateral triangular origami design[2,6].

Moreover, origami with the same geometrical symmetry also behaved differently. Compared to the rhombic origami, the ladder-shaped origami presented more minimally populated trapped local minimum states when simulated without the well confinement effect ($\alpha = 0$) (Supplementary Fig. 19a), while no local minimum state was predicted when the well confinement effect was implemented, even with a less rigid DNA origami structure (smaller free energy penalty) ($\alpha = 1$) (Supplementary Fig. 19b). In the case of the rhombic origami, with $\alpha = 1$, simulation predicted multiple trapped local minimum states similar to when no well confinement effect was considered (Supplementary Fig. 18a).

**Supplementary Note 5. Capillary-force-induced NP reconfiguration in the well**

The positioning of nanoparticles in well structures is significantly influenced by capillary forces during the drying process. As illustrated in Supplementary Fig. 29, when water quickly evaporates from the PMMA well, the receding liquid meniscus can create asymmetric capillary forces that drag nanoparticles at the water–air interface toward the well sidewalls. This effect causes particles to deviate from their intended central positions, with their final location and orientation determined by both well geometry and the random dynamics of the drying meniscus. The impact is particularly evident for AuNRs, with most particles observed against well sidewalls post-drying. In the case of QRs, their height (5 ± 0.6 nm) being comparable to the 6HB origami structure (approximately 5.5 nm in solution) creates additional complexity, as capillary forces can pull them partially or entirely into the triangular holes of the wireframe origami structure, resulting in clusters of orientation deviations of ±30° from their designed alignment, as confirmed by AFM imaging (Supplementary Fig. 37). In future work, capillary forces can be either mitigated or strategically harnessed to control NP alignment. For example, sequential solvent exchange using liquids of decreasing surface tension can reduce meniscus forces. Conversely, capillary forces can be advantageously exploited through rational design choices. Asymmetric well geometries with strategic pinning points can guide meniscus retreat in desired directions, promoting specific particle orientations[7,8], while patterned surface chemistry can create energy gradients that direct particle alignment[9].

**Supplementary Note 6. Second-order photon correlation analysis**

Photon-correlation measurements were performed in a Hanbury Brown–Twiss configuration, in which the collected emission was split by a 50:50 beamsplitter onto the two avalanche photodiodes described in Methods. Detector outputs were recorded by a PicoQuant MultiHarp in T3 time-tagged mode, which stores each detected photon as a sync-pulse index together with a micro-time relative to that pulse at 40 ps resolution. Excitation was pulsed at 5.0007 MHz, corresponding to a repetition period $T_{rep}$ = 199.97 ns. Acquisition times were typically 10–110 s per emitter.

For each emitter the full pairwise cross-correlation between the two detectors was computed over a delay range of ±650 ns, corresponding to three side peaks on either side of zero delay, in 1 ns bins on a grid centered on $\tau = 0$. The delay of a pair is obtained as $\Delta\tau = k \cdot T_{rep} + \Delta t_{micro}$, where $k$ is the difference in sync-pulse index and $\Delta t_{micro}$ the difference in micro-time. A constant electronic delay between the two detector channels was removed by aligning their prompt micro-time peaks; the required offset was below 0.5 ns for all emitters.

All $g^{(2)}(0)$ values were calculated as ratios of peak areas using background-corrected raw coincidence data with 1 ns bin size. Each peak was integrated over a window of ±50 ns about its center, defined as a fixed number of bins so that the central and side peaks receive identical integration ranges. The flat coincidence level $a$ arising from uncorrelated events was measured in the region 60–95 ns from any peak center and subtracted from every peak (background correction):

$$g^{(2)}(0) = \frac{N_0 - a \times n}{N_{side} - a \times n}, \tag{1}$$

where $N_0$ is the central-peak area, $N_{side}$ the mean area of the six side peaks (three on either side of zero delay), and $n$ the number of bins per integration window.

The $g^{(2)}(\tau)$ data were fitted with the two-sided exponential model

$$g^{(2)}(\tau) = BL + A_0 e^{-\frac{|\tau|}{w}} + A_s \sum_{k \neq 0} e^{-\frac{|\tau - kT_{rep}|}{w}}, \tag{2}$$

where $BL$ is a constant baseline, $A_0$ and $A_s$ are the central- and side-peak amplitudes, and $w$ is the peak width, taken to be common to all peaks. The fit was performed by maximum likelihood assuming Poisson-distributed bin counts, with the peak spacing fixed at the excitation repetition period $T_{rep}$ and a single amplitude $A_s$ shared by all side peaks. The resulting curve (solid line) is shown overlaid on the raw coincidence data and is used solely to normalize the display; the reported value of $g^{(2)}(0)$ was obtained independently from the raw peak areas using equation (1) as described above.

The statistical uncertainty of $g^{(2)}(0)$ was estimated as a 68% credible interval by propagating the Poisson counting noise of the central peak, side peaks and accidental window by Monte Carlo sampling, and the raw data, analysis code and per-emitter results are deposited in the Dryad repository[10].

**Supplementary Note 7. Cavity-coupled bulk QR emission spectrum analysis**

Spectra were integrated over the CCD rows spanning the emission spot, background-corrected using signal-free rows of the same frame, despiked with a Hampel filter, and normalized to the peak. Because the cavity center was covered with bulk QR clusters, the collected signal contains both cavity-coupled emission and uncoupled ensemble emission, so the resonance was fit to a Lorentzian on a local linear background,

$$I(\lambda) = \frac{A}{\left[1 + \frac{4(\lambda - \lambda_0)^2}{\Delta\lambda^2}\right]} + c_0 + c_1(\lambda - \lambda_0), \tag{3}$$

where $A$ is the peak amplitude, $\lambda_0$ is the resonance wavelength and $\Delta\lambda$ its full width at half maximum (FWHM). The quality factor follows from the fit as

$$Q \approx \lambda_0/\Delta\lambda. \tag{4}$$

Fits used unweighted nonlinear least squares over a ±6 nm window, approximately 2.5 times the linewidth, which is wide enough that the half-maximum points and the wings constrain $\Delta\lambda$ and the background independently. For the bullseye device measured in Supplementary Fig. 48d, this gives $\lambda_0$ = 608.8 nm and $\Delta\lambda$ = 2.14 nm, hence $Q$ = 284.

**Supplementary Note 8. Cavity-coupled single QR emission spectrum analysis**

Our device consists of a broad dipole emitter coupled to a lossy cavity, which forms an open quantum system. We model this system based on the approach reported previously[11,12], with the introduction of a finite bandwidth to the dipole emitter. Such a system is described by a Jaynes–Cummings Hamiltonian

$$H = \hbar\omega_c \hat{a}^\dagger\hat{a} + \hbar\omega_d\hat{\sigma}_+\hat{\sigma}_- + i\hbar g(\hat{a}^\dagger\hat{\sigma}_- - \hat{a}\hat{\sigma}_+), \tag{5}$$

where $\hbar$ is the reduced Planck constant, $\omega_{d,c}$ are the resonance frequencies of the dipole and cavity, respectively, $\hat{\sigma}_+, \hat{\sigma}_-, \hat{a}$ are the respective lowering/annihilation operators and $g$ is the coupling coefficient. The corresponding electric field at position $\boldsymbol{r}$ and time $t$ is given by[12]

$$\hat{\mathbf{E}}(\boldsymbol{r},t) = \boldsymbol{\epsilon}_d(\boldsymbol{r})e^{-i\phi_d}\sqrt{\gamma_s}\hat{\sigma}_-(t) + \boldsymbol{\epsilon}_c(\boldsymbol{r})e^{-i\phi_c}\sqrt{2\kappa}\hat{a}(t) + h.c., \tag{6}$$

where $\boldsymbol{\epsilon}_{d,c}(\boldsymbol{r})$ represent the field profiles at position $\boldsymbol{r}$, $\phi_{d,c}$ are the respective phases, $\gamma_s$ is the spontaneous emission rate into free space, and $2\kappa$ is the energy decay rate in the cavity. The spectrum measured for such a system is given by[11]

$$S(\omega) \propto \mathrm{Re}\left[\int_0^\infty dt \int_0^\infty dt'\, e^{i\omega(t-t')}\langle\hat{\mathbf{E}}^\dagger(\boldsymbol{r},t)\cdot\hat{\mathbf{E}}(\boldsymbol{r},t')\rangle\right]. \tag{7}$$

This spectrum can be derived from two-time correlators which, following the quantum regression theorem, evolve as[12]

$$\frac{\mathrm{d}}{\mathrm{dt}'}\langle\hat{a}^\dagger(t)\hat{a}(t')\rangle = -(\mathrm{i}\omega_\mathrm{c} + \kappa)\langle\hat{a}^\dagger(t)\hat{a}(t')\rangle + g\langle\hat{a}^\dagger(t)\hat{\sigma}_-(t')\rangle, \tag{8}$$

$$\frac{\mathrm{d}}{\mathrm{dt}'}\langle\hat{a}^\dagger(t)\hat{\sigma}_-(t')\rangle = -(\mathrm{i}\omega_\mathrm{d} + \Gamma_d)\langle\hat{a}^\dagger(t)\hat{\sigma}_-(t')\rangle - g\langle\hat{a}^\dagger(t)\hat{a}(t')\rangle, \tag{9}$$

$$\frac{\mathrm{d}}{\mathrm{dt}'}\langle\hat{\sigma}_+(t)\hat{a}(t')\rangle = -(\mathrm{i}\omega_\mathrm{c} + \kappa)\langle\hat{\sigma}_+(t)\hat{a}(t')\rangle + g\langle\hat{\sigma}_+(t)\hat{\sigma}_-(t')\rangle, \tag{10}$$

$$\frac{\mathrm{d}}{\mathrm{dt}'}\langle\hat{\sigma}_+(t)\hat{\sigma}_-(t')\rangle = -(\mathrm{i}\omega_\mathrm{d} + \Gamma_d)\langle\hat{\sigma}_+(t)\hat{\sigma}_-(t')\rangle - g\langle\hat{\sigma}_+(t)\hat{a}(t')\rangle, \tag{11}$$

where $\Gamma_d = \frac{\gamma_s}{2} + \gamma_p$ represents the dephasing of the dipole, due to both spontaneous emission and pure dephasing. In our system $\Gamma_d \gg \gamma_s, \kappa, g$. After non-resonant excitation, our system further begins in the state where the emitter is excited and the cavity is empty. Under these assumptions, and defining $\Delta\omega_{d,c} = \omega_{d,c} - \omega$, equations (8)–(11) give[12]

$$C_{cc}(\omega) = \boldsymbol{F}[\langle\hat{a}^\dagger(t)\hat{a}(t')\rangle] = \frac{g^2}{\Gamma_d\gamma_s\kappa}\frac{1}{\mathrm{i}\Delta\omega_\mathrm{c} + \kappa}, \tag{12}$$

$$C_{cd}(\omega) = \boldsymbol{F}[\langle\hat{a}^\dagger(t)\hat{\sigma}_-(t')\rangle] = \frac{g}{\Gamma_d\gamma_s}\frac{1}{\mathrm{i}\Delta\omega_\mathrm{d} + \Gamma_d}, \tag{13}$$

$$C_{dc}(\omega) = \boldsymbol{F}[\langle\hat{\sigma}_+(t)\hat{a}(t')\rangle] = \frac{g}{\Gamma_d\gamma_s}\frac{\mathrm{i}\Delta\omega_\mathrm{d} + 2\Gamma_d}{(\mathrm{i}\Delta\omega_\mathrm{d} + \Gamma_d)(\mathrm{i}\Delta\omega_\mathrm{c} + \kappa)}, \tag{14}$$

$$C_{dd}(\omega) = \boldsymbol{F}[\langle\hat{\sigma}_+(t)\hat{\sigma}_-(t')\rangle] = \frac{1}{\gamma_s}\frac{1}{\mathrm{i}\Delta\omega_\mathrm{d} + \Gamma_d}, \tag{15}$$

where $\boldsymbol{F}[g(t,t')] = \int_0^\infty dt \int_0^\infty dt'\, e^{i\omega(t-t')}g(t,t')$. These functions, combined with equations (6) and (7), give the collected spectrum

$$S(\omega) \propto \mathrm{Re}\left[\frac{1}{\mathrm{i}\Delta\omega_\mathrm{d} + \Gamma_d}\left(1 + \frac{\kappa}{\Gamma_d}\frac{1}{\mathrm{i}\Delta\omega_\mathrm{c} + \kappa}\left[F_0(\mathrm{i}\Delta\omega_\mathrm{d} + \Gamma_d) + \sqrt{F_0}(\mathrm{i}\Delta\omega_\mathrm{d} + 2\Gamma_d)e^{i\Delta\phi}\right]\right)\right], \quad (16)$$

where we have simplified by assuming strong overlap between the direct and cavity emission ($\epsilon_d \approx \epsilon_c$), and $\Gamma_d \gg \gamma_s, \kappa, g, |\omega_d - \omega_c|$. We further defined $\Delta\phi = \phi_d - \phi_c$, and a coupling parameter, $F_0 = 2g^2/\kappa\gamma_s$.

This model was fitted to the experimental data, as shown in Fig. 5f. This model produces a line shape that approximately follows a Fano profile with a Lorentzian envelope, with the fitted parameters presented in Supplementary Table 2. Our moderate coupling parameter ($F_0$) gives rise to the Fano-type profile, due to coherent interference between the direct and cavity-mediated emission of the QR.

**Supplementary Note 9. Image processing and statistics**

Raw AFM files (.h5) were generated by the proprietary *Ergo* software operating the Asylum Jupiter XR instrument (Oxford Instruments). These raw files can be opened and processed using the open-source software *Gwyddion*. Most AFM files acquired in this study were processed using the "Analysis" mode of the *Ergo* software. Specifically, the height channels or z-sensor channels of the AFM files were processed with standard masking and flattening, followed by exporting as .png images. Raw AFM files of QR-origami with $SiN_x$ chips were processed with *Gwyddion* using the three-point flattening function to level data to the top surface of the $SiN_x$ structures. Cross-section height profiles of structures were measured using either of the two software packages mentioned above.

PMMA well sizes of the rhombic wells after EBL with various doses were acquired from the AFM images of corresponding samples using a MATLAB script. The script cleans up the image, applies thresholds, and detects the boundary of the wells, followed by calculation of the widths and the lengths of the wells by fitting the boundary shape with a bounding box.

Origami, gold NP and QR placement yields and distributions were acquired from the AFM images of corresponding array samples with the help of a MATLAB script. The script cleans up the image, applies thresholds, and detects and labels connected objects (target structures). The script then allows manual correction of mislabeled objects and finally counts the number of labeled items. Data from two to five images are combined to calculate the placement yield and the distribution ($N$ = 288 to 800 total sites including empty sites). QR placement yields on $SiN_x$ photonic structures can be hard to determine due to the quality of the AFM images. AFM imaging of nanometer-sized QRs bound to the 6HB origamis with a similar height requires AFM probes with sharp tips and steady imaging parameters to resolve the QR from the origami structure. Imaging QR-origami on top of 310-nm-high $SiN_x$ structures is very challenging due to the drastic changes in the scanning height profile, resulting in frequently dulled or damaged AFM probes. Hence, when counting QRs on $SiN_x$ structures from AFM images, ambiguous sites of the QR-origami structure were not counted. Attempts to image QR-origami with SEM were also unsuccessful in resolving the QRs, potentially due to their small sizes and surface charging.

Placement precision was measured from AFM images of origami arrays after resist lift-off (3 × 3 µm, 512 × 512 pixels, 5.86 nm per pixel). The z-position-sensor channel was used rather than the z-drive channel. Images were flattened by a plane fit, a per-row low-percentile pre-leveling, and iterative per-row leveling against a height-masked background, applied to heights only so that lateral coordinates were preserved. Correctly placed single origamis were identified by visual inspection, then each was measured inside a 250 nm window using a local background taken from the window's outer annulus, with its footprint defined as the half-maximum contour of its own peak height so that the centroid is independent of the absolute height threshold. Sites whose footprint was clipped by the image border, or whose footprint area fell outside 0.45–1.8 times the single-origami footprint of that image, were excluded from the positional statistics (1.1%). Because the EBL array is a square lattice of accurately known 250 nm pitch, only its pose in the AFM frame is unknown; an ideal lattice was registered to the measured centroids by robust least squares with both lattice vectors free, which simultaneously calibrates the AFM scanner against the lithographic pitch (fitted pitch 250.51 ± 0.16 nm and 254.33 ± 0.72 nm along the fast and slow axes, $N$ = 13 images). The reported displacements are therefore site-to-site precision: a displacement common to all origamis within an image is absorbed by the pose fit and is not observable after lift-off. Placement precision is reported as the standard deviation of the displacements of all origamis of a given well size along each axis, and its uncertainty as the standard error of the per-image standard deviations (N = 4, 5 and 4 images for well sizes 1, 2 and 3). All processed data are available in the Dryad repository[10].

Rhombic origami orientation and QR orientation were measured from the AFM images of corresponding array samples with the help of a MATLAB script. The script allows the user to draw lines on the AFM images and record their orientations within −90° to 90°, given the two-fold rotational symmetry. However, when calculating the standard deviation for vertically aligned samples, angles between 0° and −90° were added to 180° to calculate the distribution around 90° (e.g., 90° ± 30°). For origami orientations, the line was drawn along the long diagonal axis connecting the two opposing vertices of the rhombic structure. For QR orientations, the line was drawn along the elongation direction of the rod shape. Orientation distribution was plotted, and the accuracy was assessed as the percentage of origamis/QRs with angles in a certain range of the target orientation. Note that the AFM images might contain slight drifts and/or global image rotations which render the absolute mean angle of individual structures less relevant.

Widefield PL images were acquired from raw videos collected using the open-source software *µManager* at 100 ms per frame for 100 frames (10 s). The raw data was processed using *FIJI* to project the maximum value of pixels over the 100 frames onto one image (maximum-projection image), to showcase the PL of all individual QRs despite blinking over the 10 s period. Confocal PL image files were collected using the *PicoQuant SymPhoTime* software. The raw data can be rendered using said software or open-source Python code and exported as image files. To compensate for QR blinking during the raster scan, 6 PL images were collected and averaged for each analyzer angle. A MATLAB script was employed to extract the average PL intensity of an area of interest, which was chosen manually to avoid extra bright spots on the PL image (potential aggregations). The PL intensity of the same area of interest was extracted over various analyzer angles and plotted to demonstrate the controllable emission polarization.

The reproducibility of the TMOP method is evident considering its repeated usage on different origami shapes and designs (6HB wireframe rhombus and ladder shape, and single-layer triangle), and its usage under different conditions for gold NP and QR templating on silicon chips. To optimize the NP templating, around 30 individual experiments (chips) were carried out for gold NPs (including AuNSs and AuNRs), and more than 25 individual experiments were carried out for QRs. AFM images acquired from separate optimized experiments were combined to count and calculate the yield (i.e., sample sizes from separate experiments of the same optimized conditions, if available, were combined into the *N* reported). To test TMOP-directed QR incorporation with $SiN_x$ photonic structures, 6 individual experiments were carried out, including 2 chips with the optimized condition.

Raw image files, additional images, processed data, and all scripts used for image processing are available on the open-source online data repository Dryad[10].

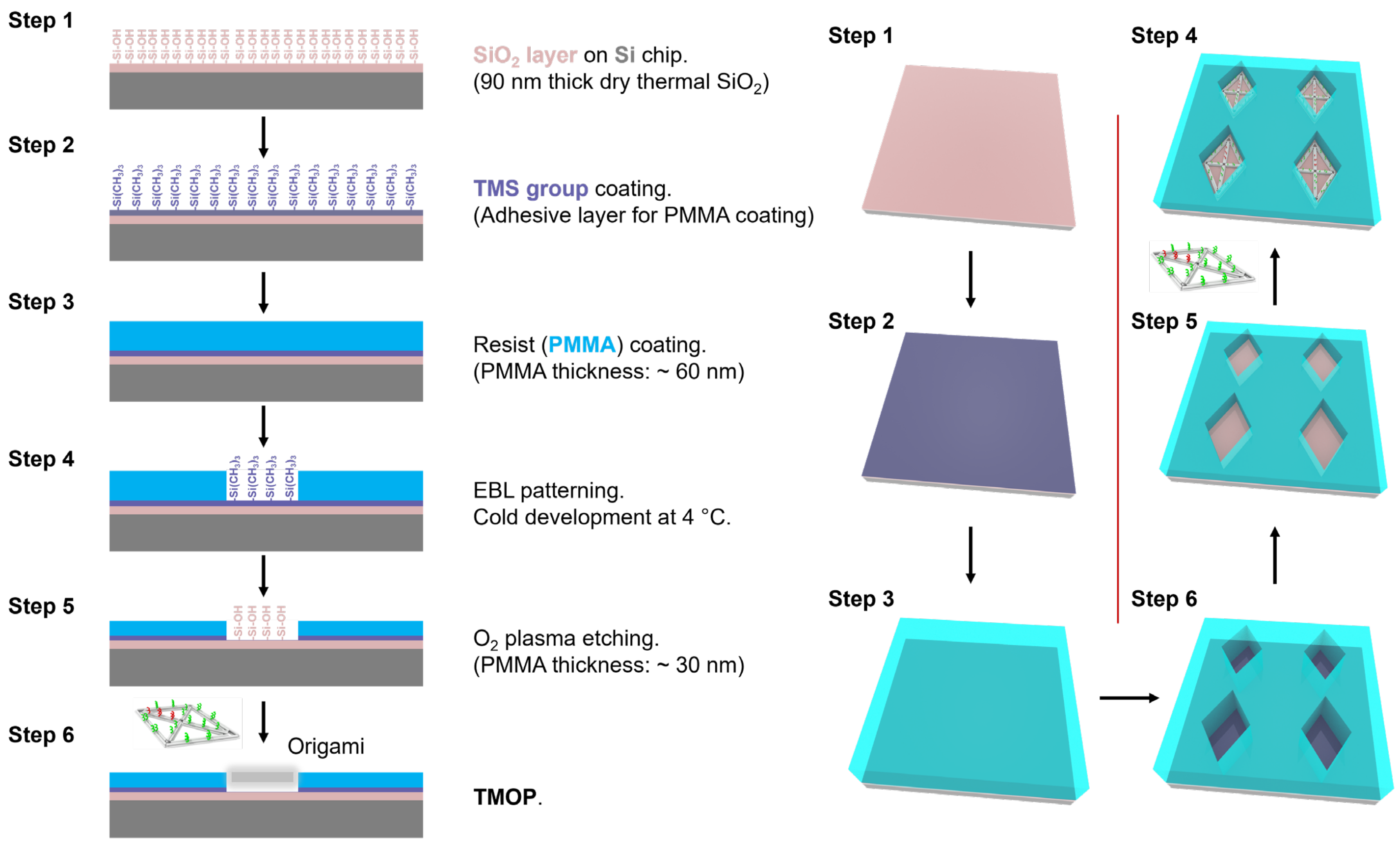


**Supplementary Fig. 1 | Schematic illustration of the fabrication of PMMA wells on silicon chips for TMOP.**

a

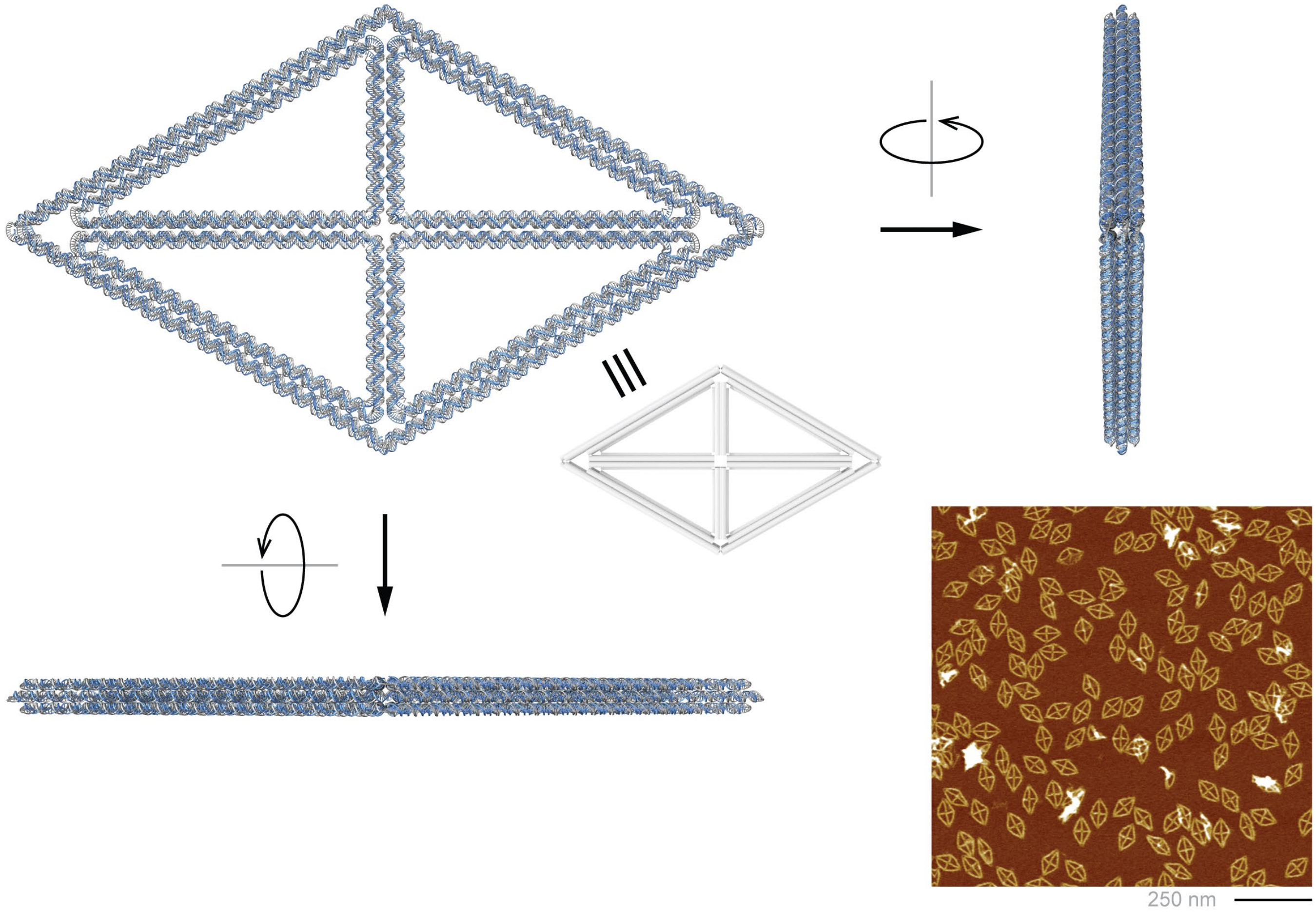


b

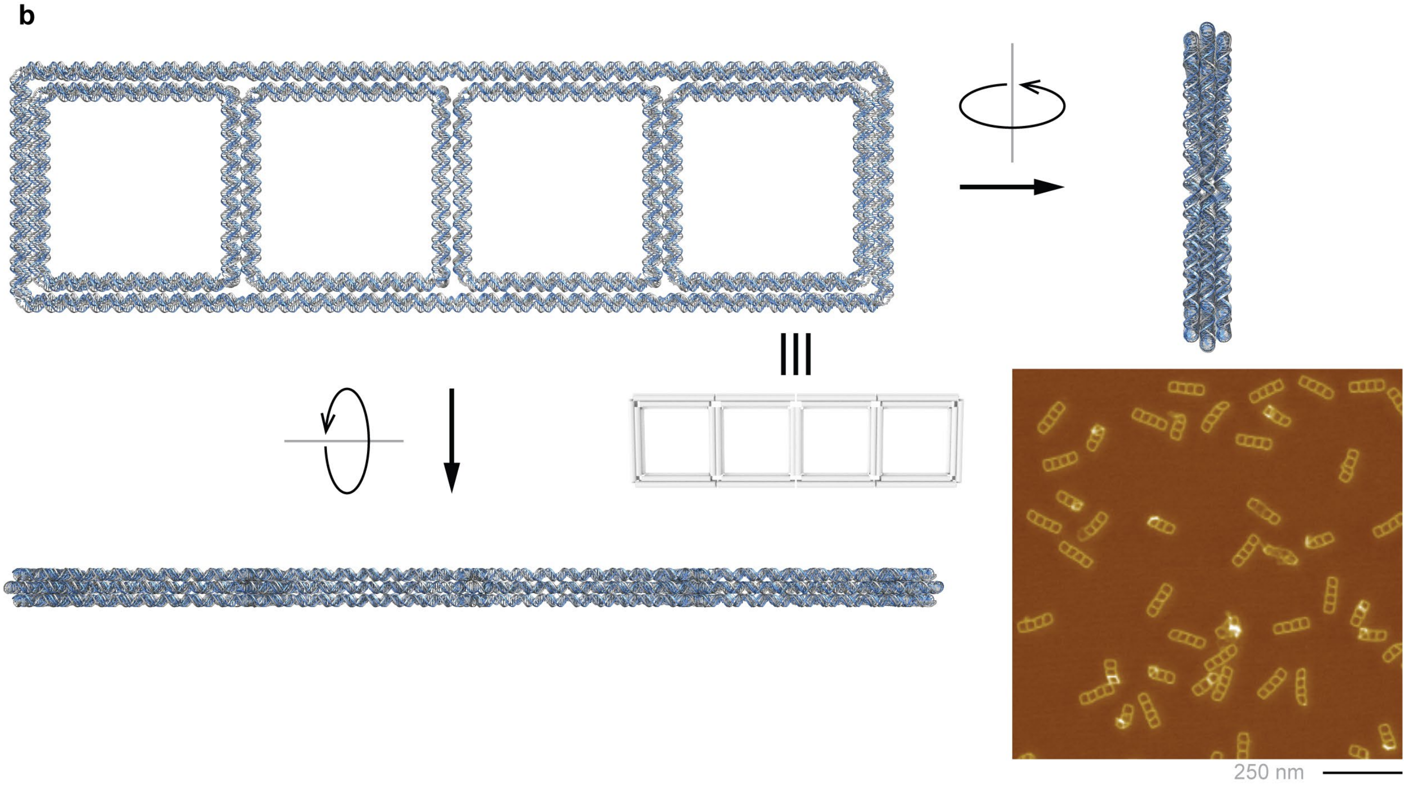


**Supplementary Fig. 2 | Schematic illustration and AFM images of the two-dimensional 6HB wireframe origami designs. a**, 6HB rhombic origami. **b**, 6HB ladder-shaped origami.

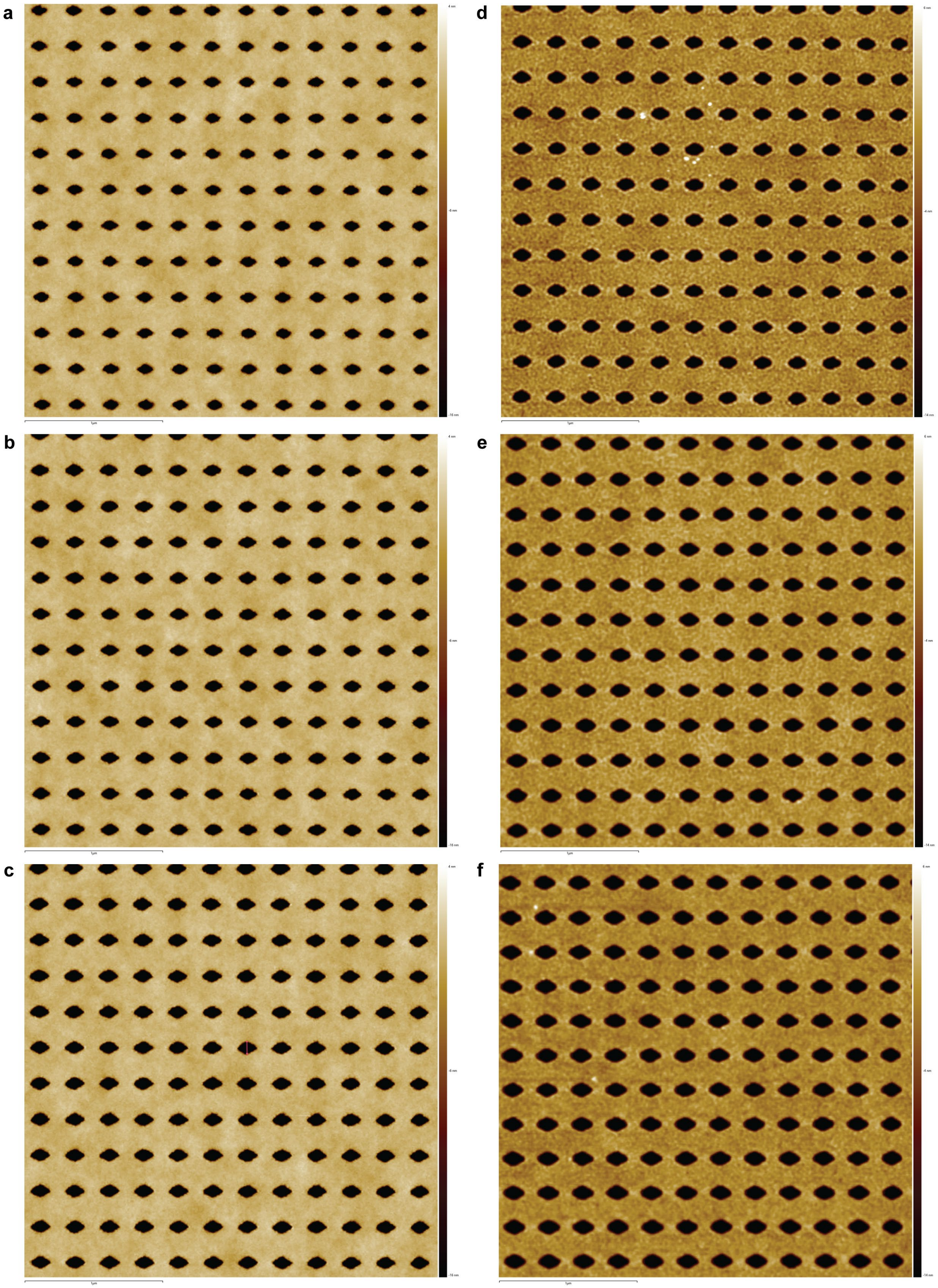


**Supplementary Fig. 3 | AFM images of PMMA well size 1 (a,d), size 2 (b,e) and size 3 (c,f) before (a–c) and after (d–f) $O_2$ plasma reactive-ion etching.**

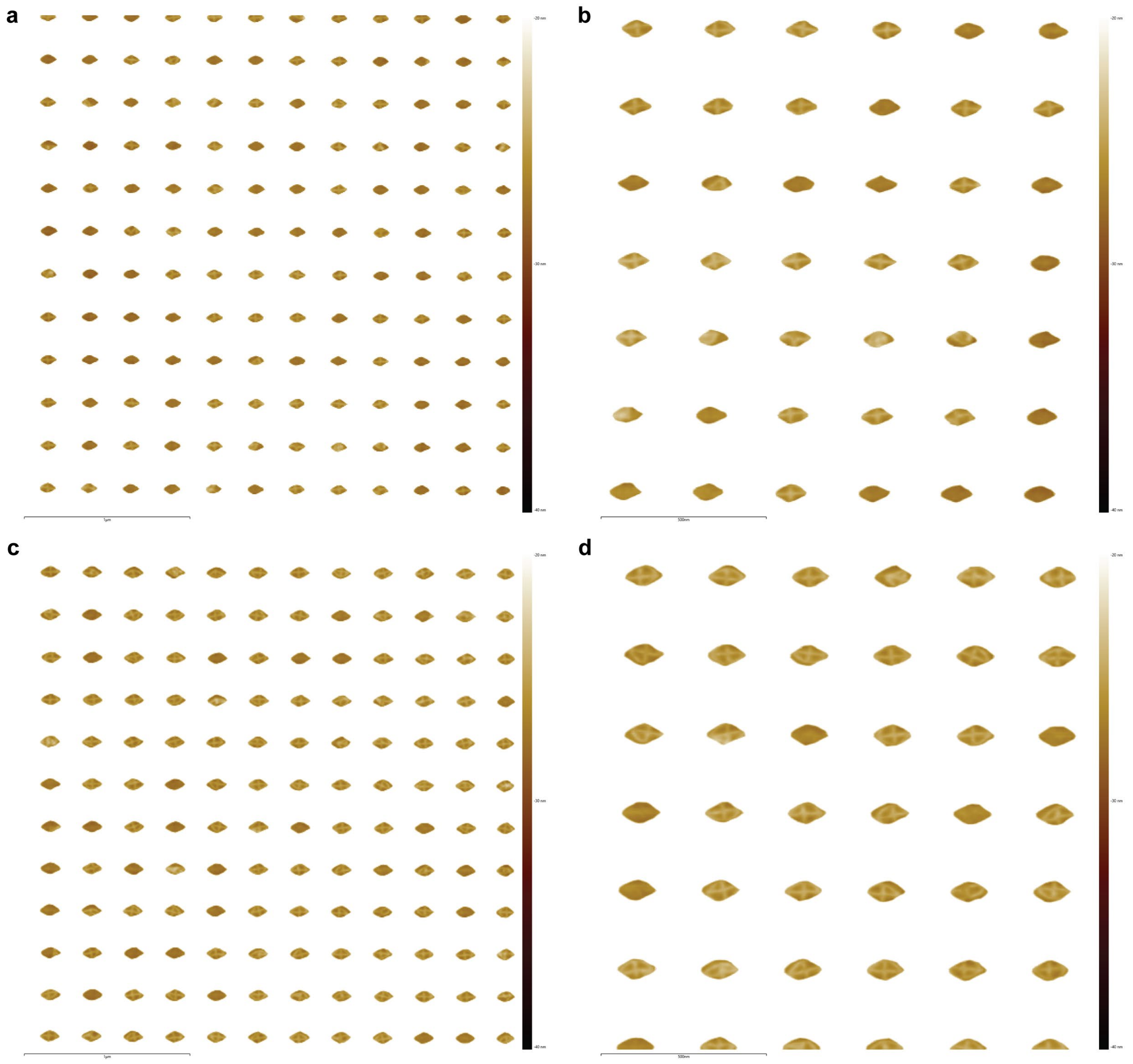


**Supplementary Fig. 4 | AFM images of rhombic origami placed at the base of PMMA wells. a,b**, Well size 1. **c,d**, Well size 2. No NaCl used for origami placement.

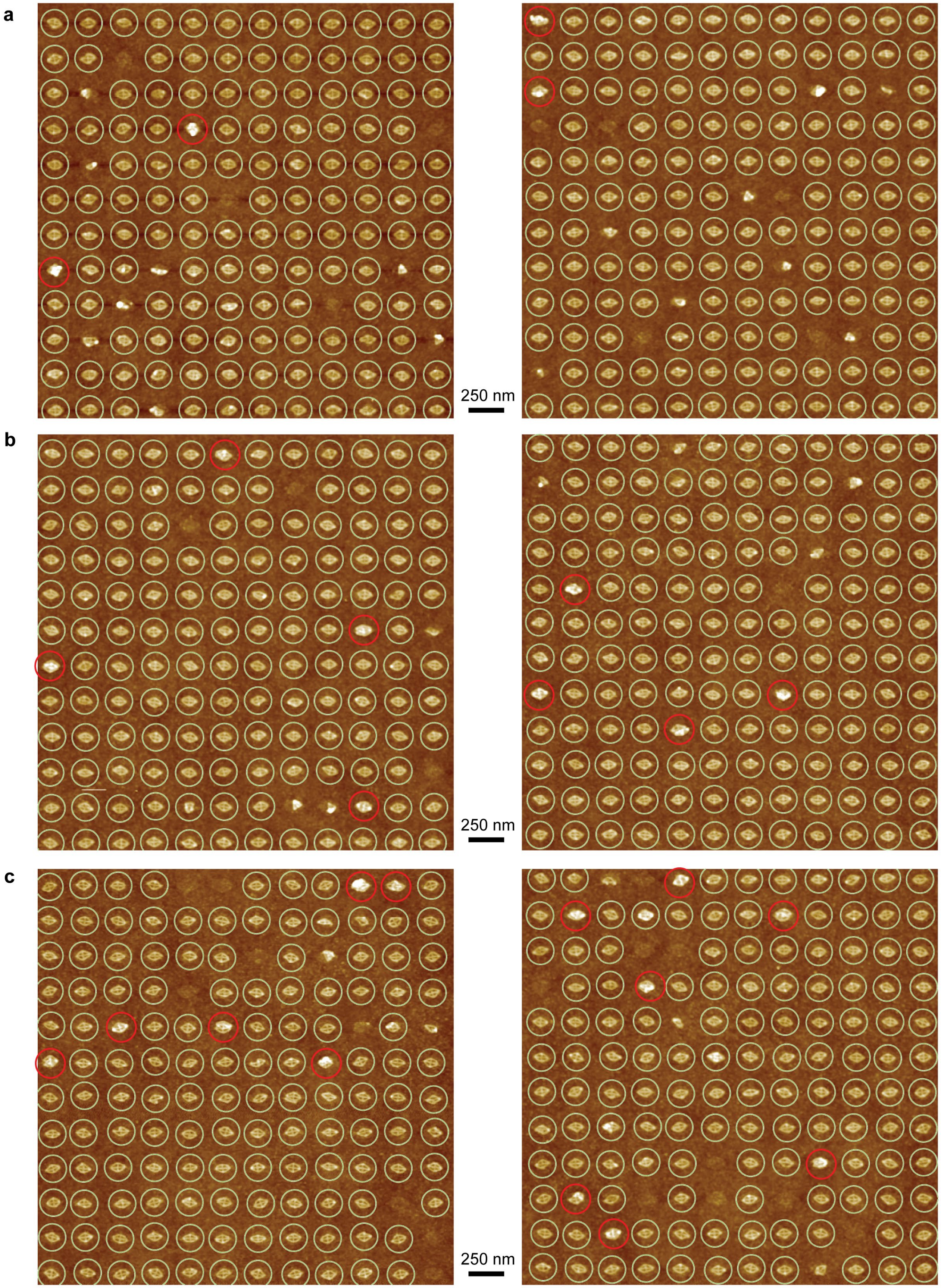


**Supplementary Fig. 5 | AFM images of TMOP-placed rhombic origami arrays with well size 1 (a), size 2 (b) and size 3 (c).** 100 mM NaCl used for placement. Multiple origamis placed at the same site were circled in red. Single-origami placement yields are 90%, 91%, and 88% for well sizes 1, 2, and 3.

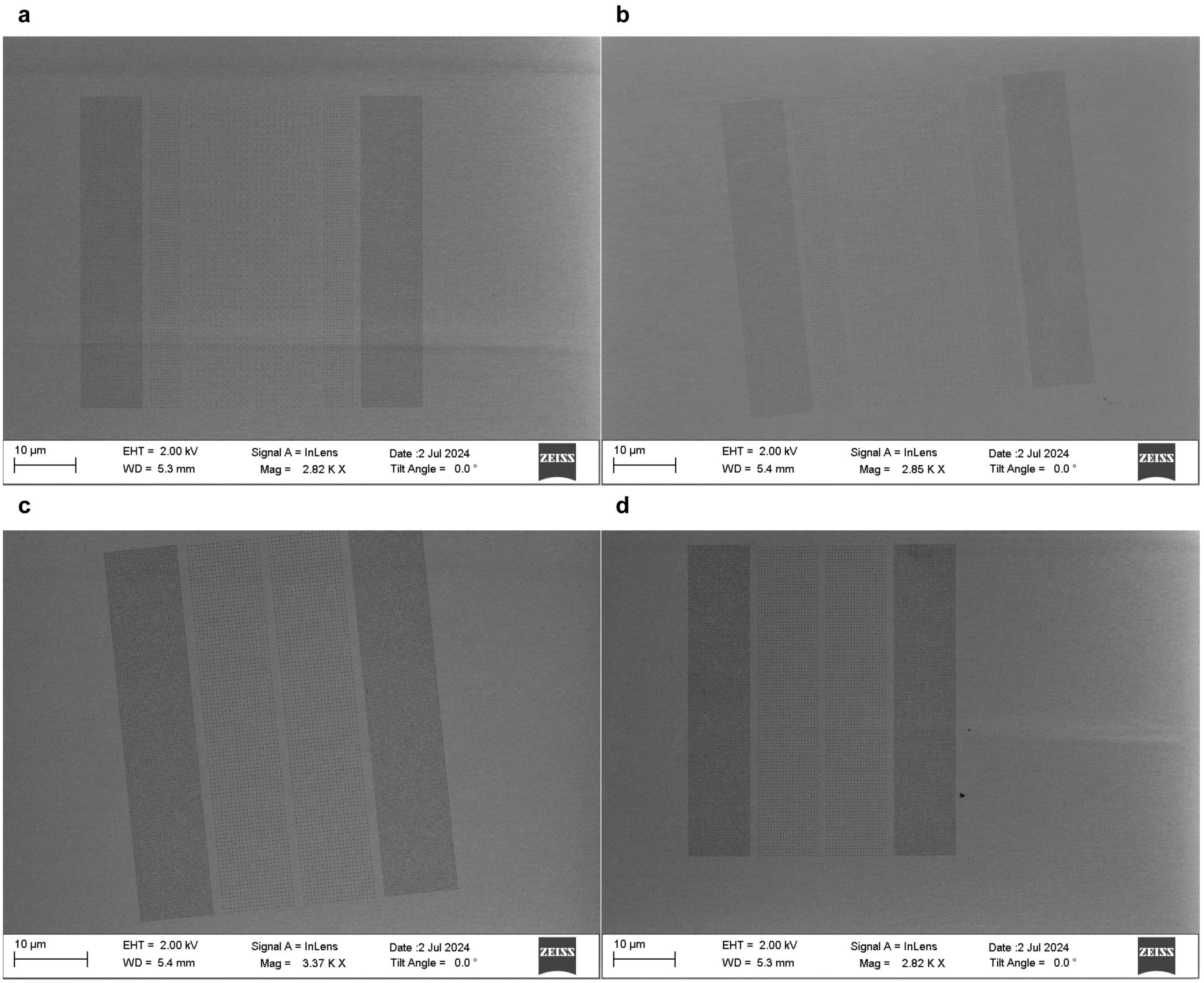


**Supplementary Fig. 6 | SEM images of additional TMOP-fabricated rhombic origami arrays containing 6 strips (used for QR arrangement) (a,b) and 4-strip arrays (used for AuNR arrangement) (c,d).**

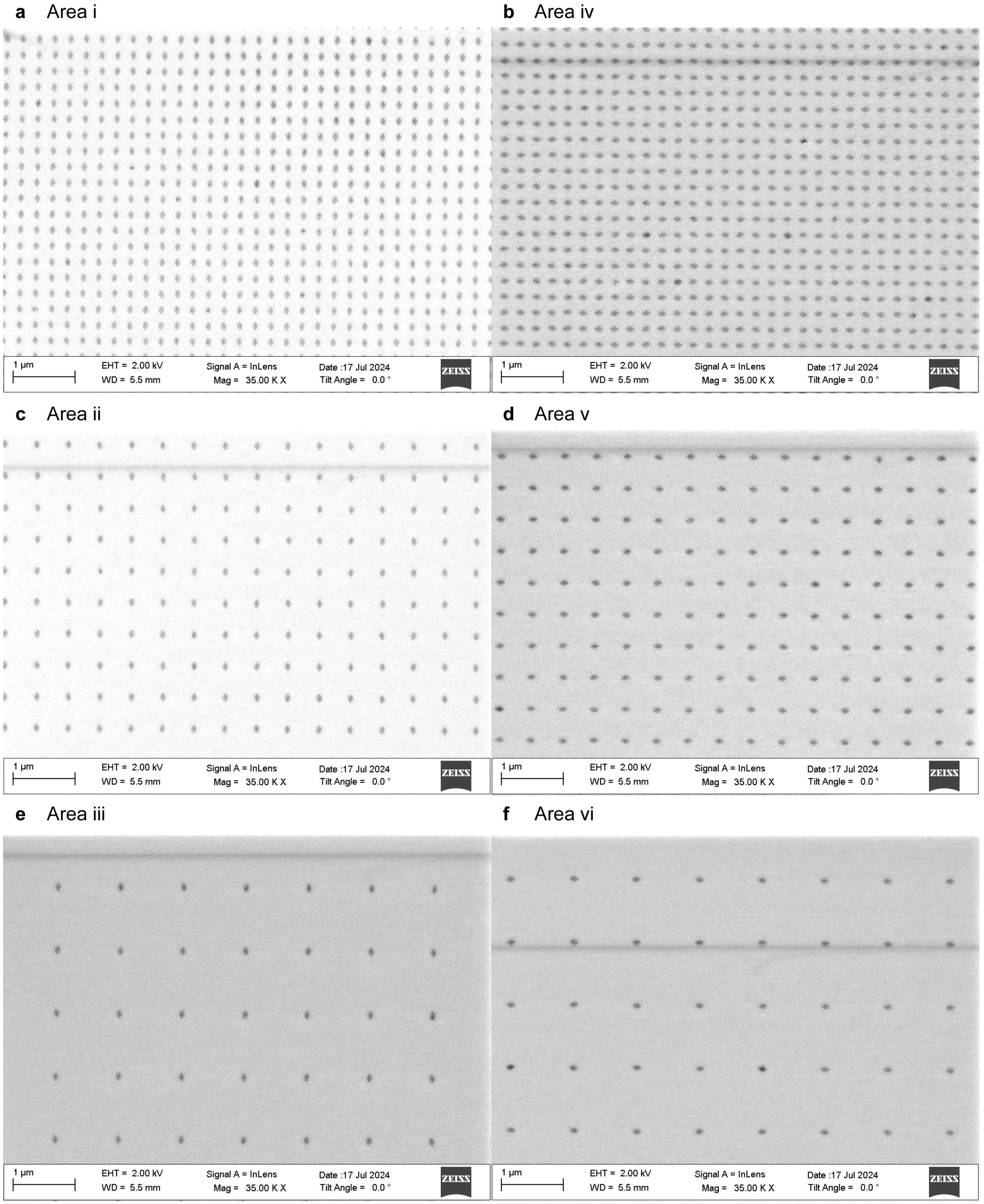


**Supplementary Fig. 7 | SEM images of the 6 strips in the TMOP rhombic origami array with 250 nm (a,b), 500 nm (c,d), or 1 µm (e,f) inter-origami spacings and vertical (a,c,e) or horizontal (b,d,f) orientations.**

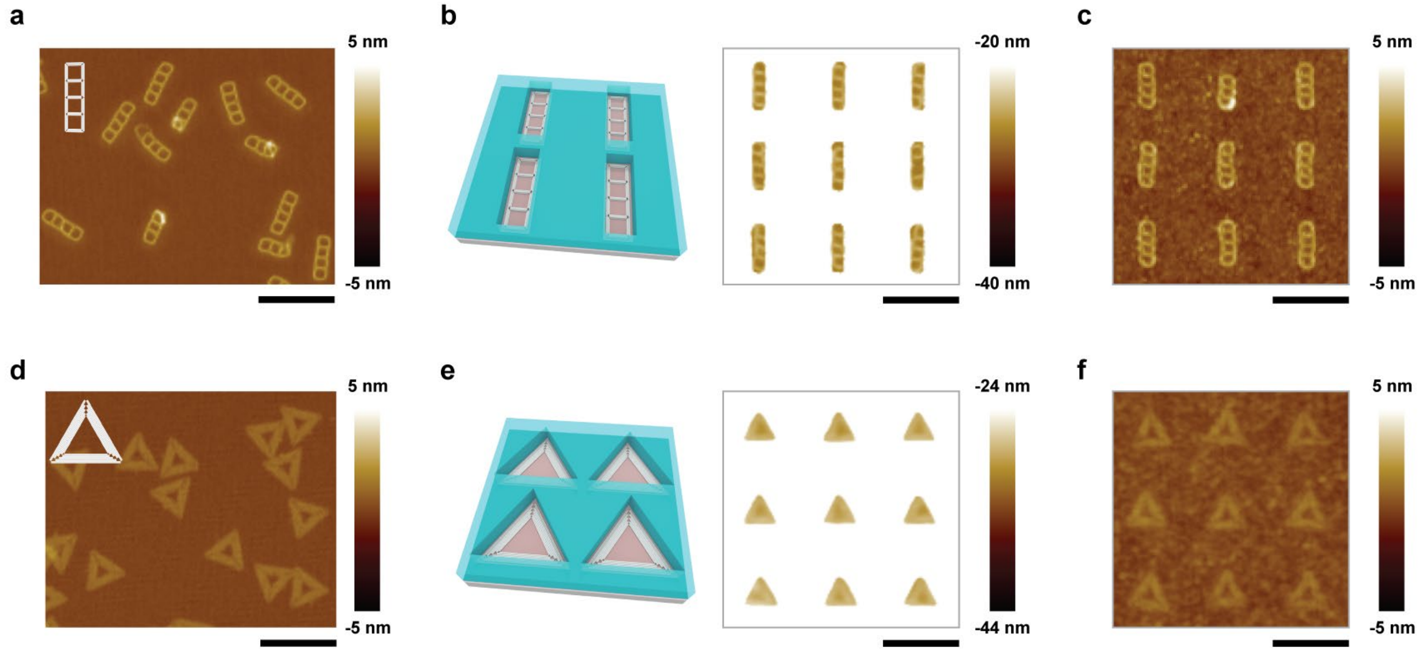


**Supplementary Fig. 8 | TMOP of a ladder-shaped 6HB wireframe origami (a–c) and a single-layer triangular origami (d–f). a,d**, Free origami deposited on mica. **b,e**, Origami placed at the bottom of the shape-matched wells. **c,f**, Origami array after lift-off. Scale bars, 250 nm.

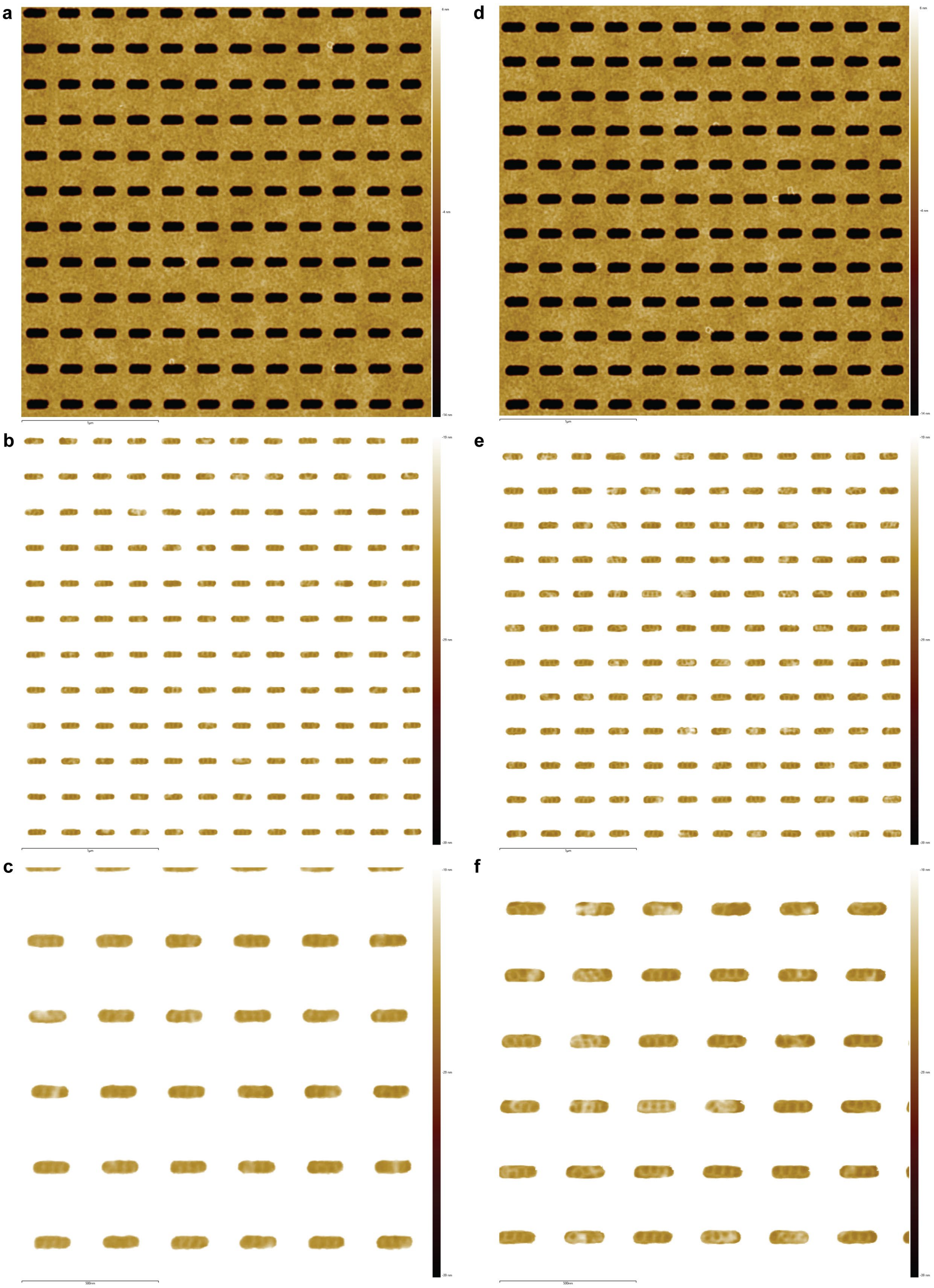


**Supplementary Fig. 9 | AFM images of ladder-shaped origami wells fabricated with 1,320 µC/cm$^2$ EBL dose (a) and 1,722 µC/cm$^2$ EBL dose (d).** Ladder-shaped origamis were placed at the bottom of the wells (b,c,e,f) after TMOP. No NaCl used for origami placement.

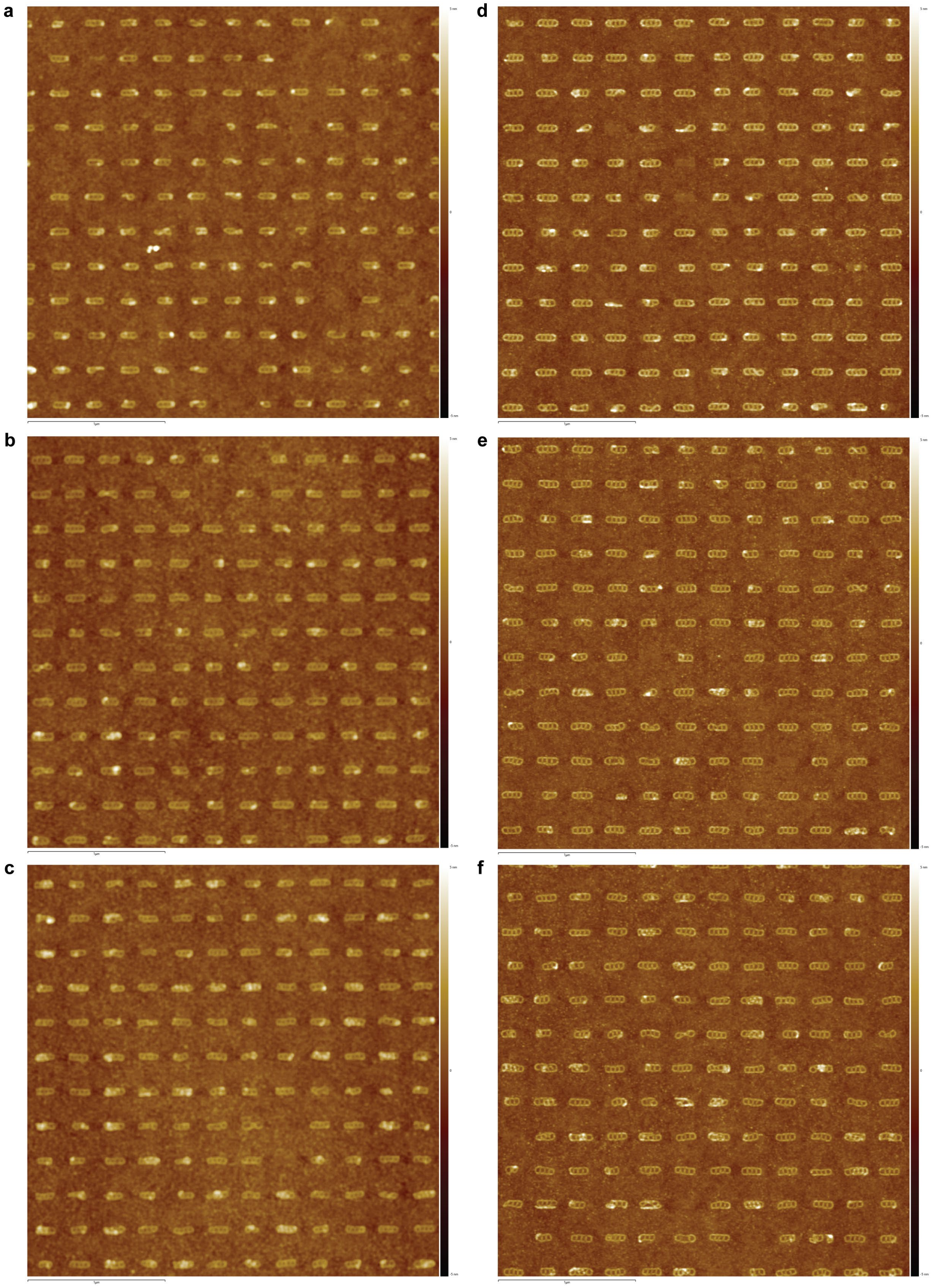


**Supplementary Fig. 10 | AFM images of TMOP-placed 6HB ladder-shaped origami arrays with wells fabricated with 1,020 µC/cm² (a,d), 1,320 µC/cm² (b,e) and 1,722 µC/cm² (c,f) EBL doses, respectively.** No NaCl used for placement in (a–c). 100 mM NaCl used for placement in (d–f).

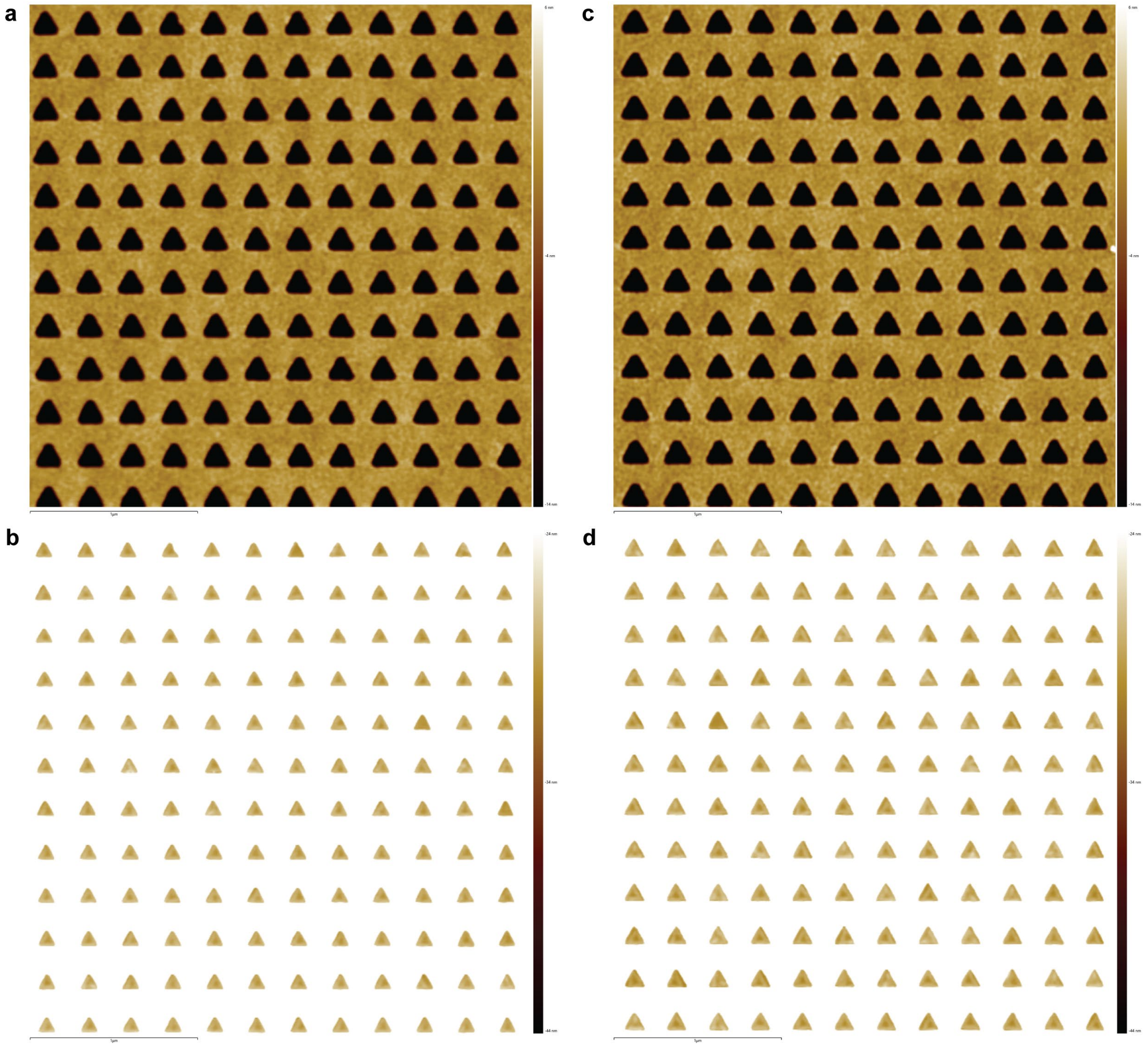

**Supplementary Fig. 11 | AFM images of single-layer triangle origami wells fabricated with 1,320 µC/cm$^2$ EBL dose (a) and 1,722 µC/cm$^2$ EBL dose (c).** Single-layer triangle origamis were placed at the bottom of the wells (b,d) after TMOP. 100 mM NaCl used for origami placement.

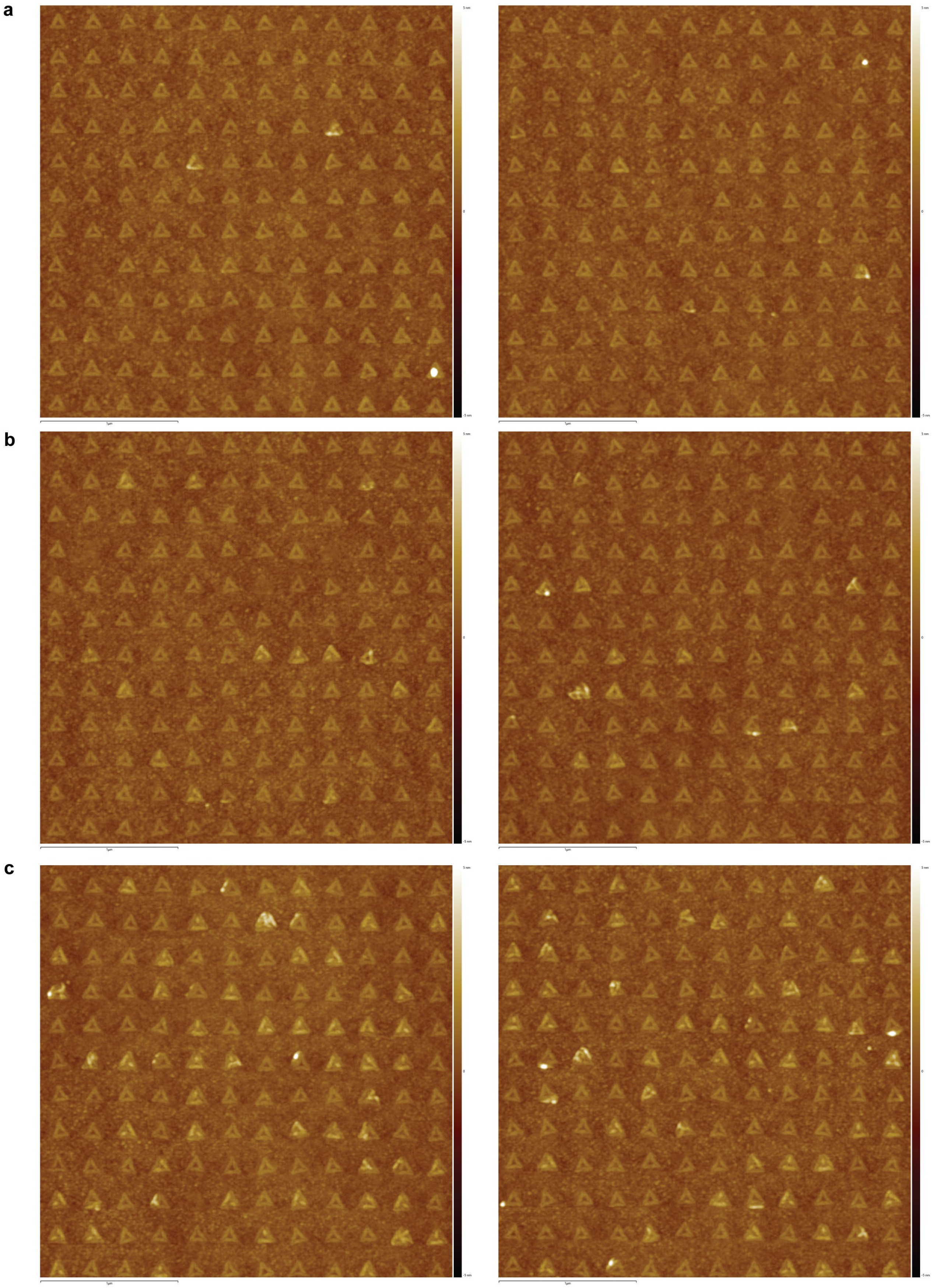


**Supplementary Fig. 12 | AFM images of TMOP-placed single-layer triangle origami arrays with wells fabricated with 1,020 μC/cm$^2$ (a), 1,320 μC/cm$^2$ (b) and 1,722 μC/cm$^2$ (c) EBL doses, respectively.** 100 mM NaCl used for placement.

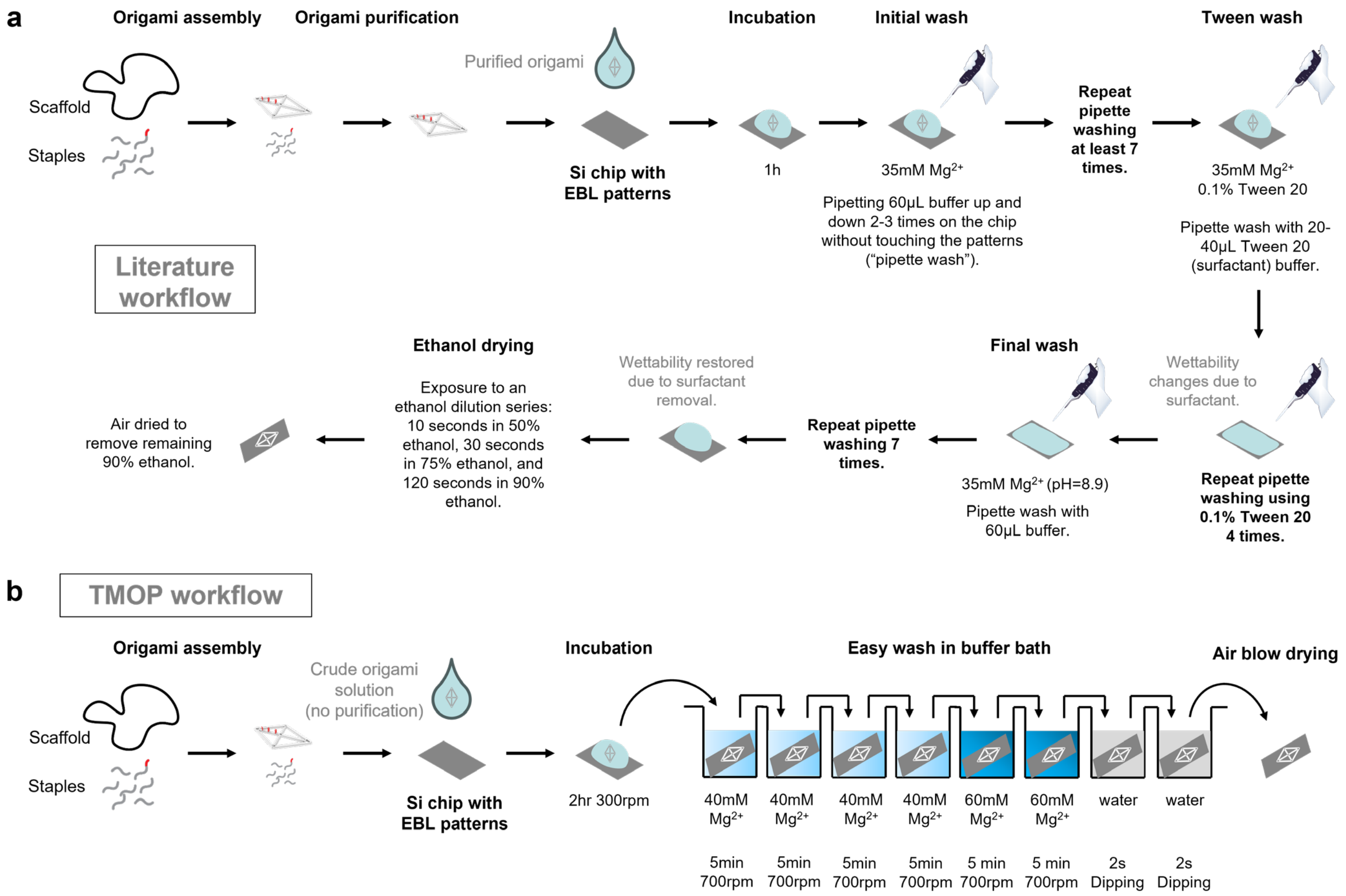


**Supplementary Fig. 13 | Schematic illustration of origami placement workflows.** **a**, Workflow of the DOP method from previous research by Gopinath et al.[2,5]. **b**, Workflow of the TMOP method developed in this study.

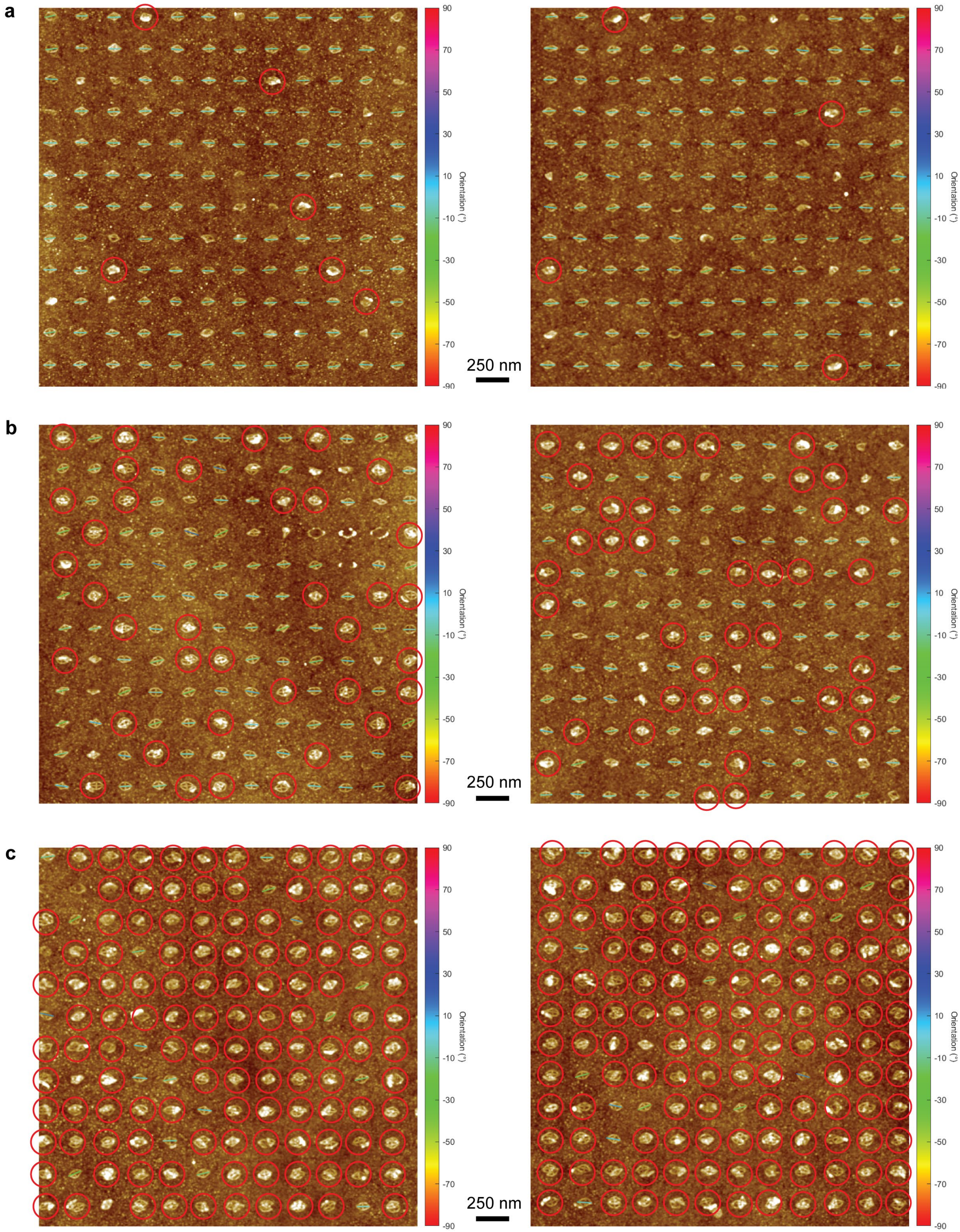


**Supplementary Fig. 14 | AFM images of TMOP-placed rhombic origami with well size 1 (a), size 2 (b) and size 3 (c).** No NaCl used for origami placement. Multiple origamis placed at the same site were circled in red. Single origami orientations were measured as the angle of the long axis direction.

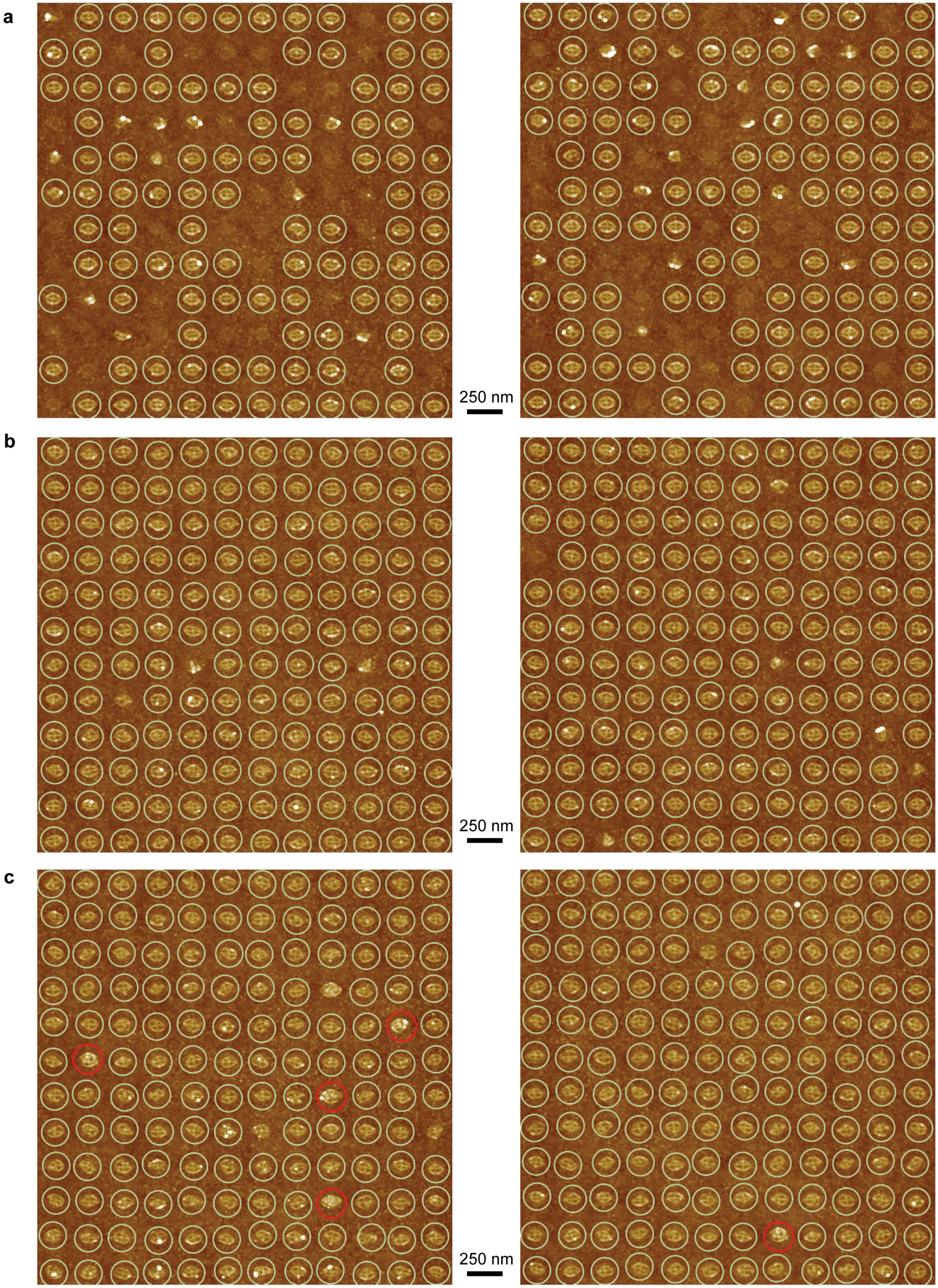
a
250 nm
b
250 nm
c
250 nm

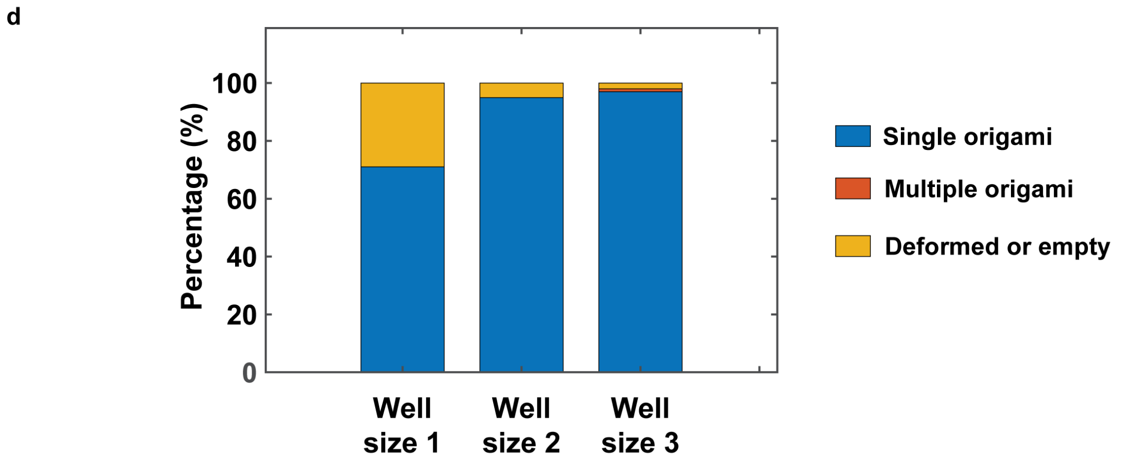


**Supplementary Fig. 15 | Silicification of TMOP rhombic origami arrays with APTES and TEOS. a–c**, AFM images of silicified rhombic origami arrays with well size 1 (a), size 2 (b) and size 3 (c). **d**, Statistics of TMOP origamis placed using wells of various sizes after silicification. 100 mM NaCl used for placement.

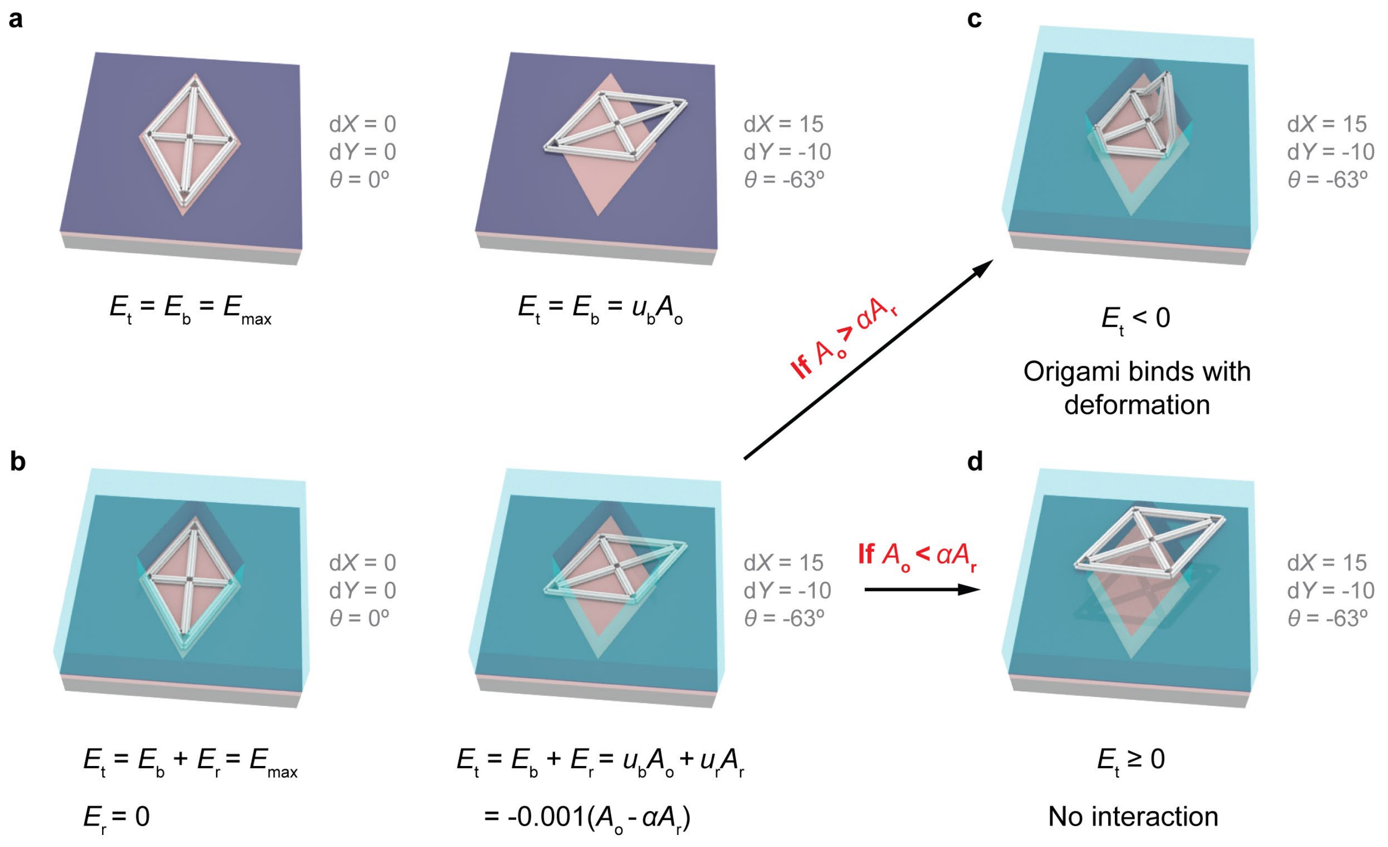


**Supplementary Fig. 16 | Total interaction free energy calculation for kinetic simulation of origami placement with landing pad (a) or with resist well (b). c**, If the binding free energy is greater than the calculated free energy penalty with a certain conformation (d$X$, d$Y$, $\theta$), origami can bind to the base of the well with certain structural deformation (real free energy penalty). **d**, If the binding free energy is smaller than the calculated free energy penalty, the origami cannot interact with the well base through electrostatic interaction with this specific conformation (d$X$, d$Y$, $\theta$).

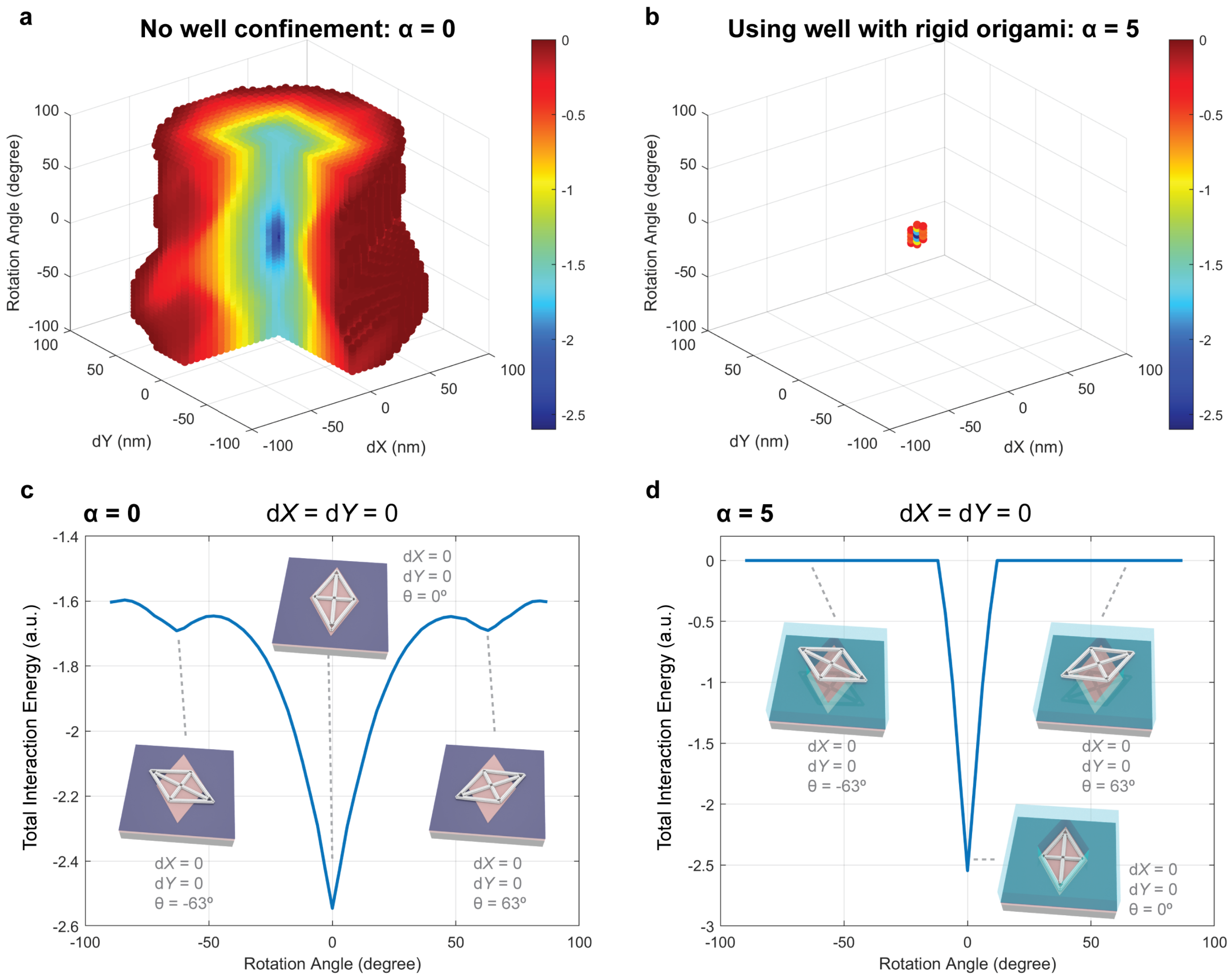


**Supplementary Fig. 17 | Total interaction free energy of a library of rhombic origami-target conformations with translational displacements (d*X* and d*Y*) and rotational displacement (*θ*) for origami-landing pad binding (a) and origami-well binding (b).** Data points of conformations with both d*X* ≤ 0 and d*Y* ≤ 0 are omitted for better data visualization. **c**, Origami-landing pad total interaction free energy (= binding free energy) profile as a function of rotation angle when d*X* = d*Y* = 0. **d**, Origami-well total interaction free energy profile as a function of rotation angle (*θ*) when d*X* = d*Y* = 0. Local free energy minima are highlighted with schematic illustrations. Note that conformations with total interaction free energy ≥ 0 were eliminated and represented with a total interaction free energy of 0, due to the lack of contact (i.e., binding) with the well base (d, top insets). In reality, these conformations do not exist as the origami will diffuse back into the bulk solution.

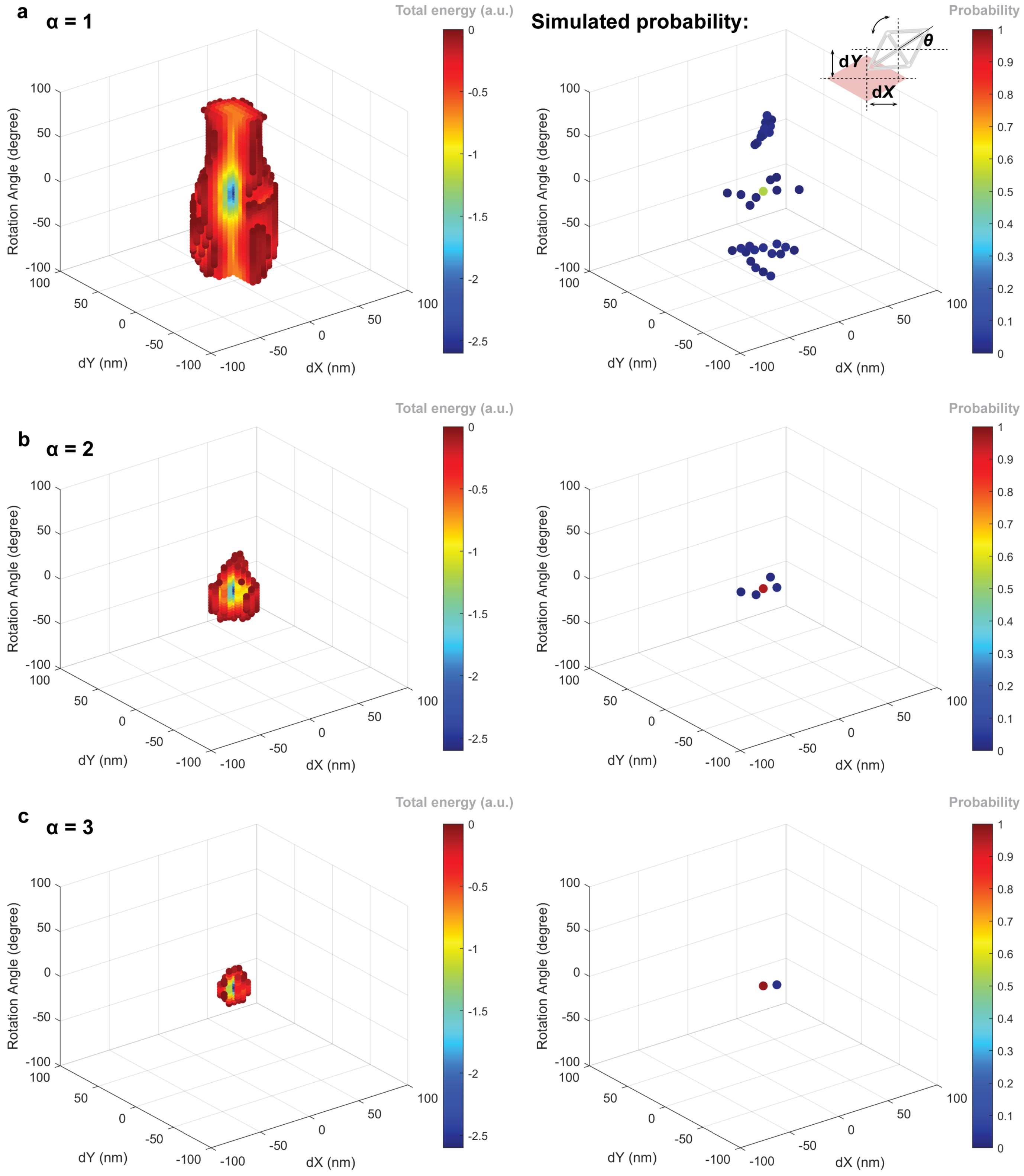


**Supplementary Fig. 18 | Total interaction free energy ensembles of rhombic origami-well conformations with translational displacements (d$X$ and d$Y$) and rotational displacement ($\theta$) (a–c left) and the simulated probability of the final conformations (a–c right) for origami-well binding with increasing free energy penalty coefficient ($\alpha$).** Conformations with total interaction free energy ≥ 0 were eliminated. Data points of conformations with both d$X \leq 0$ and d$Y \leq 0$ are omitted in total interaction free energy plots for better visualization (a–c left). Increasing free energy penalty coefficient ($\alpha$) correlates with increasing origami structural rigidity.

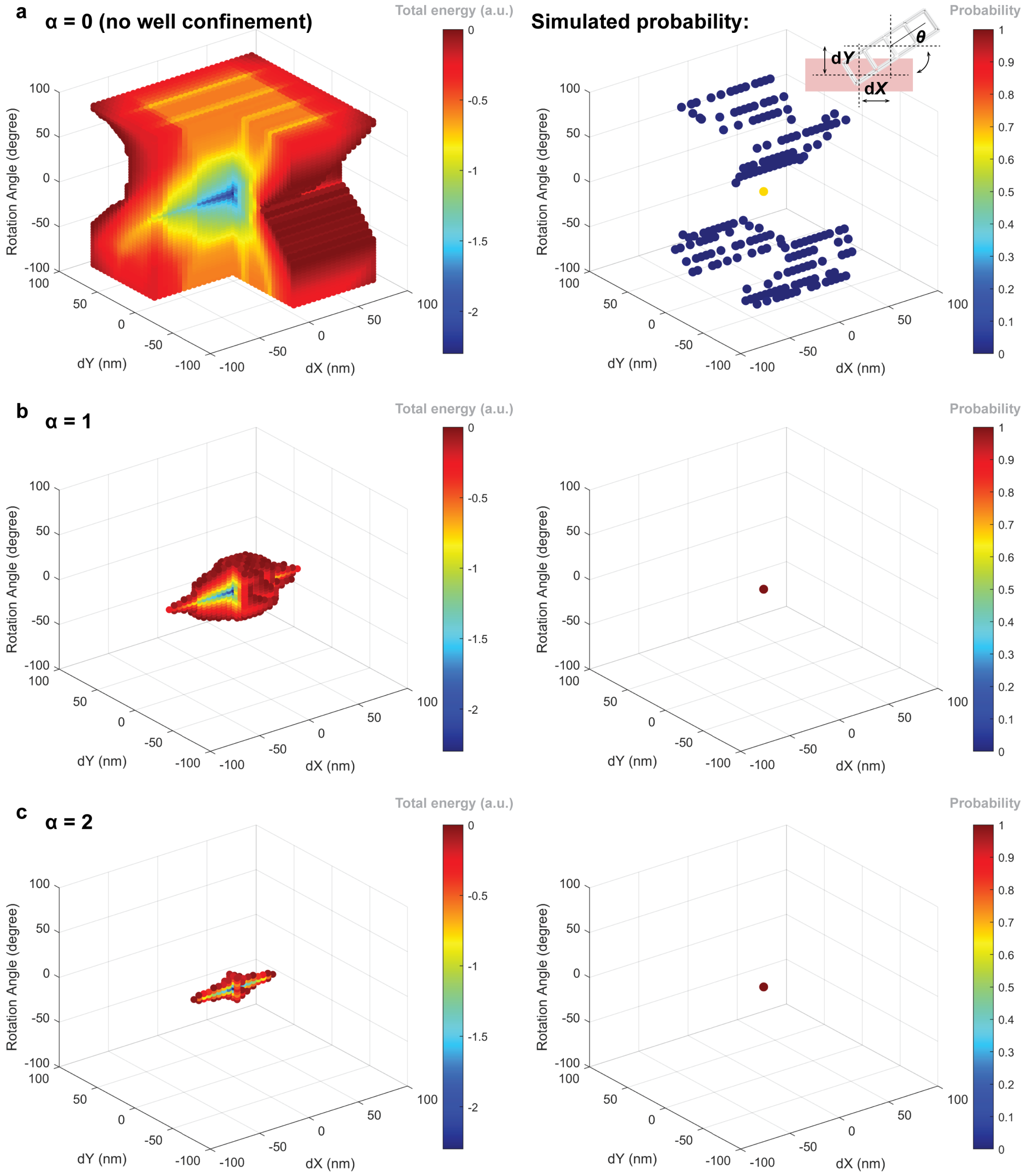
a
α = 0 (no well confinement)
Total energy (a.u.)
Rotation Angle (degree)
dY (nm)
dX (nm)
Simulated probability:
Probability
dY
dX
θ
b
α = 1
Total energy (a.u.)
Probability
c
α = 2
Total energy (a.u.)
Probability

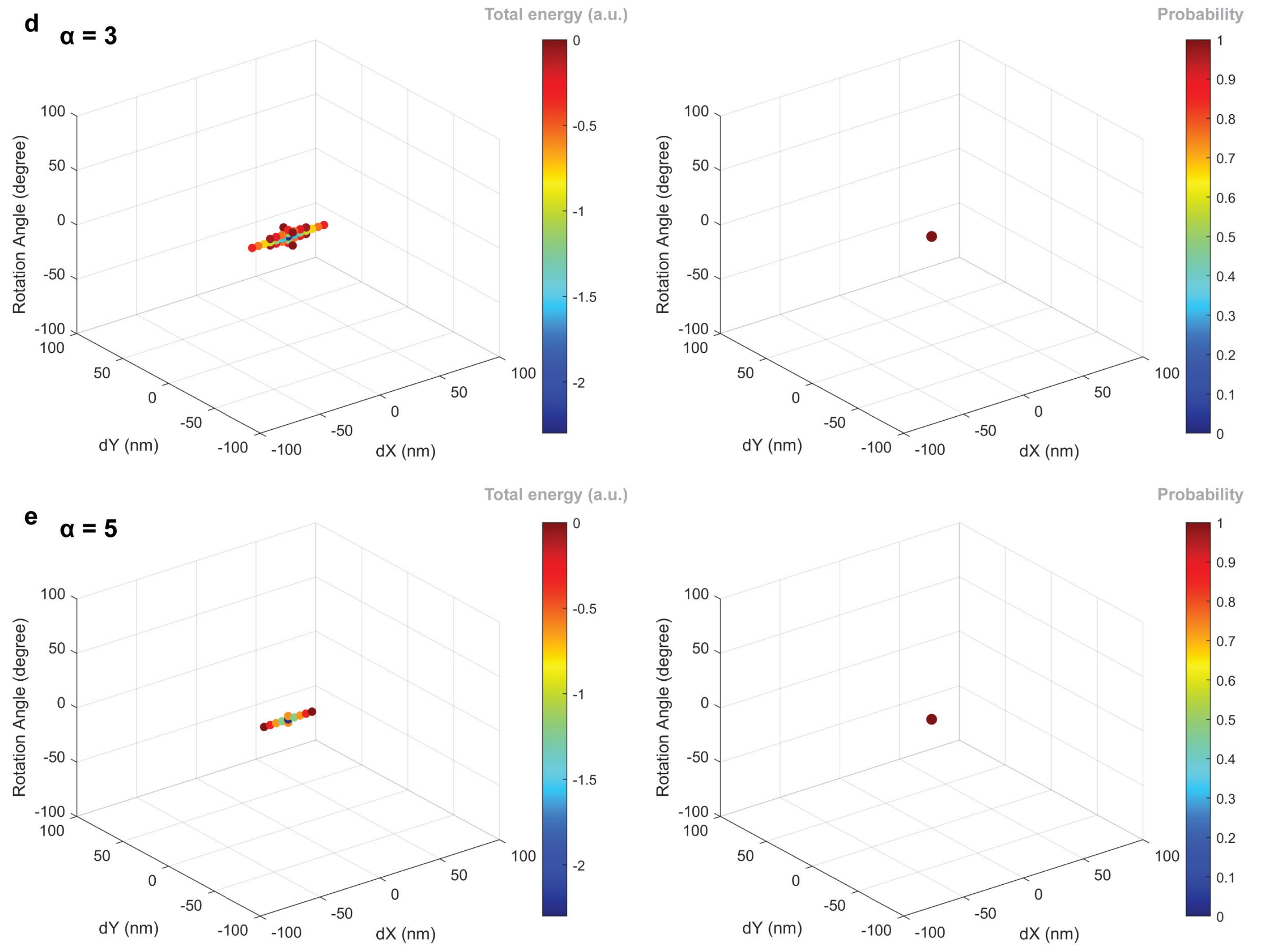


**Supplementary Fig. 19 | Total interaction free energy ensembles of ladder-shaped origami-well conformations with translational displacements (d$X$ and d$Y$) and rotational displacement ($\theta$) (a–e left) and the simulated probability of the final conformations (a–e right) for origami-well binding with increasing free energy penalty coefficient ($\alpha$).** Conformations with total interaction free energy ≥ 0 were eliminated. Data points of conformations with both d$X \leq 0$ and d$Y \leq 0$ are omitted in total interaction free energy plots for better visualization (a–e left). Increasing free energy penalty coefficient ($\alpha$) correlates with increasing origami structural rigidity.

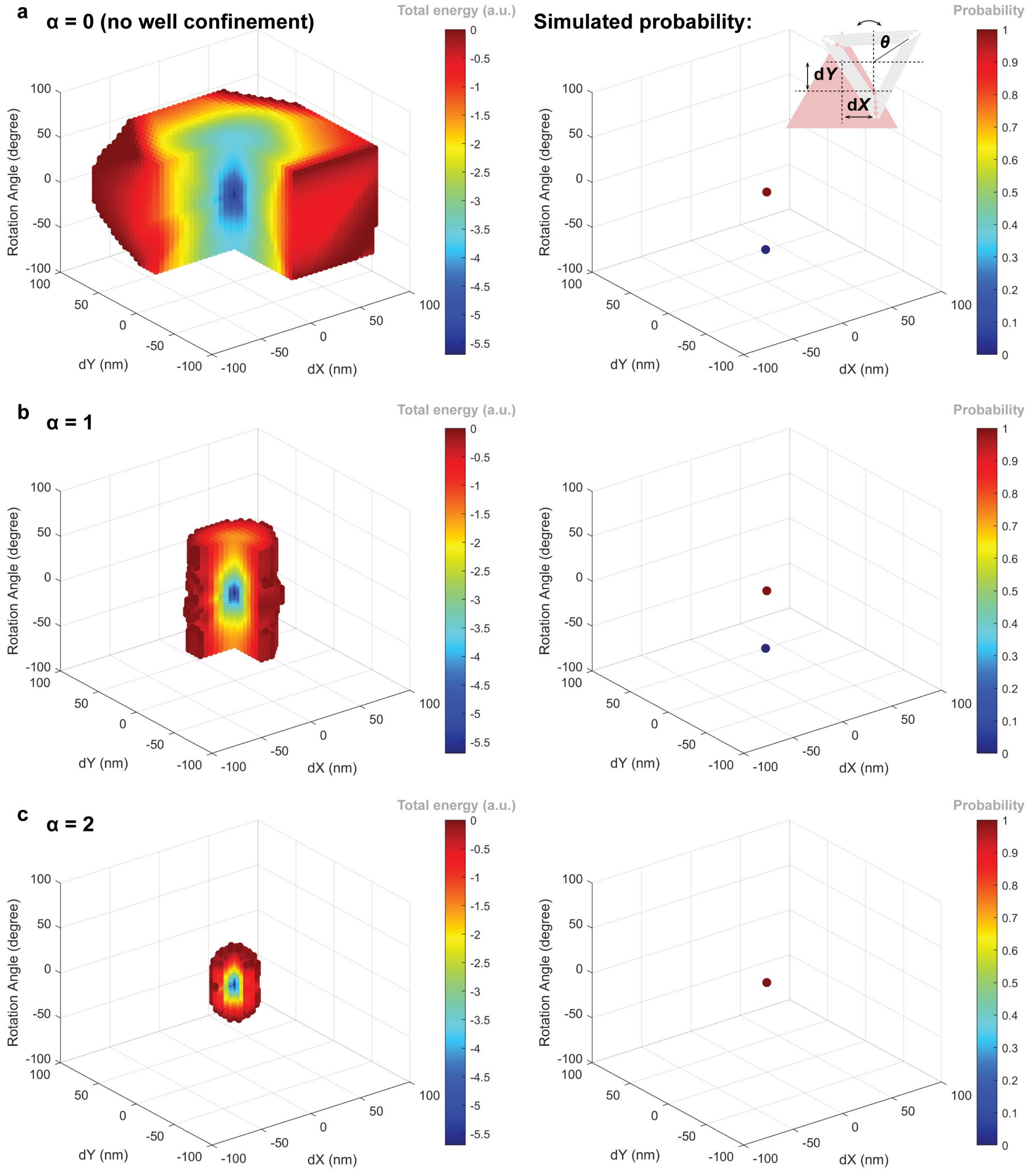
a
α = 0 (no well confinement)
Total energy (a.u.)
Simulated probability:
Probability
θ
dY
dX
Rotation Angle (degree)
dY (nm)
dX (nm)
b
α = 1
Total energy (a.u.)
Probability
Rotation Angle (degree)
dY (nm)
dX (nm)
c
α = 2
Total energy (a.u.)
Probability
Rotation Angle (degree)
dY (nm)
dX (nm)

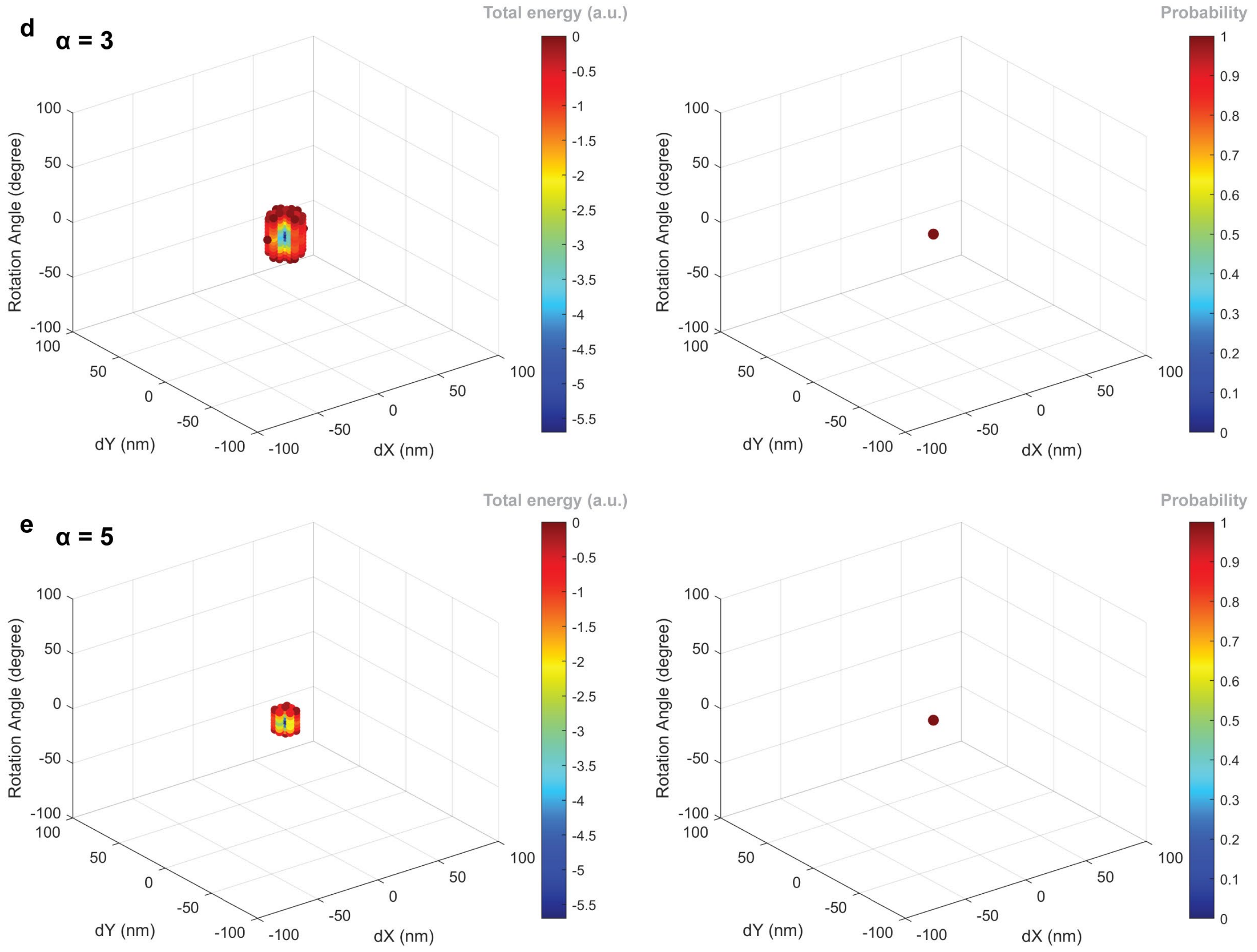


**Supplementary Fig. 20 | Total interaction free energy ensembles of triangle origami-well conformations with translational displacements (d$X$ and d$Y$) and rotational displacement ($\theta$) (a–e left) and the simulated probability of the final conformations (a–e right) for origami-well binding with increasing free energy penalty coefficient ($\alpha$).** Conformations with total interaction free energy $\geq 0$ were eliminated. Data points of conformations with both d$X \leq 0$ and d$Y \leq 0$ are omitted in total interaction free energy plots for better visualization (a–e left). Increasing free energy penalty coefficient ($\alpha$) correlates with increasing origami structural rigidity.

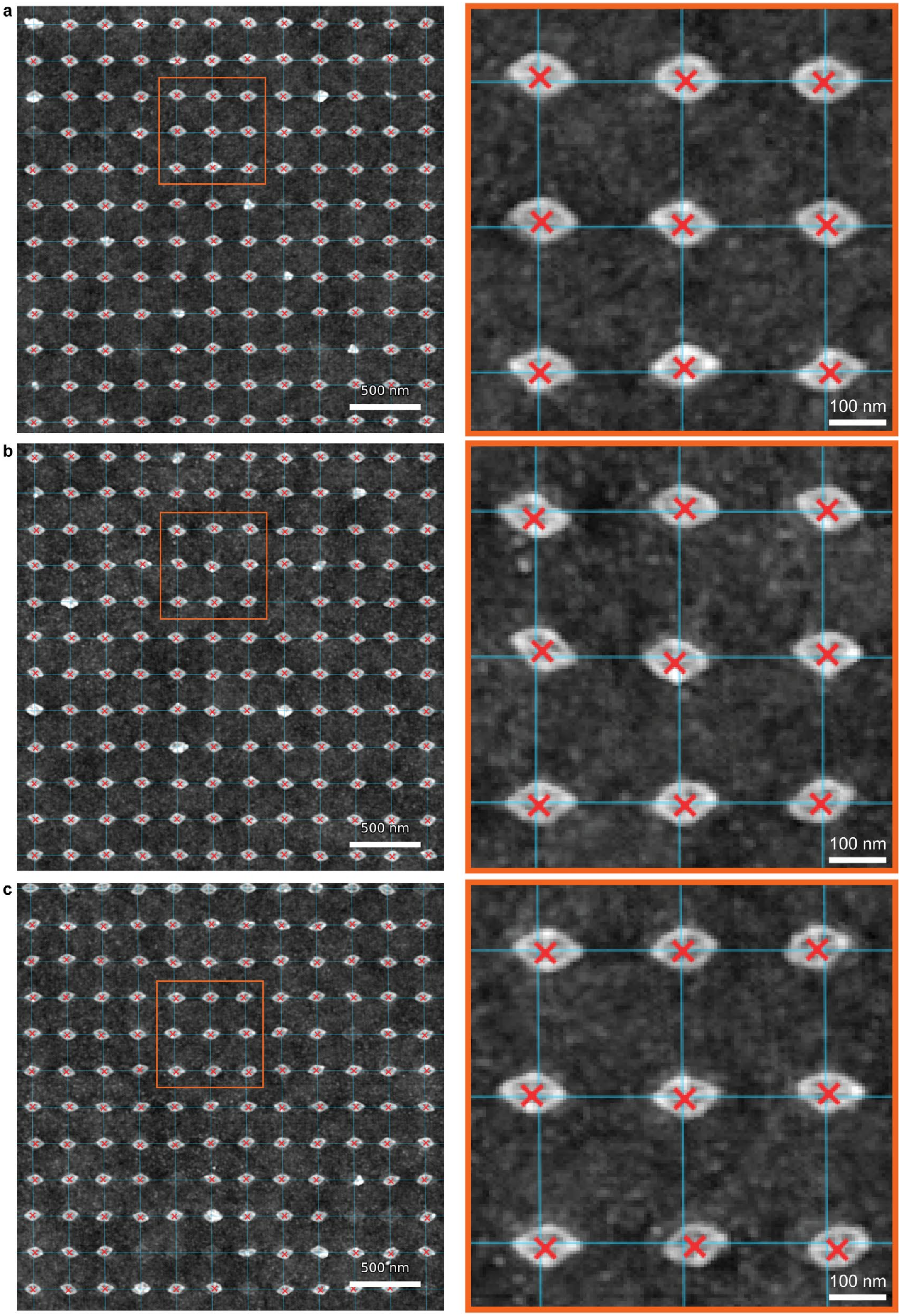


**Supplementary Fig. 21 | Placement precision of TMOP-directed rhombic origami for three well sizes. a–c**, Representative AFM images (left) and zoomed-in images (right) of rhombic origami arrays for well size 1 (a), size 2 (b) and size 3 (c). Cyan lines mark the reference lattice fitted to that image and red crosses the measured centroid of each correctly placed single origami. Displacement from the target lattice site grew with well size: mean values were 5.3 nm, 7.5 nm and 9.0 nm for sizes 1, 2 and 3 (medians 5.0 nm, 7.3 nm and 8.7 nm; 95th percentiles 9.9 nm, 13.6 nm and 17.0 nm), and the percentage of origamis within 10 nm of target fell from 94.9% to 76.2% and 59.1%. $N = 514$, 652 and 487 for sizes 1, 2 and 3, respectively. Displacement distributions are shown in Fig. 2f; image processing is described in Supplementary Note 9.

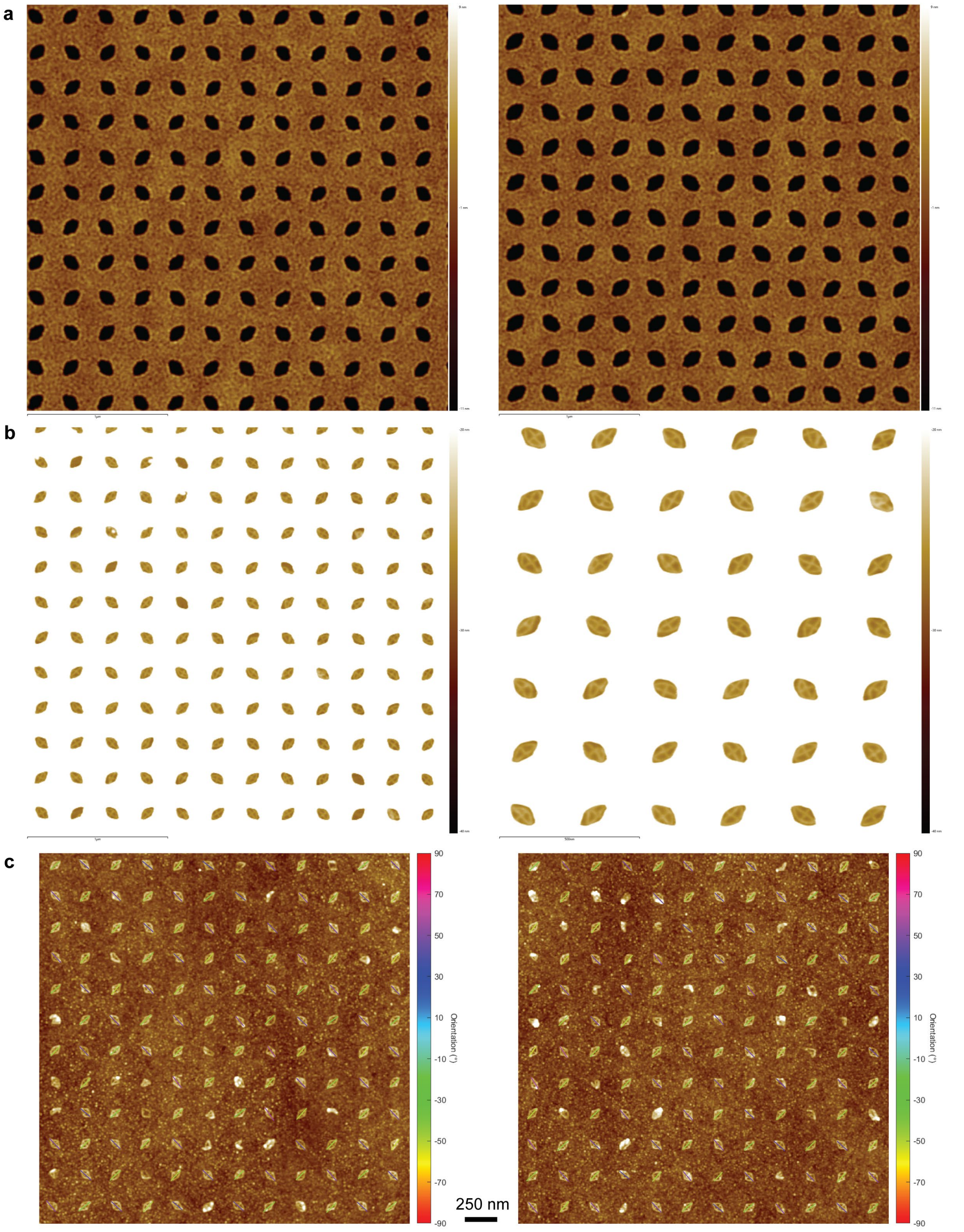


**Supplementary Fig. 22 | AFM images of TMOP with diagonally oriented (±45°) rhombic wells. a**, PMMA well arrays fabricated with 1,020 µC/cm$^2$ (size 1, left) and 1,320 µC/cm$^2$ (size 2, right) EBL doses**,** respectively. **b**, Rhombic origamis placed at the bottom of PMMA wells (size 2). **c**, Rhombic origami arrays after TMOP. No NaCl used for origami placement. Single origami orientations were measured as the angle of the long axis direction.

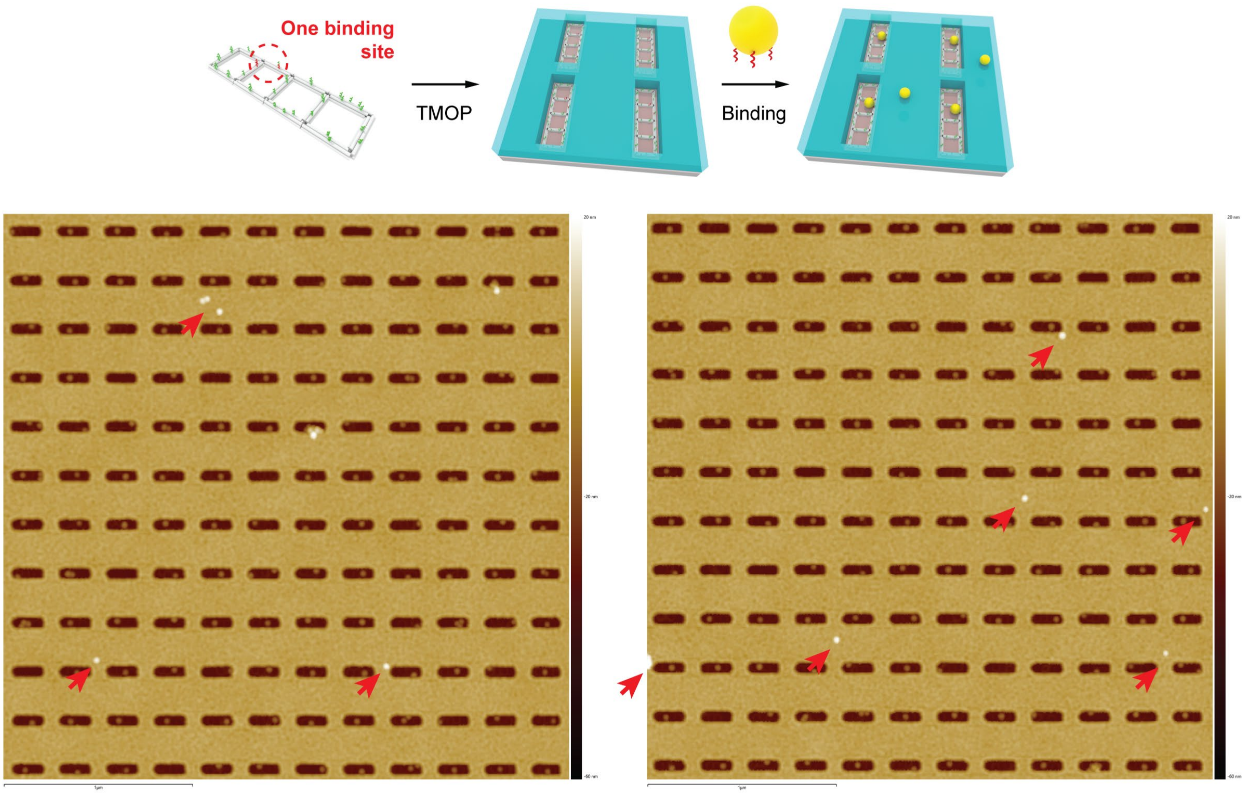


**Supplementary Fig. 23 | AFM images of TMOP-guided AuNS monomers placed within PMMA wells through ladder-shaped origami templating.** Arrows highlight AuNSs randomly landed on top of the PMMA resist layer.

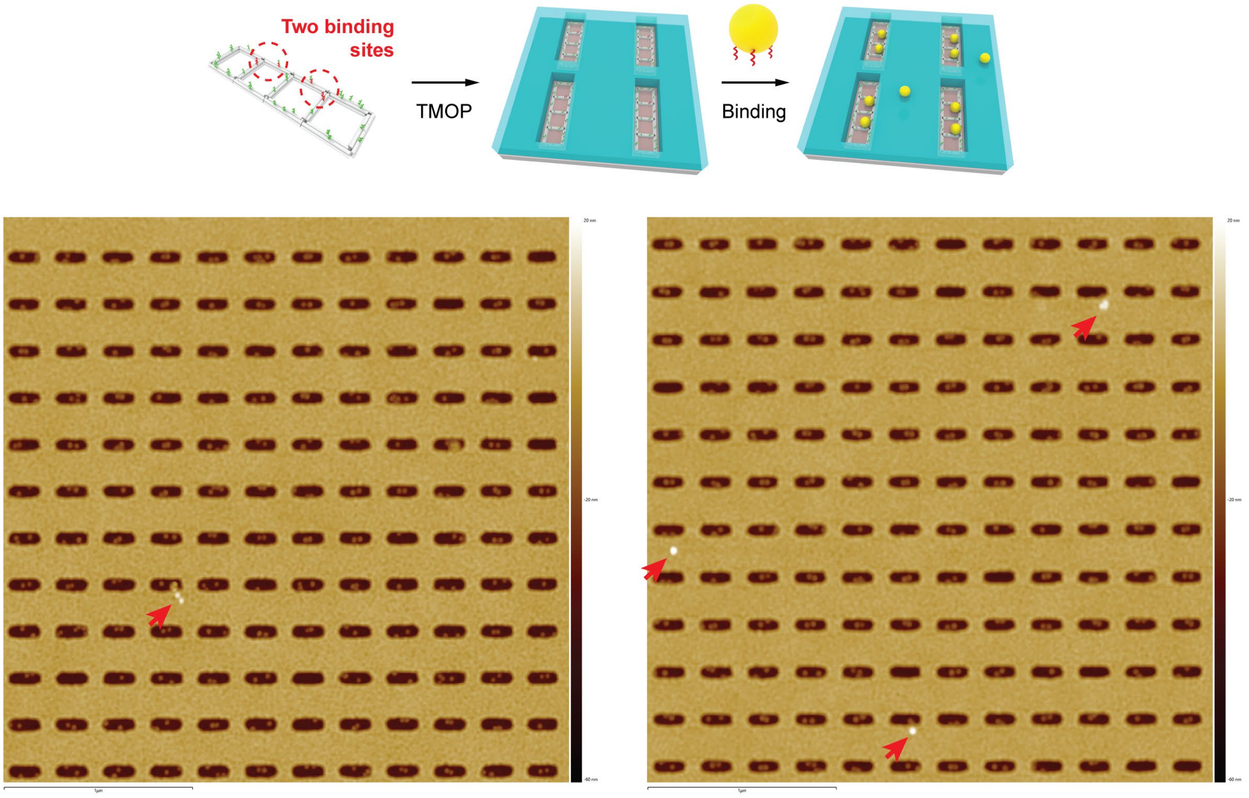


**Supplementary Fig. 24 | AFM images of TMOP-guided AuNS dimers placed within PMMA wells through ladder-shaped origami templating.** Arrows highlight AuNSs randomly landed on top of the PMMA resist layer.

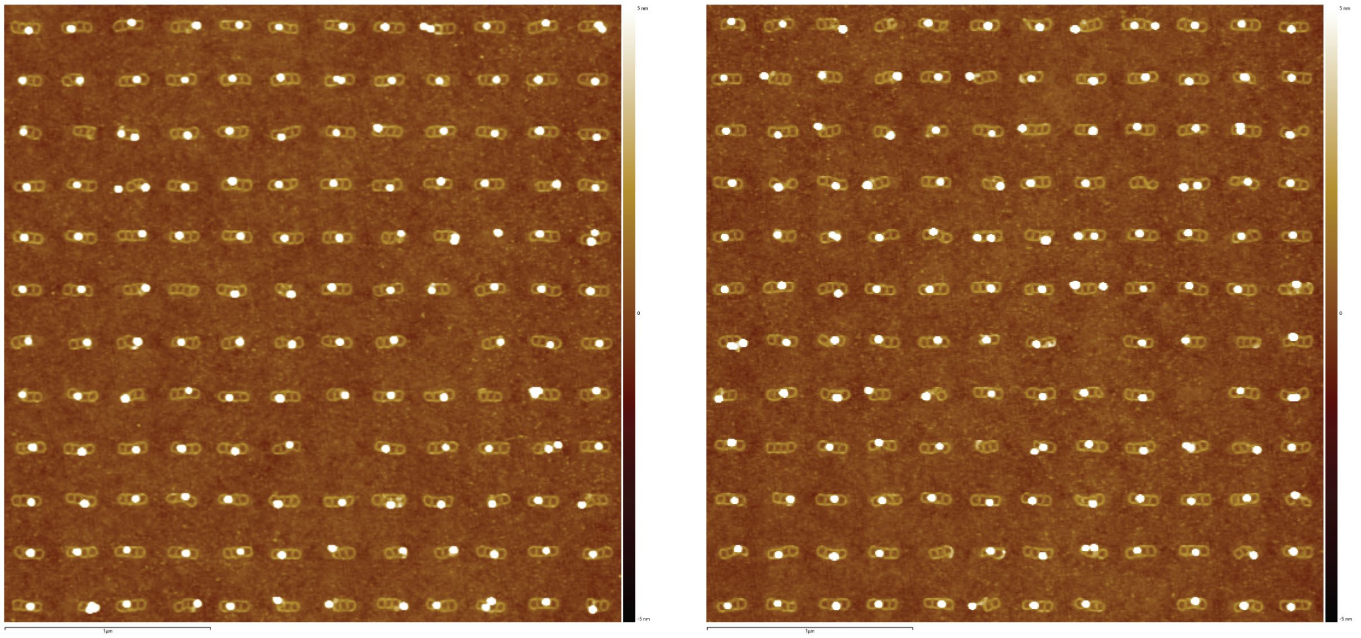

**Supplementary Fig. 25 | AFM images of TMOP-guided AuNS monomer arrays through ladder-shaped origami templating.**

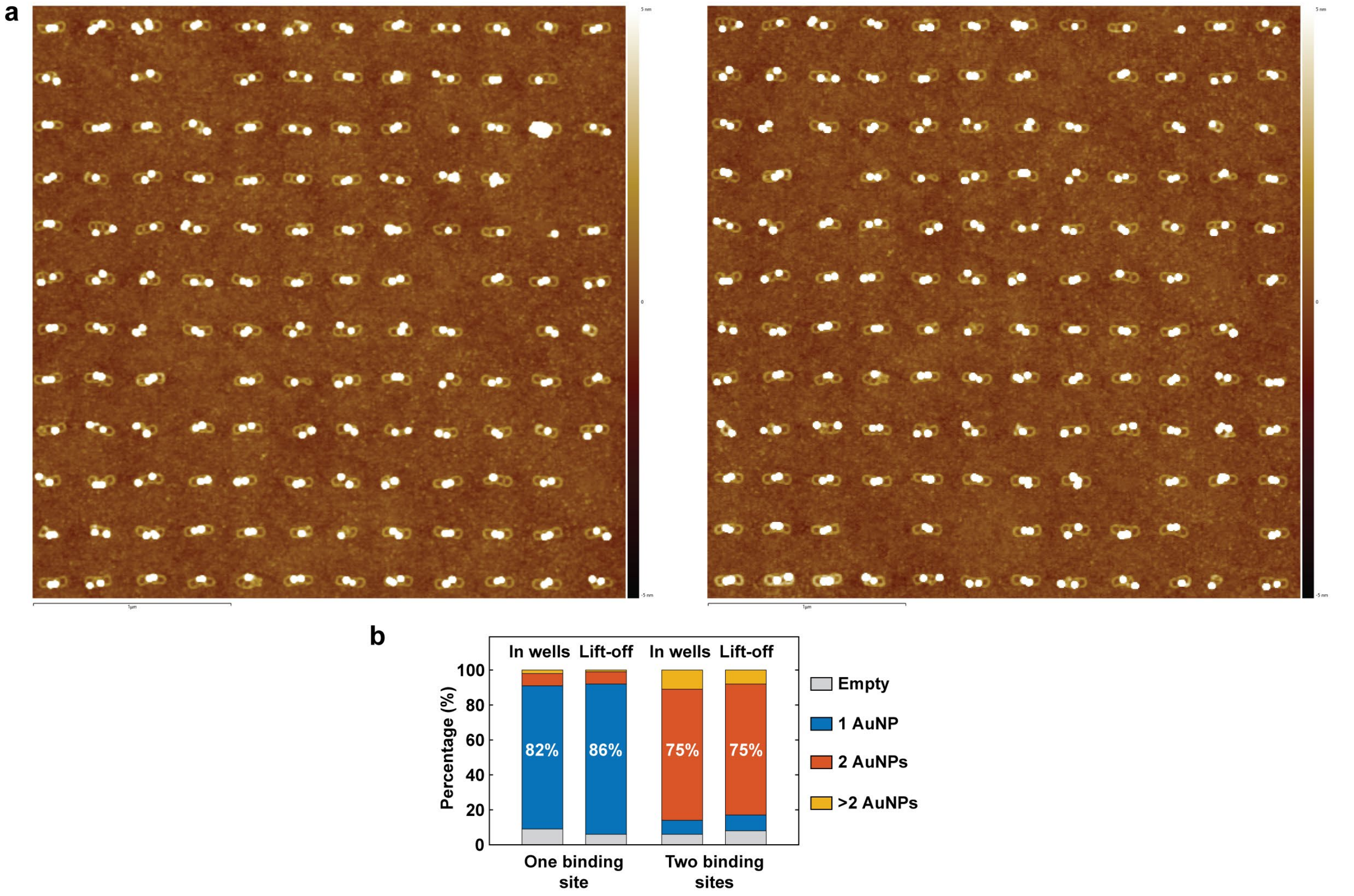


**Supplementary Fig. 26 | TMOP-guided AuNS dimers. a**, AFM images of TMOP-guided AuNS dimer arrays through ladder-shaped origami templating. **b**, Statistics of TMOP-guided AuNS arrangement before (in the well) and after lift-off using ladder-shaped origamis with 1 or 2 binding sites.

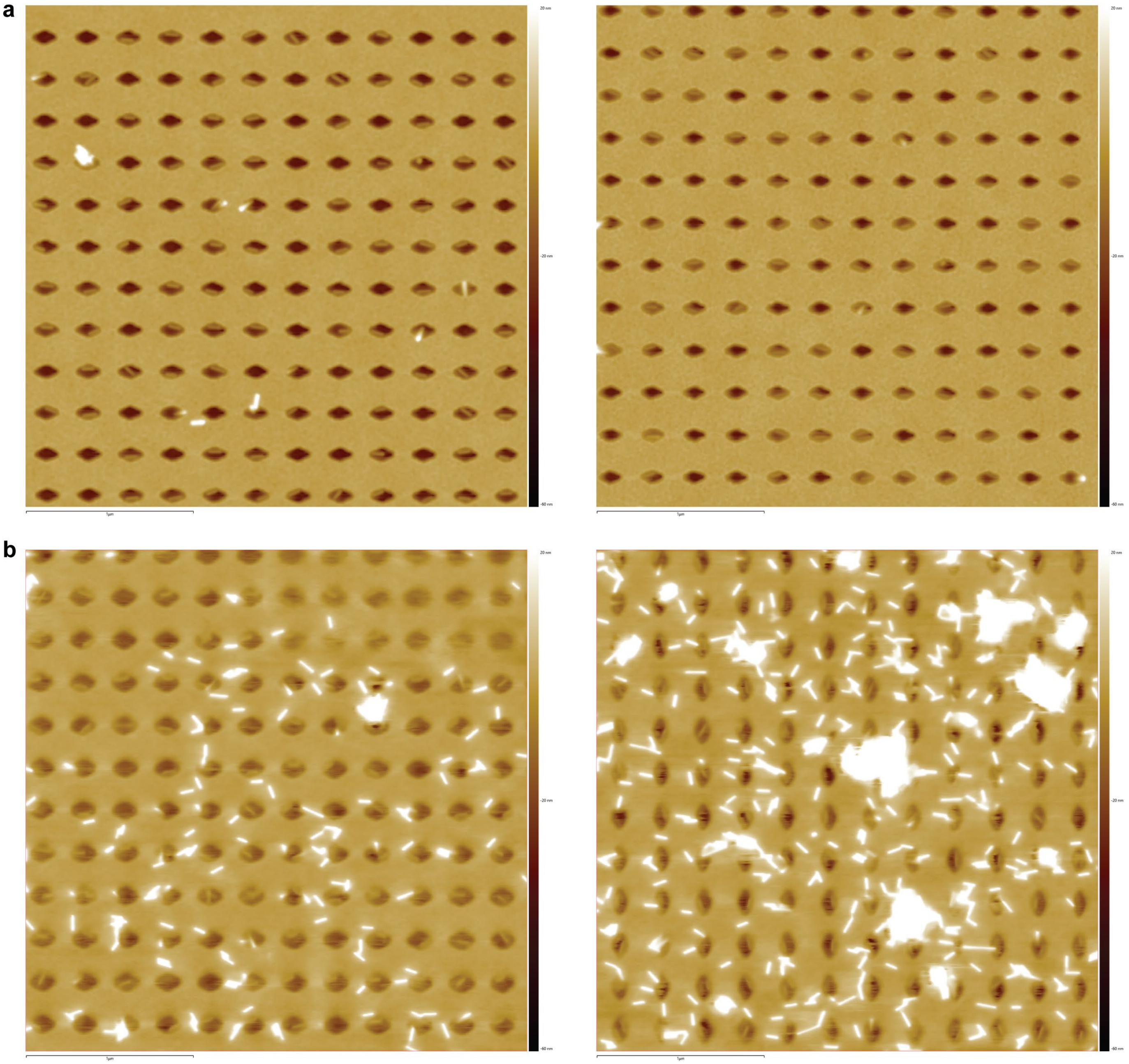


**Supplementary Fig. 27 | AFM images of TMOP-guided AuNRs placed within PMMA wells through rhombic origami templating. a**, AuNR solution incubated with well array for 1 h for AuNR binding. **b**, AuNR solution incubated with well array for 2.5 h for AuNR binding. Random AuNR depositions on top of the PMMA resist layer were clearly observed with longer incubation (binding) time.

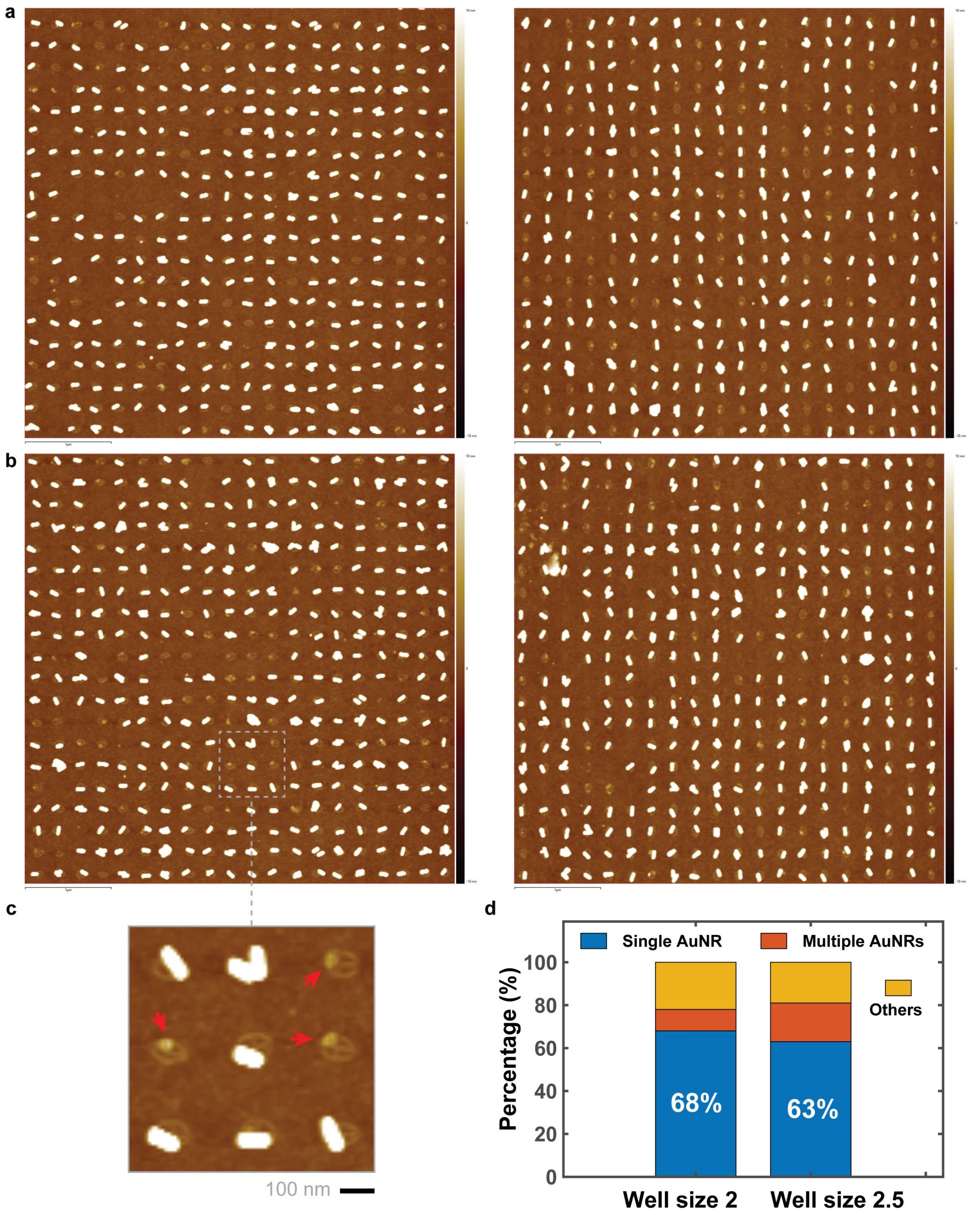


**Supplementary Fig. 28 | AFM images of TMOP-guided AuNR arrays with horizontal (a,b, left) or vertical (a,b, right) orientation arranged using well arrays fabricated with 1,320 µC/cm$^2$ (size 2, a) and 1,521 µC/cm$^2$ (size 2.5, b) EBL doses, respectively.** AuNR spacings are 250 nm. **c**, Zoomed-in image of AuNR array. Arrows highlight origami binding sites blocked by small NPs potentially from impurities from the purchased AuNR. **d**, Statistics of the AuNR placement with well arrays of different sizes. Samples processed with 2.5 h AuNR incubation (binding) time, which may increase random deposition of AuNRs.

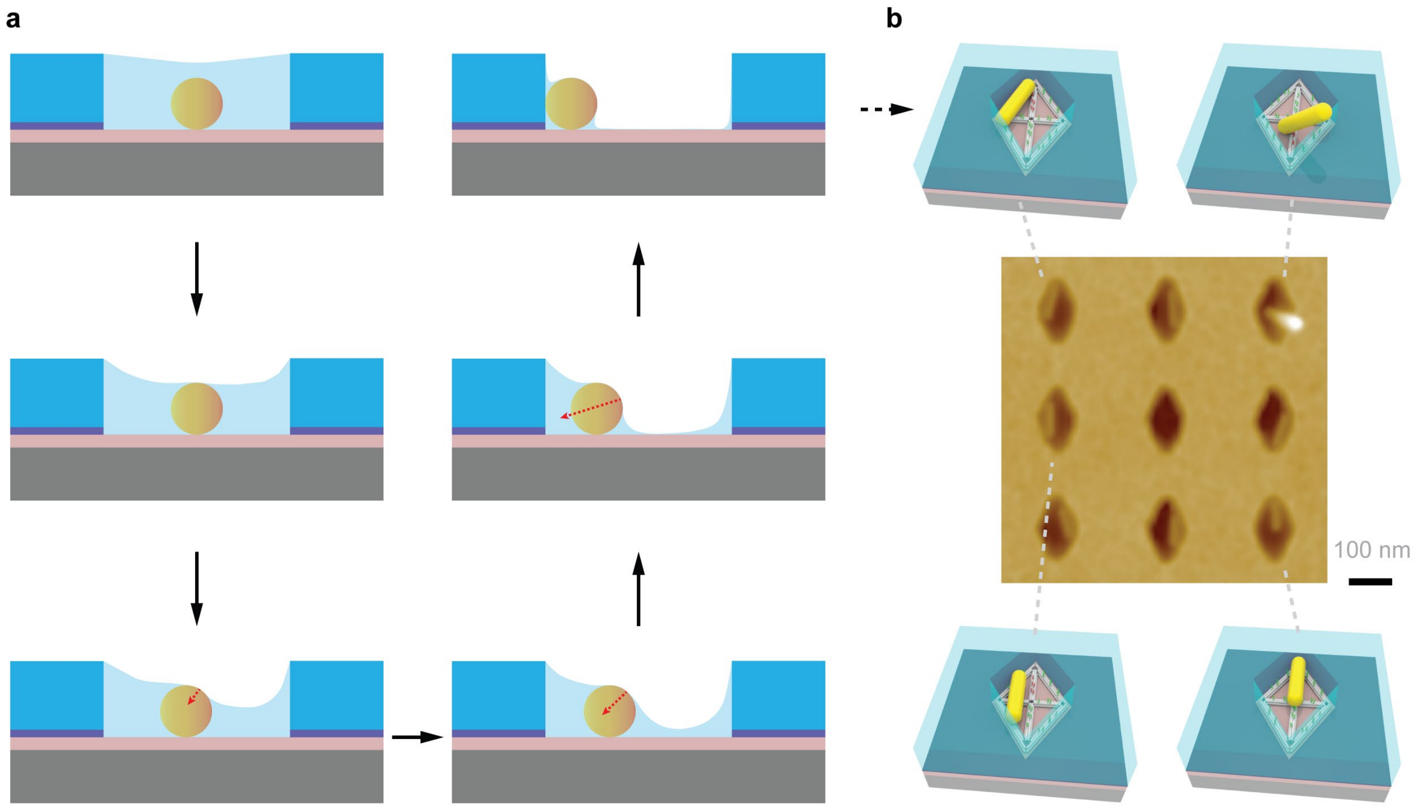


**Supplementary Fig. 29 | Schematic illustration of capillary-force-driven AuNR movement in the PMMA well. a**, As water evaporates from the PMMA well, the liquid meniscus recedes. As the liquid level decreases, asymmetric capillary forces guide the gold NP's motion and orientation through the well, which concludes with the NP's final positioning determined by the well geometry and the random drying of the water meniscus. **b**, AFM image and schematic illustration of AuNR conformations at the base of the PMMA well after drying.

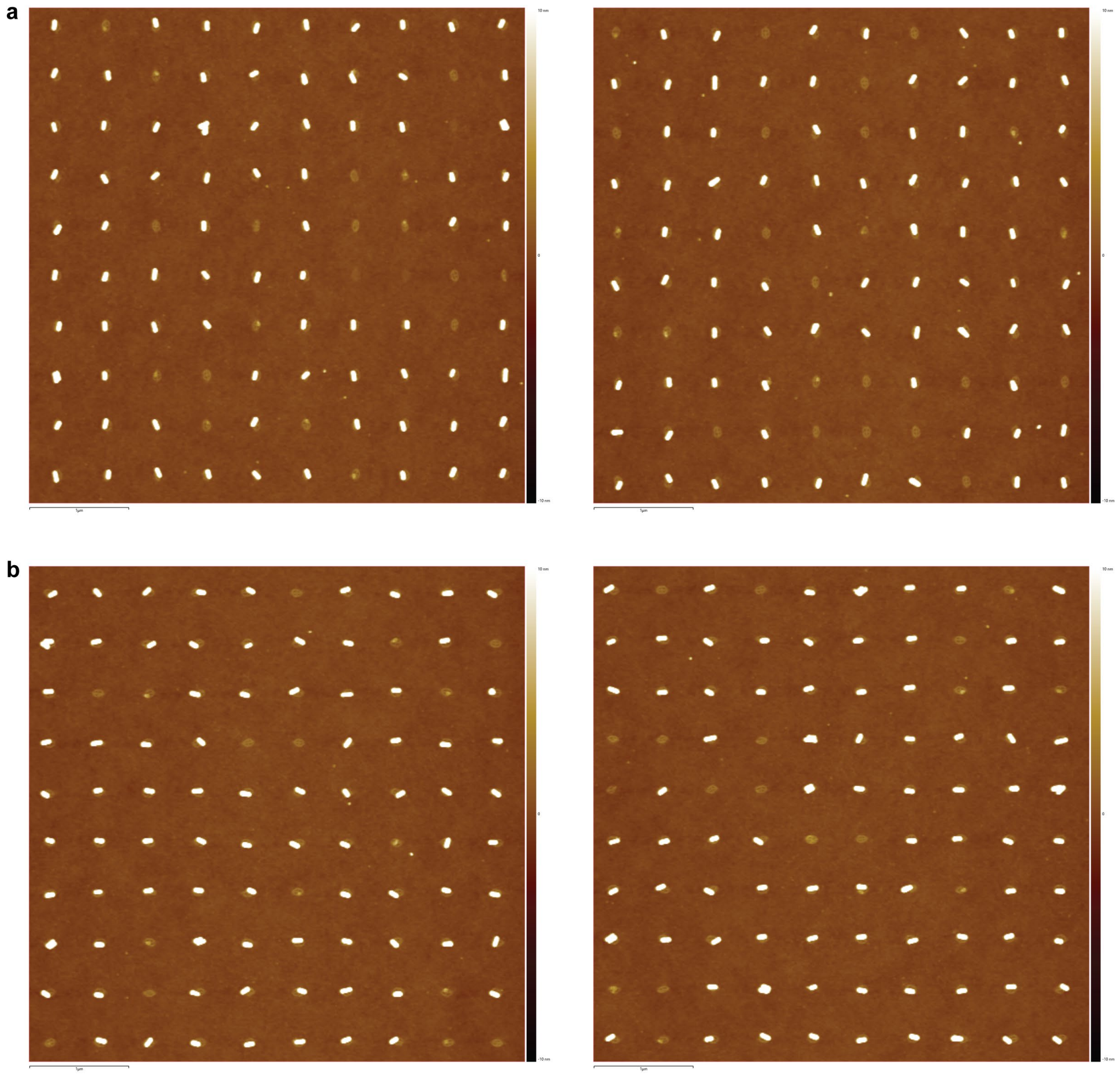


**Supplementary Fig. 30 | AFM images of TMOP-guided AuNR arrays with vertical (a) or horizontal (b) orientation arranged using well arrays fabricated with 1,521 μC/cm$^2$ (size 2.5) EBL dose.** AuNR spacings are 500 nm.

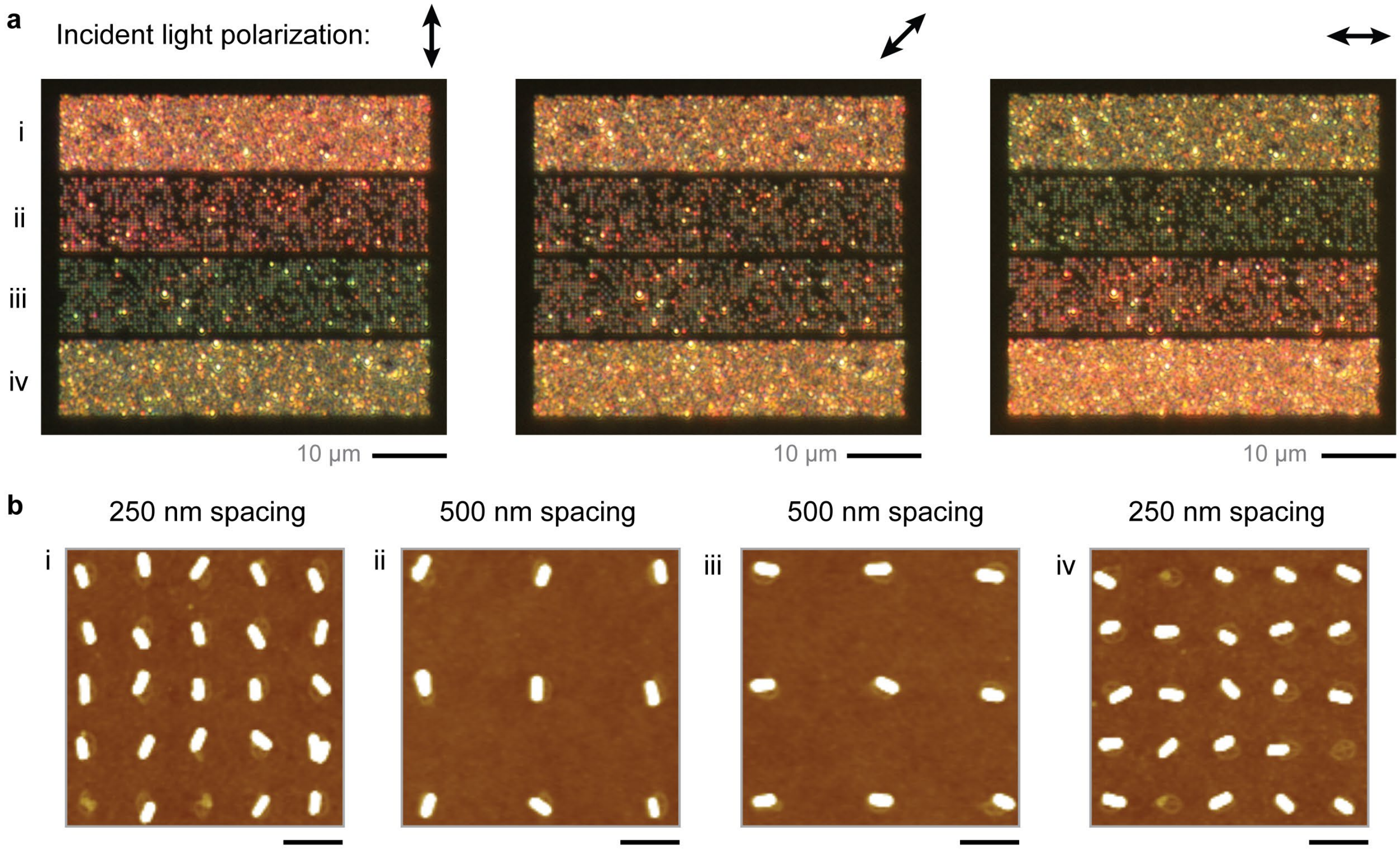


**Supplementary Fig. 31 | TMOP-directed AuNR array. a**, Dark-field microscopy images of the TMOP-directed AuNR array with incident light polarized vertically (left), diagonally (45°, middle), and horizontally (right), without an analyzer. **b**, AFM images of corresponding AuNR arrays with vertical (i, ii) or horizontal (iii, iv) orientations and 500 nm (i, iv) or 250 nm (ii, iii) pattern spacings.

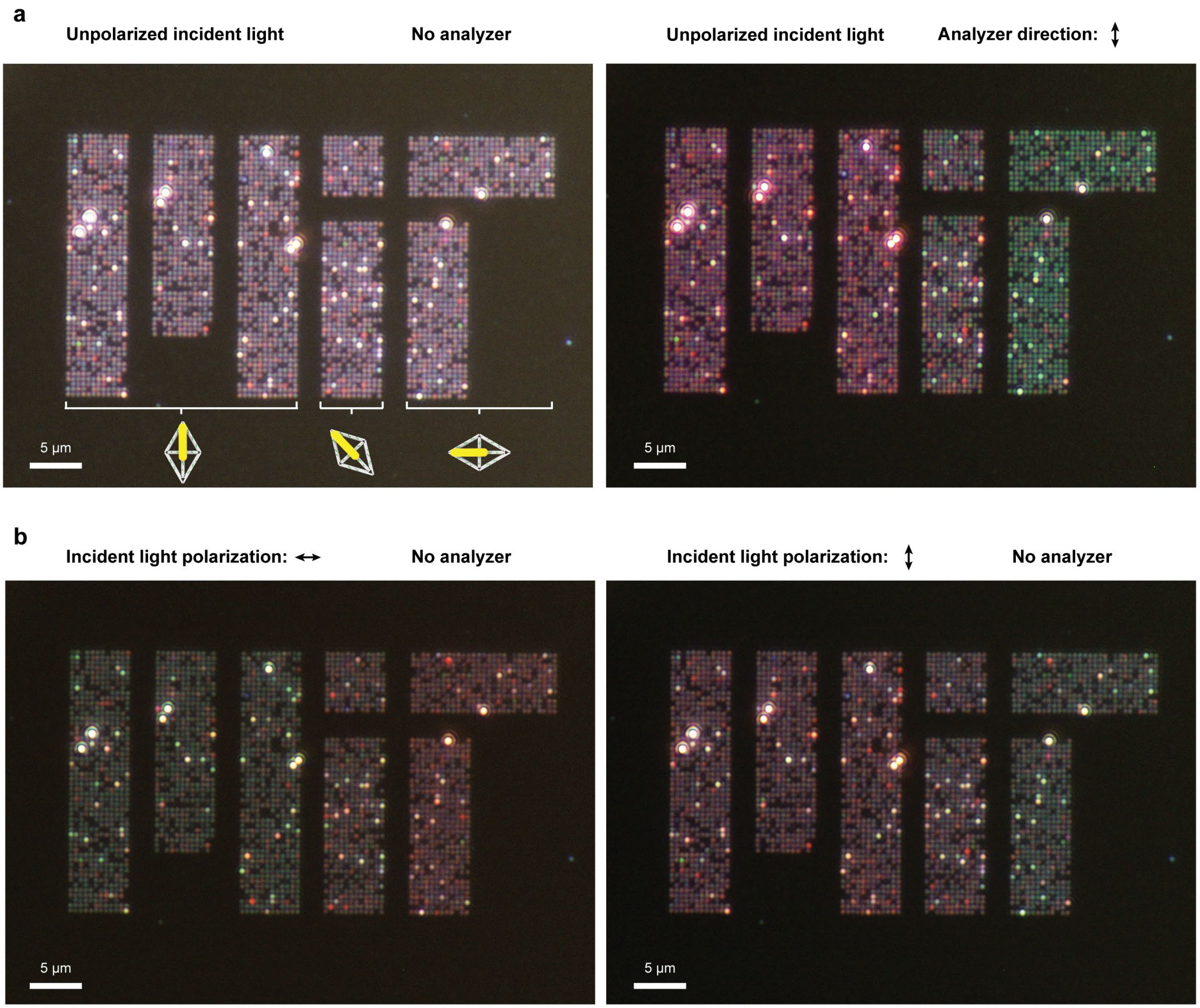


**Supplementary Fig. 32 | Dark-field microscopy images of TMOP-directed AuNR arrays in the shape of an MIT logo.** The letters "M", "I" and "T" were composed of AuNRs oriented perpendicularly, with a 45° angle, or horizontally, respectively. **a**, Dark-field microscopy images with unpolarized incident light and no analyzer (left) or with unpolarized incident light and a perpendicularly aligned analyzer (right). **b**, Dark-field microscopy images with horizontally polarized incident light and no analyzer (left) or perpendicularly polarized incident light and no analyzer (right). AuNR spacings are 500 nm.

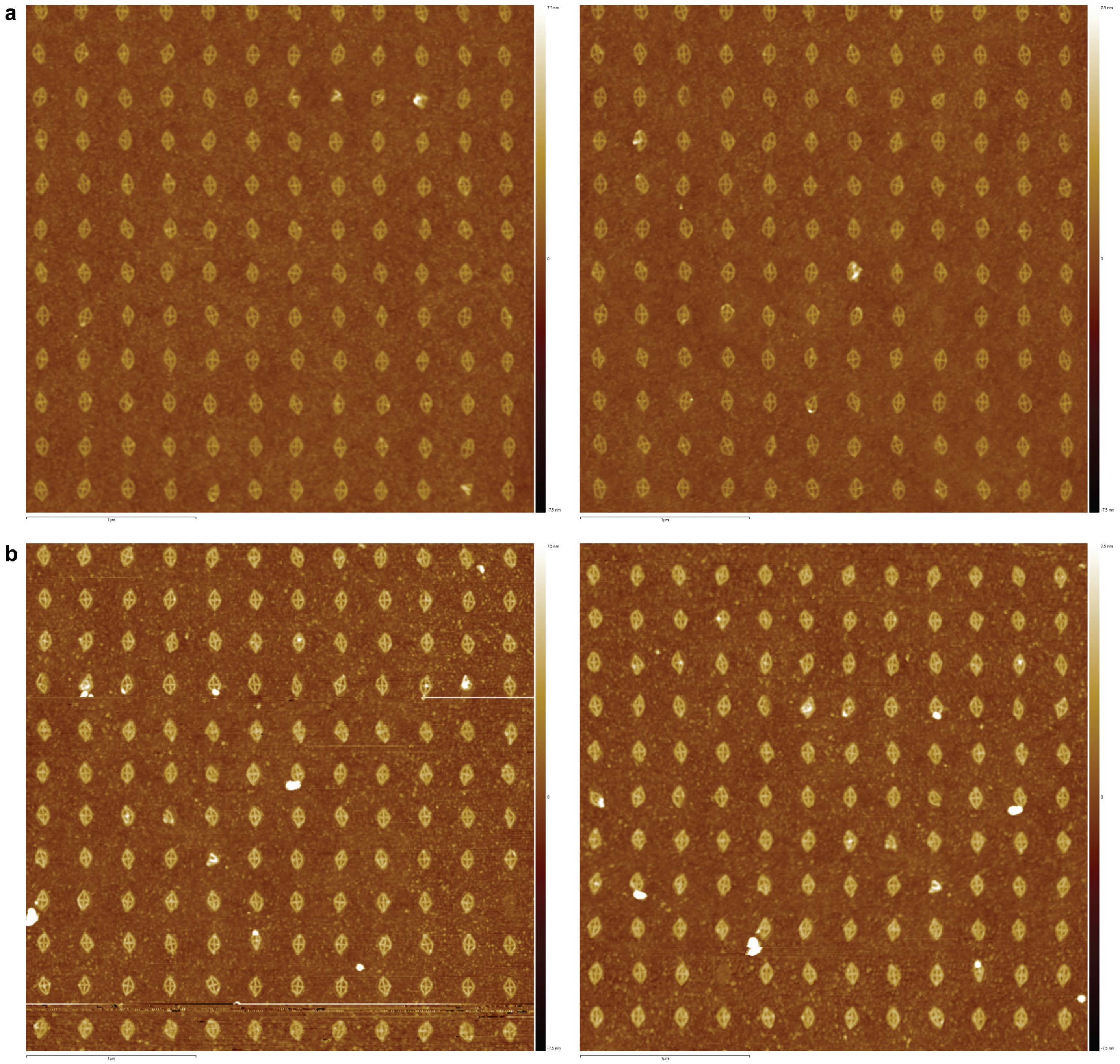

**Supplementary Fig. 33 | AFM images of TMOP rhombic origami arrays after soft-silicification of 1.5 h (a) and 22 h (b).** AFM image height scales are the same.

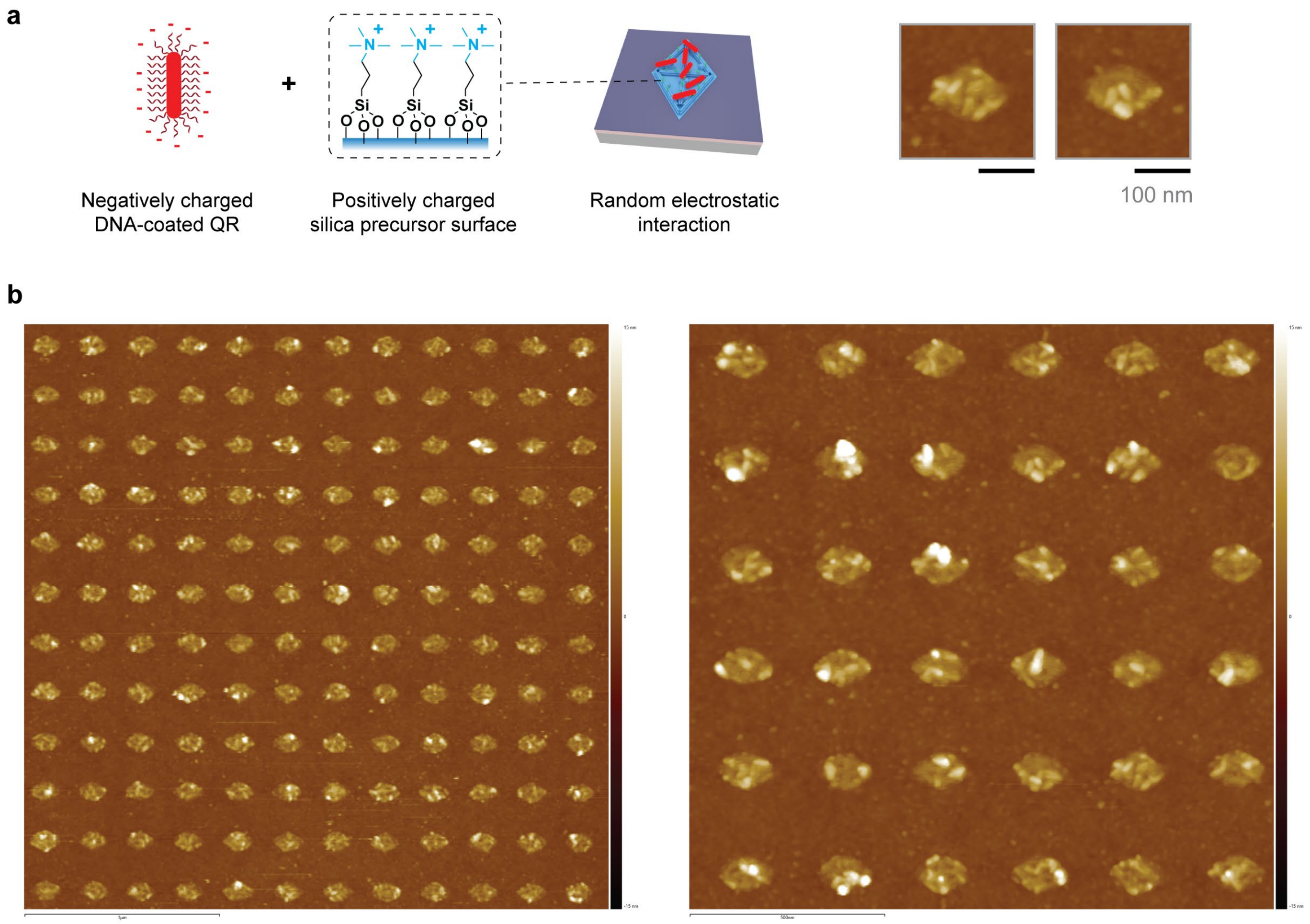


**Supplementary Fig. 34 | Random attachment of DNA-functionalized QRs onto silicified origamis. a**, Illustration and AFM images of random QR attachment onto silicified TMOP origami (without passivation) via random electrostatic interactions. **b**, AFM images of random QR attachment onto TMAPS silicified (22 h) TMOP origamis.

a

b

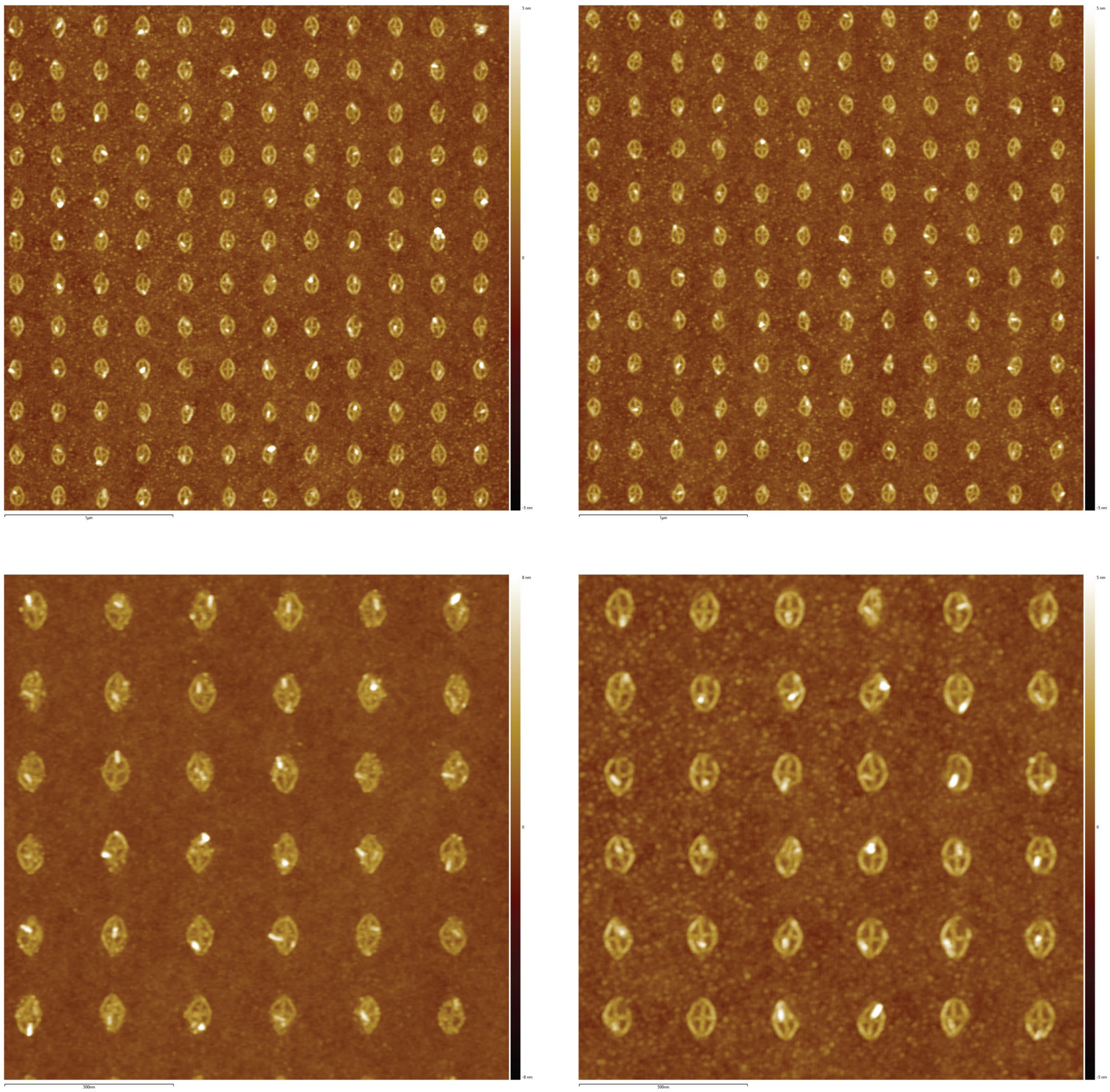

**Supplementary Fig. 35 | AFM images of TMOP-directed individual QRs binding to vertically aligned rhombic origamis. a**, 3 µm AFM scan. **b**, 1.5 µm AFM scan.

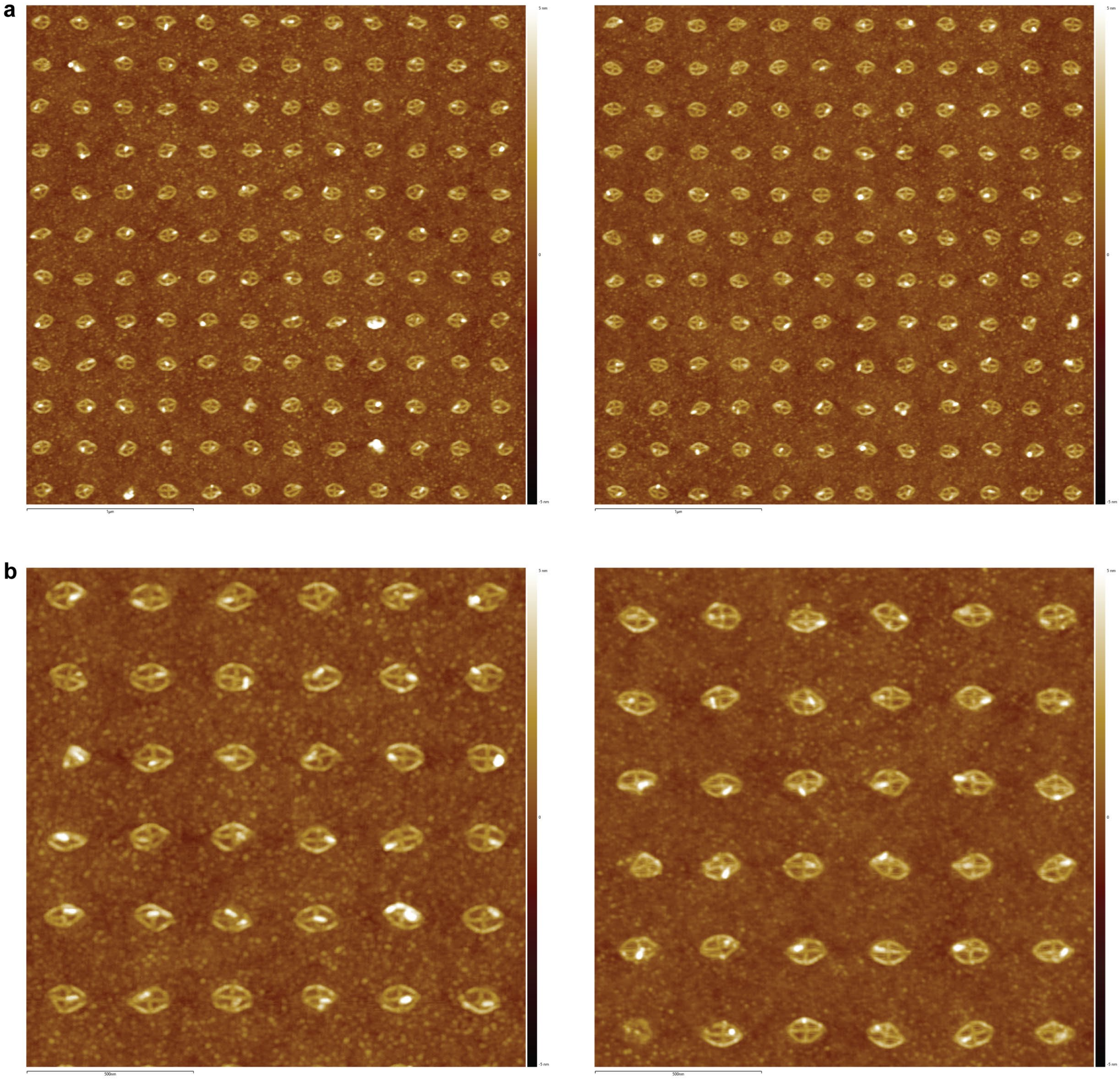

**Supplementary Fig. 36 | AFM images of TMOP-directed individual QRs binding to horizontally aligned rhombic origamis. a**, 3 µm AFM scan. **b**, 1.5 µm AFM scan.

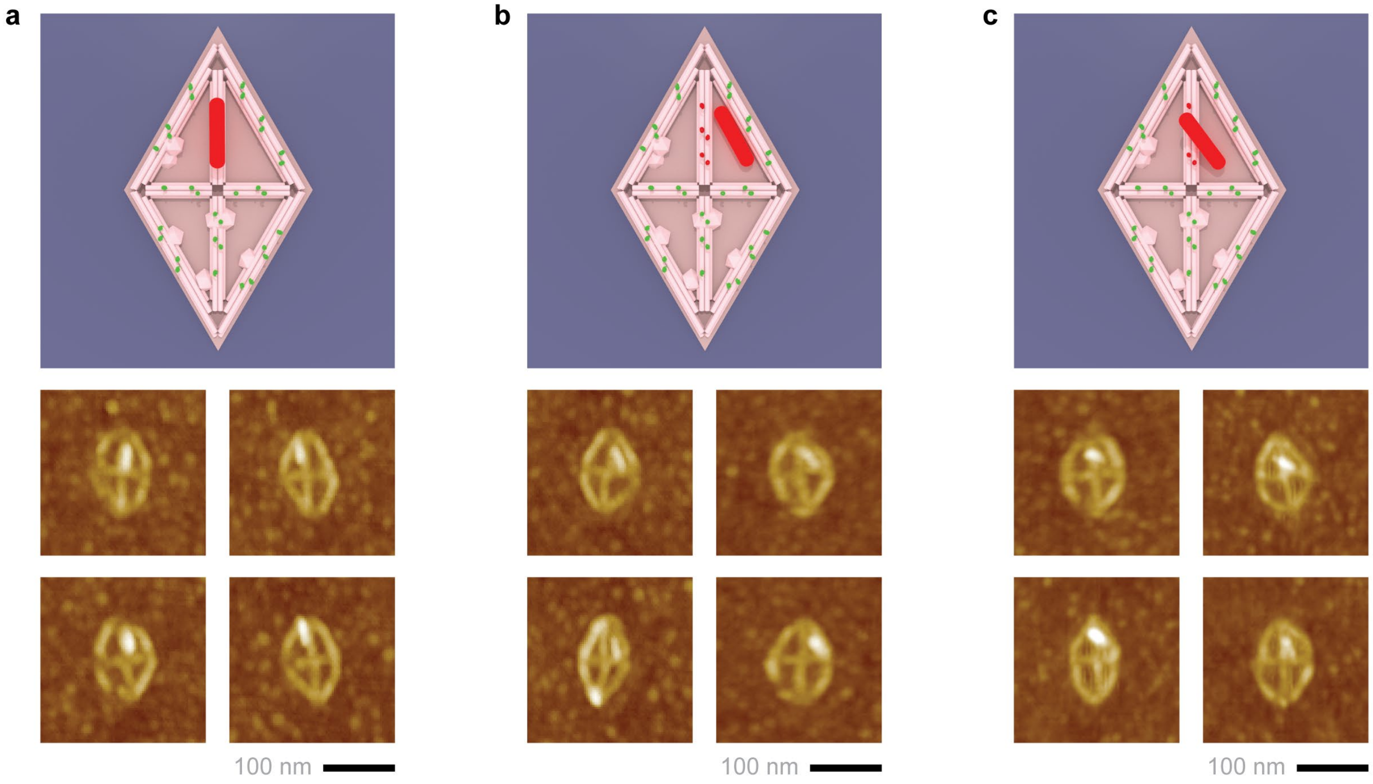


**Supplementary Fig. 37 | Schematic illustration and AFM images of QR alignment (a) on TMOP origami templates, and misalignment (b,c) potentially caused by capillary-force-driven reconfiguration into the topography of the soft-silicified wireframe 6HB origami structure.** Due to the comparable heights of the 6HB origami (*ca.* 5.5 nm in solution) and the QR employed (5 ± 0.6 nm), we speculate that QRs bound to the binding site could likely be pulled partially or entirely into neighboring triangular holes within the wireframe origami structure by the drying water meniscus within the well. If correct, then orientation control could potentially be improved through rational design of origami hole geometry and DNA binding overhang positions.

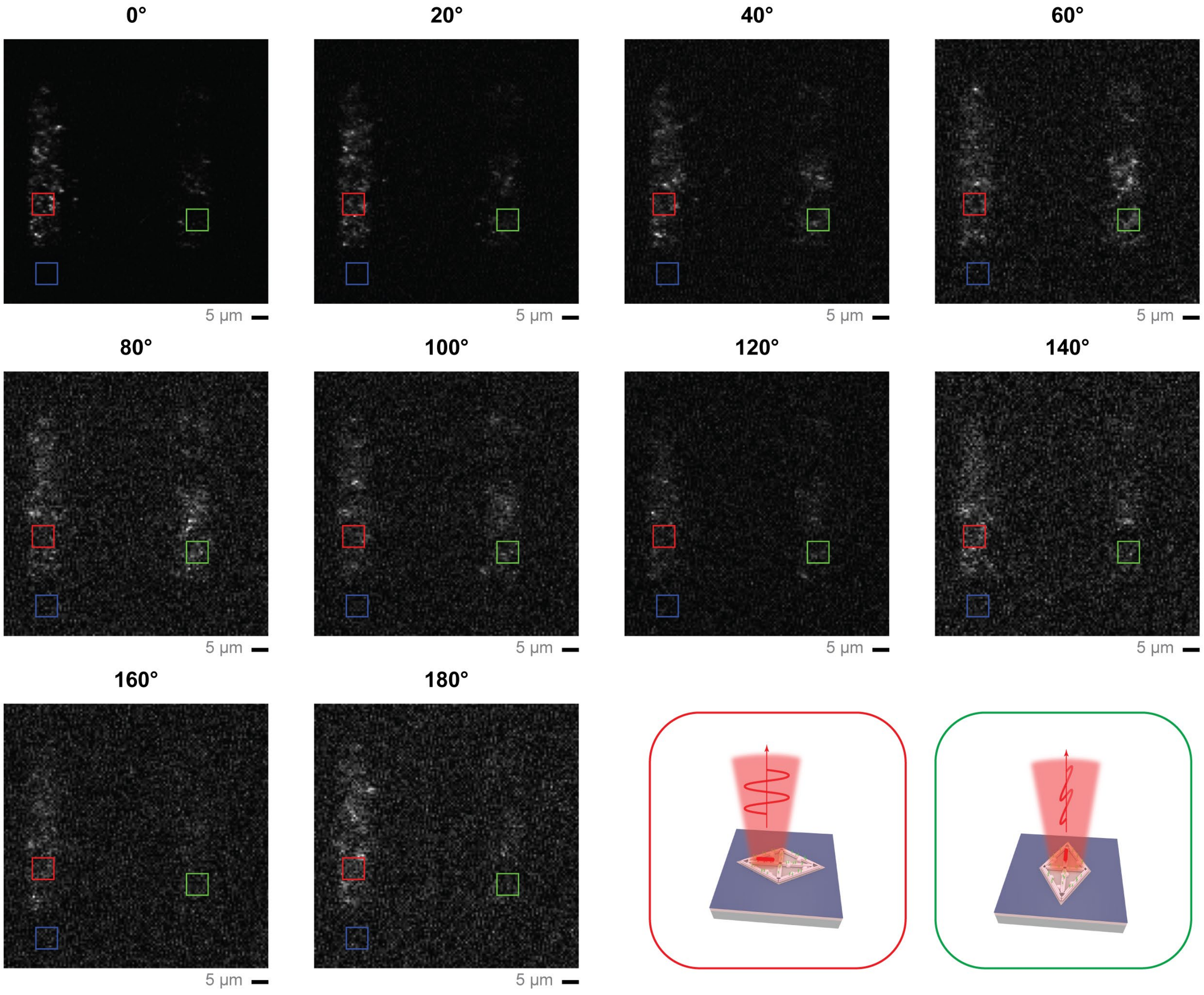


**Supplementary Fig. 38 | Example raster scanning confocal fluorescence images of the TMOP QR array collected through an analyzer with an angle from 0° to 180°.** Six images were collected at each angle; areas on QR arrays with orthogonal orientations (red and green boxes) were selected to extract the average fluorescence intensity and background-corrected against an empty area (blue box). The net intensities were then averaged over the six images for each angle and plotted in Fig. 4h. Complete images collected are included in the online data repository.

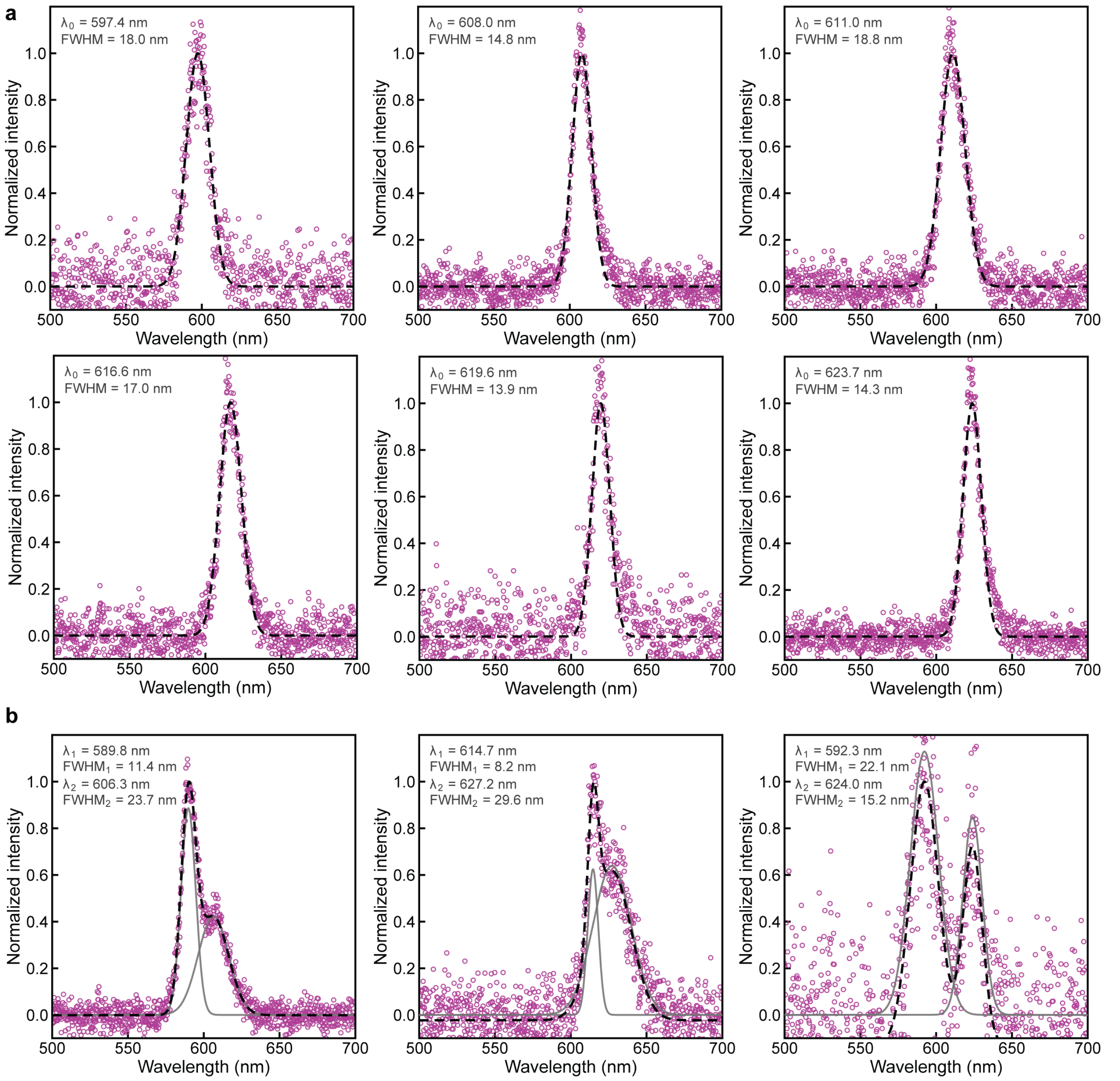


**Supplementary Fig. 39 | Additional examples of PL spectra collected from TMOP-directed QR arrays. a**, PL spectra with single peaks. **b**, PL spectra with multiple peaks, attributed to occasional multiple QRs placed at the same origami location.

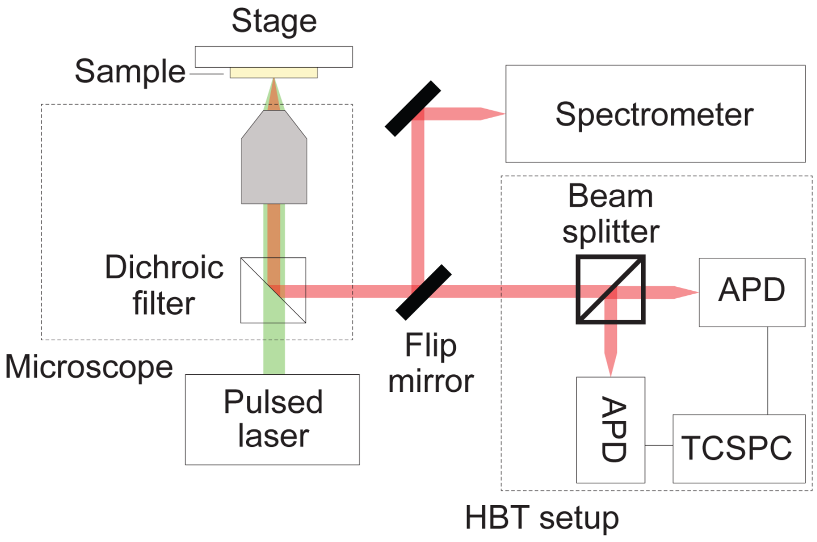


**Supplementary Fig. 40 | Optical setup for second-order photon correlation and PL spectrum measurements.** Top arm sends the emitted light to a spectrometer setup, consisting of a monochromator and a thermoelectrically cooled CCD. Bottom arm sends light to a Hanbury Brown–Twiss setup, in which the photons pass through a beamsplitter to two avalanche photodiodes, whose counts are then correlated by a time-correlated single-photon counting (TCSPC) module.

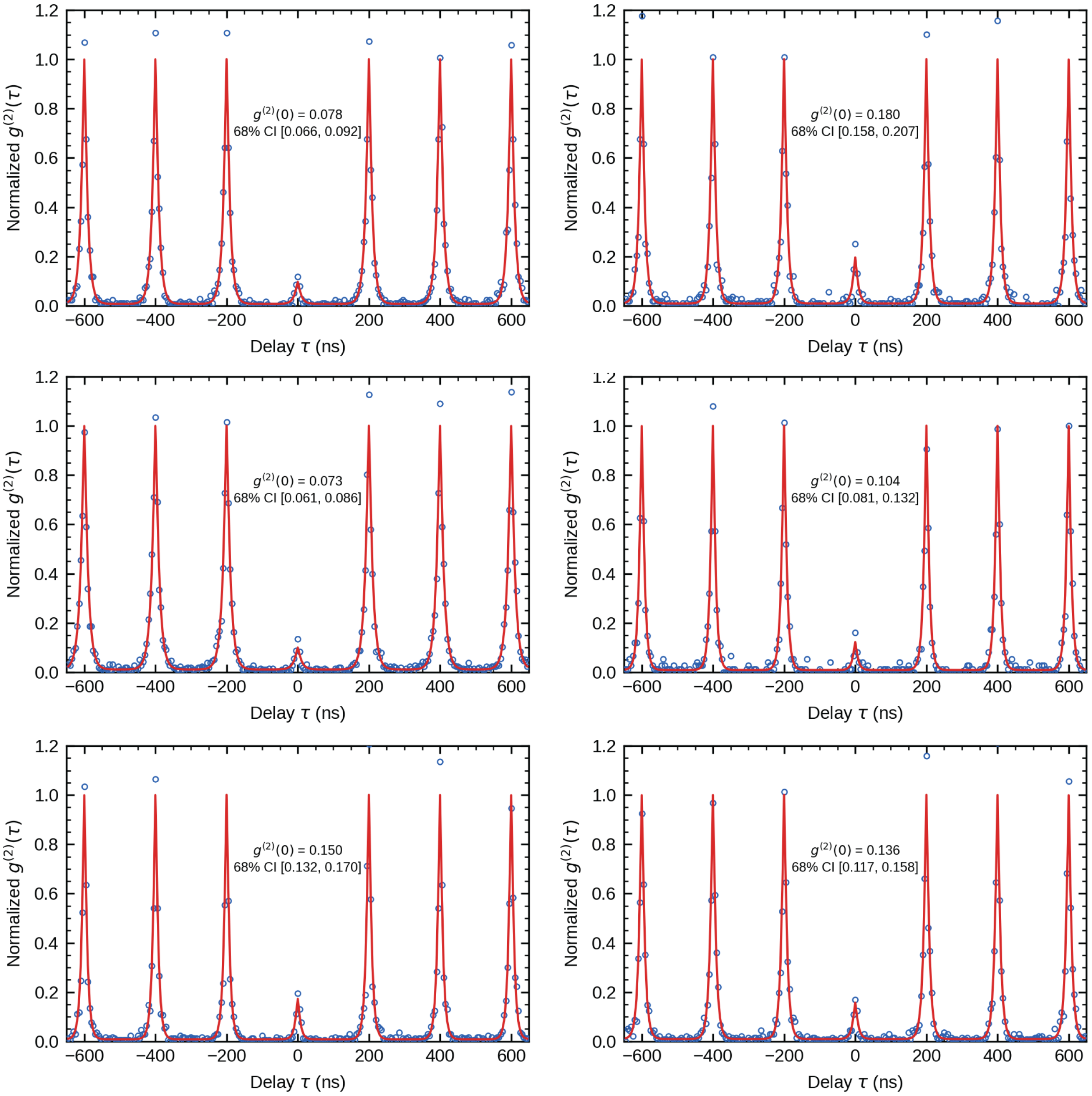


**Supplementary Fig. 41 | Additional examples of second-order correlation traces, $g^{(2)}(\tau)$, collected from TMOP-directed QR arrays.** $g^{(2)}(0)$ values calculated from raw peak areas (not fitted) as described in Supplementary Note 6. Brackets: 68% credible interval.

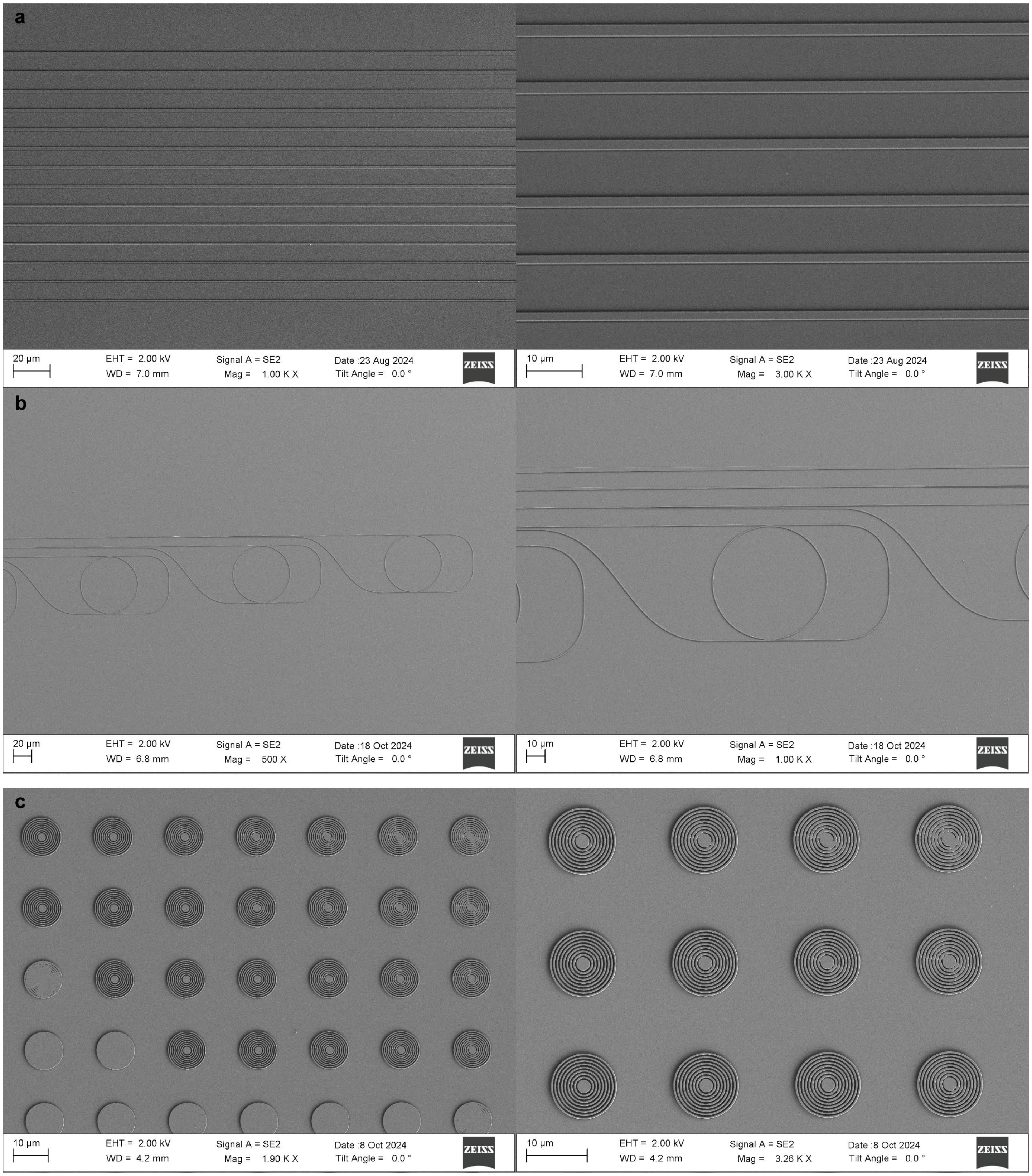


**Supplementary Fig. 42 | SEM images of $SiN_x$ photonic structures (310 nm high). a**, Slab waveguides of various widths (2 µm to 0.5 µm). **b**, Micro-ring resonators. **c**, Bullseye cavities of varied periodicity and duty cycle.

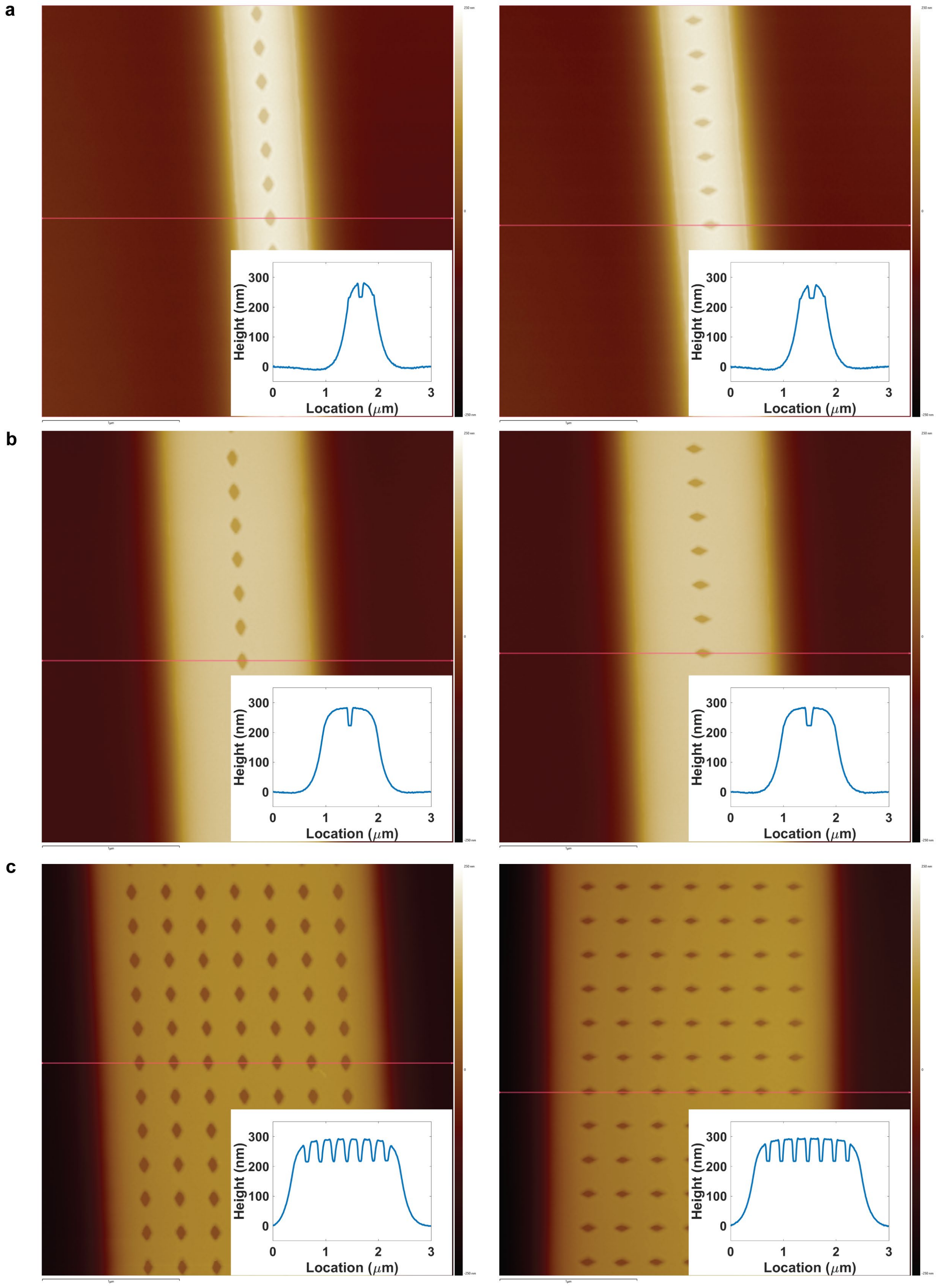


**Supplementary Fig. 43 | AFM images of PMMA wells fabricated on $SiN_x$ waveguides of 0.5 µm (a), 1 µm (b) and 2 µm (c) width.** Insets show height profiles along the red lines in each image. Well depth (PMMA thickness on waveguides) was higher on wider waveguides.

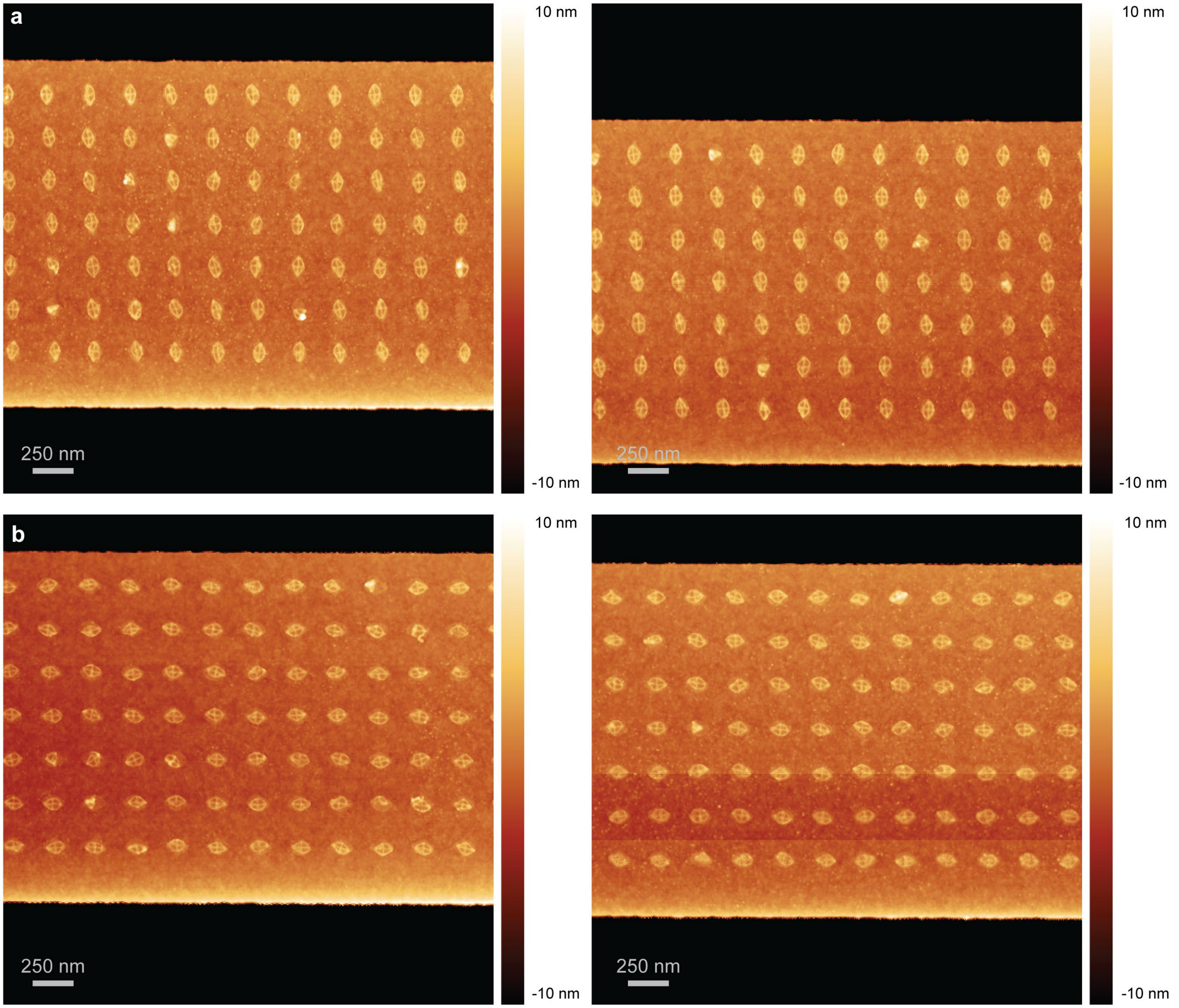


**Supplementary Fig. 44 | AFM images of TMOP rhombic origami arrays (soft-silicified) with orthogonal (a) or parallel (b) orientations on 2 µm $SiN_x$ waveguides.**

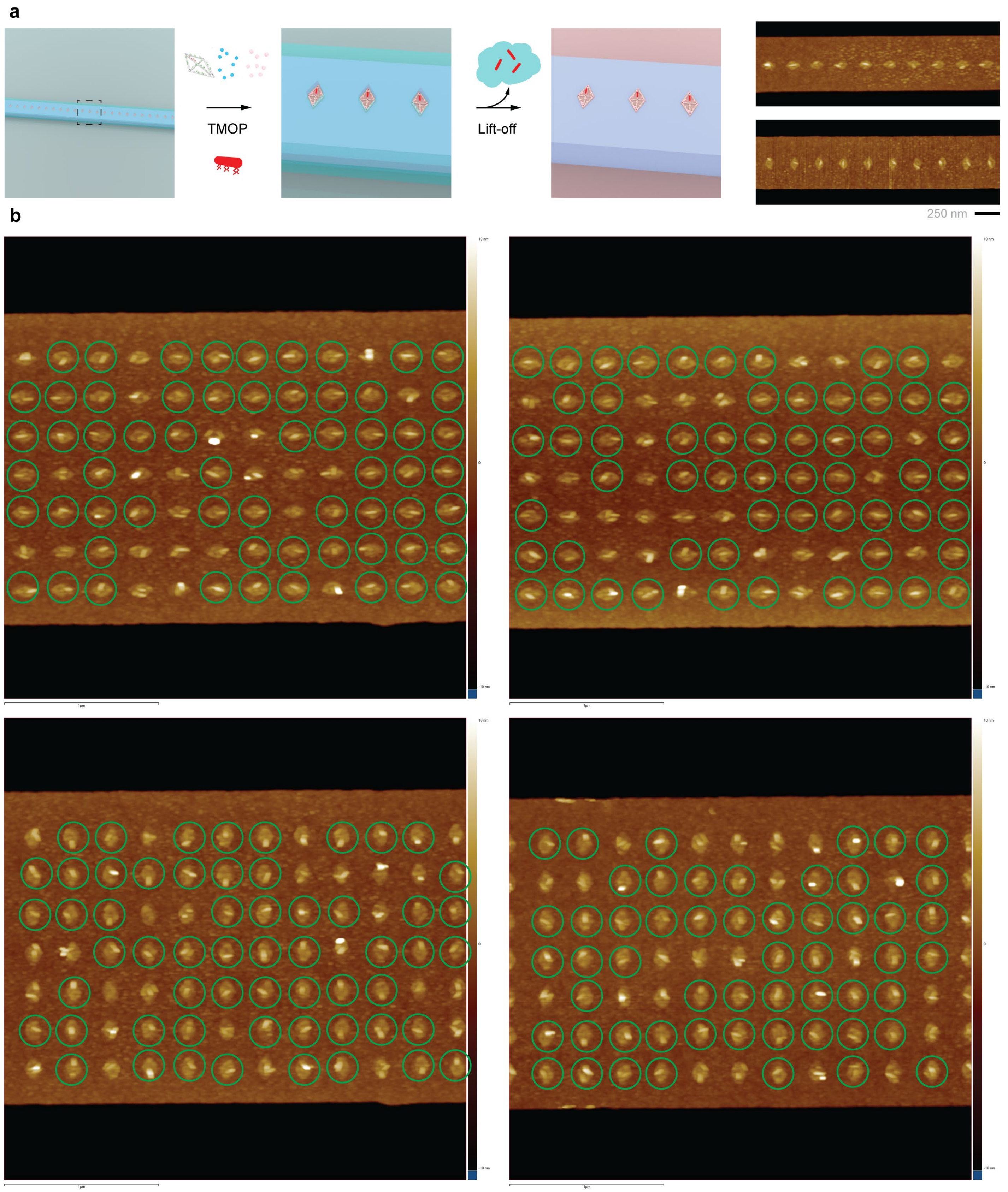


**Supplementary Fig. 45 | TMOP-directed QR placement on $SiN_x$ slab waveguides. a**, Schematic of TMOP-directed QR placement and resist lift-off on a $SiN_x$ waveguide (trimethylsilyl coatings omitted), and AFM images of orthogonal and parallel QRs on 0.5 µm slab waveguides. **b**, AFM images of TMOP rhombic origami arrays with QRs on 2 µm $SiN_x$ waveguides. Green circles mark origamis with a single QR.

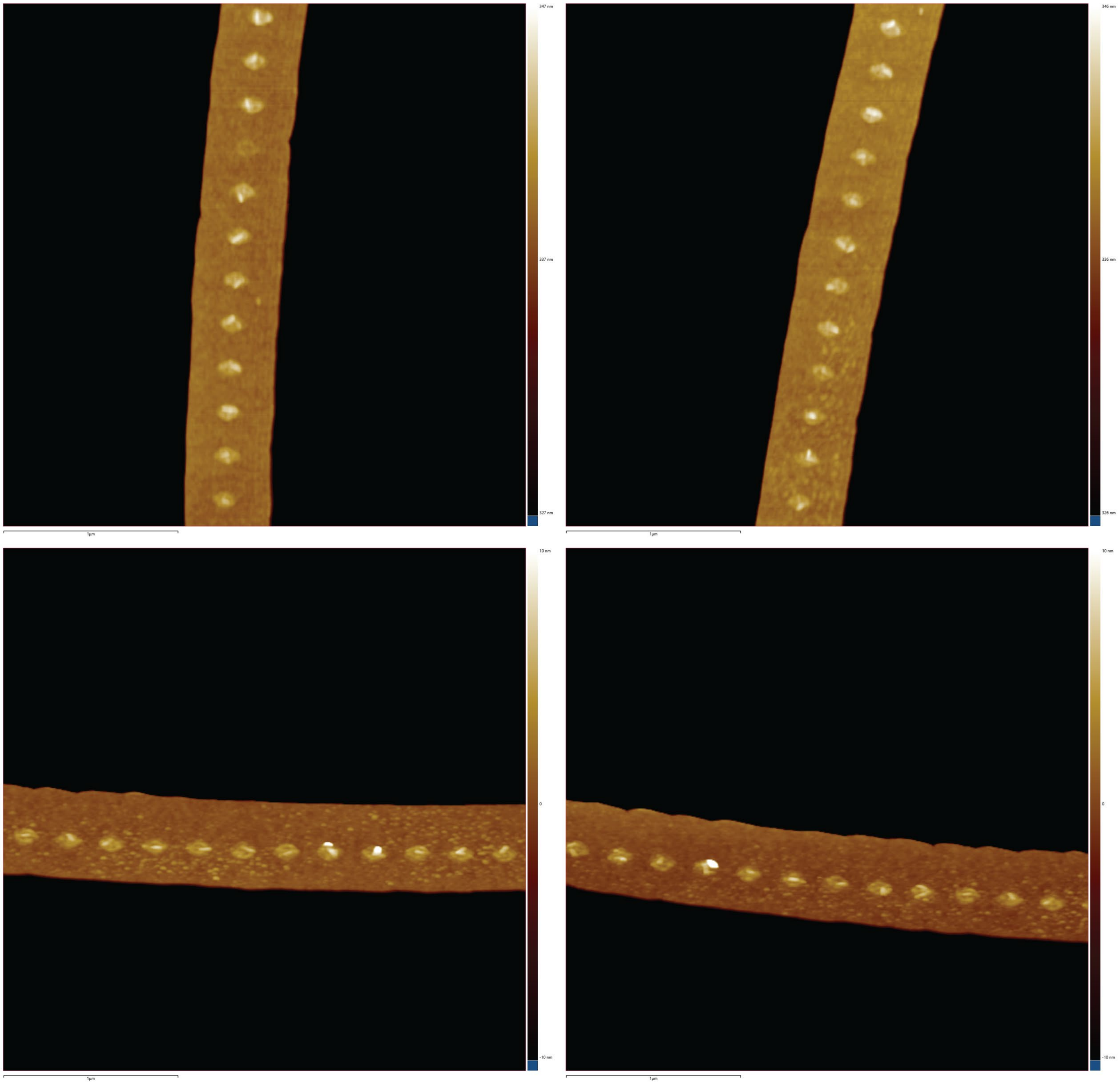


**Supplementary Fig. 46 | AFM images of TMOP-directed QRs with orthogonal or parallel orientations on SiN$_x$ micro-ring resonators.**

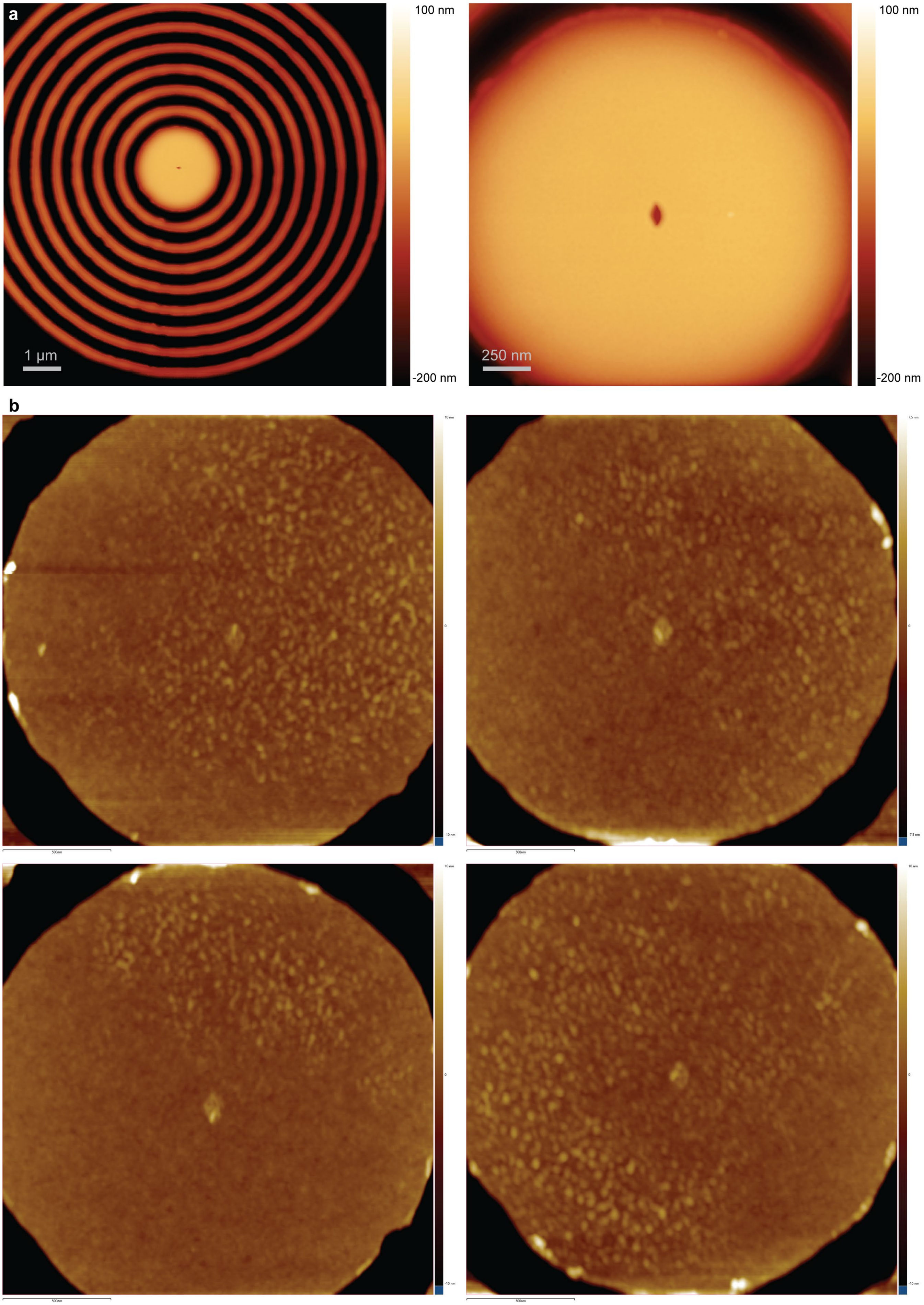
a
100 nm
-200 nm
1 µm
100 nm
250 nm
-200 nm
b

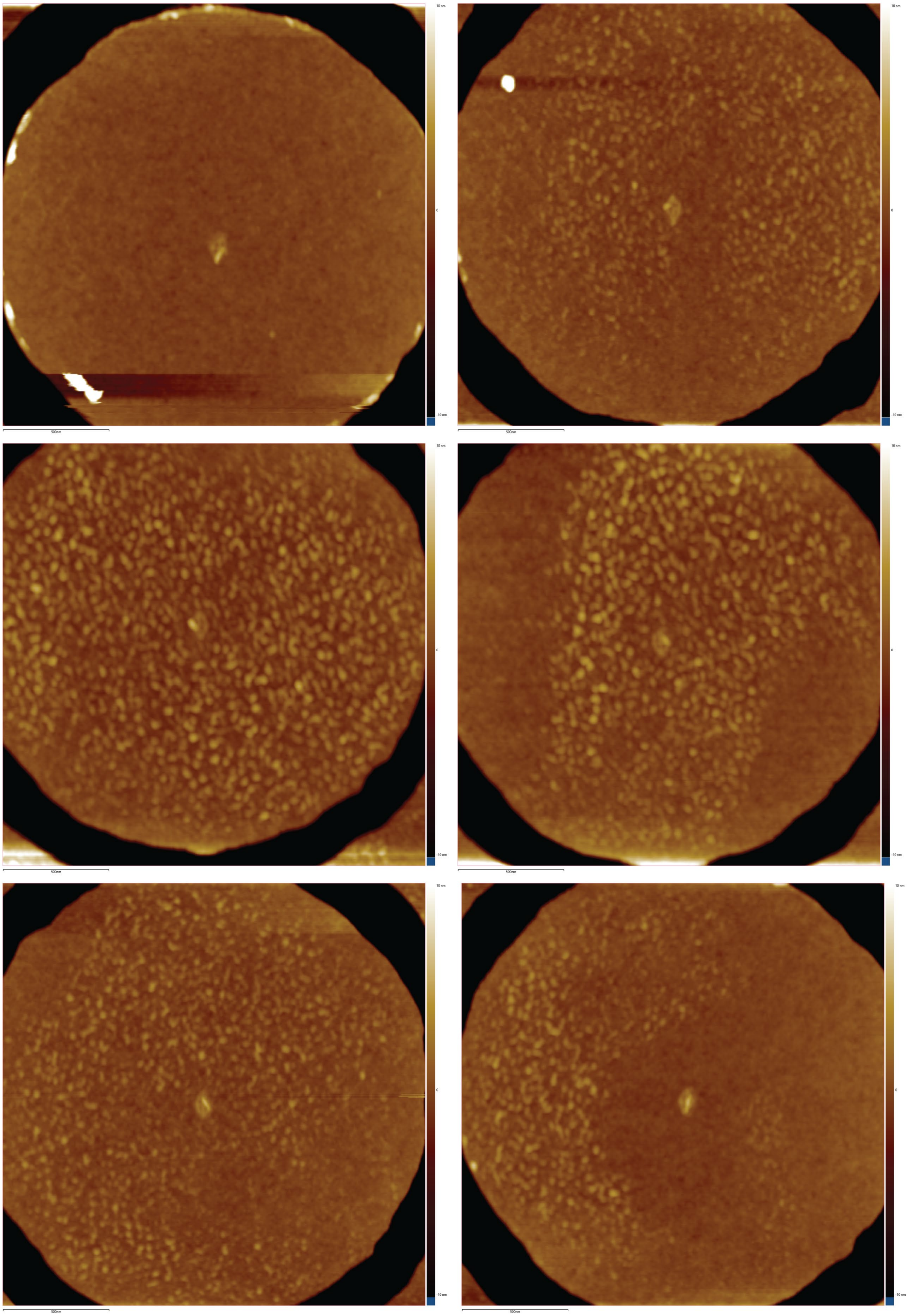

**Supplementary Fig. 47 | TMOP-directed origamis with a single QR at the center of $SiN_x$ bullseye cavities. a**, AFM images of single PMMA wells at the center of bullseye structures. **b**, AFM images of TMOP-directed single origami-QRs at the center of bullseye structures.

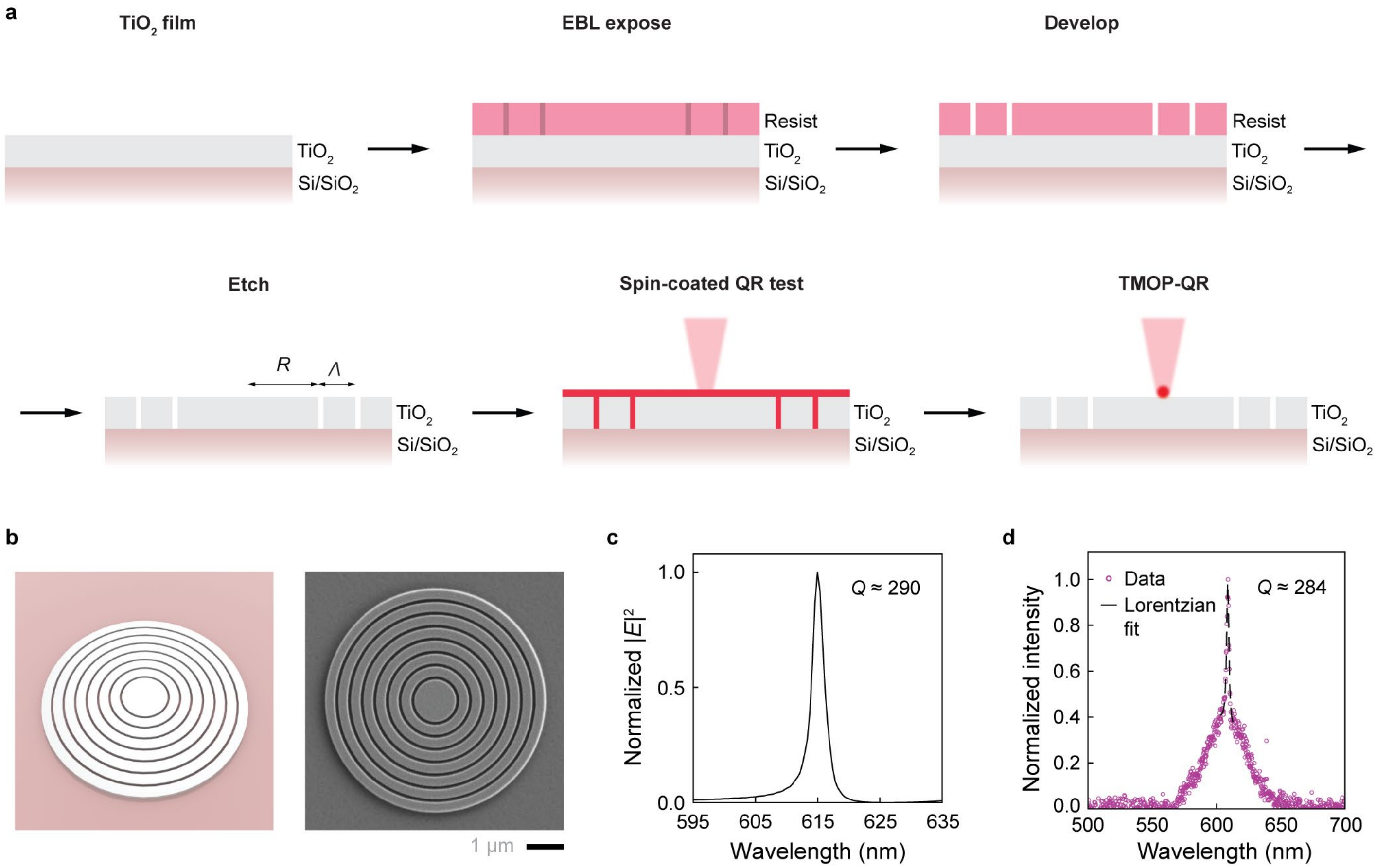


**Supplementary Fig. 48 | Fabrication and optimization of $TiO_2$ bullseye optical cavities. a**, Schematics of the fabrication process. Parameters *R*, *Λ*, and duty cycle were swept in the fabrication of the bullseye cavities. Fabrication steps are described in Methods. **b**, Schematic and SEM image of a fabricated $TiO_2$ bullseye cavity (100 nm thick, 570 nm eye radius). **c**, Simulated local electric field intensity $|E|^2$ versus wavelength at the cavity center of the target bullseye cavity (570 nm eye radius, 343 nm period, 0.79 duty cycle). **d**, PL spectrum (purple) and Lorentzian fit (black) of the representative cavity (570 nm eye radius, 363 nm period, 0.71 duty cycle) coated with a QR film, showing a strong emission resonance ($Q \approx$ 284).

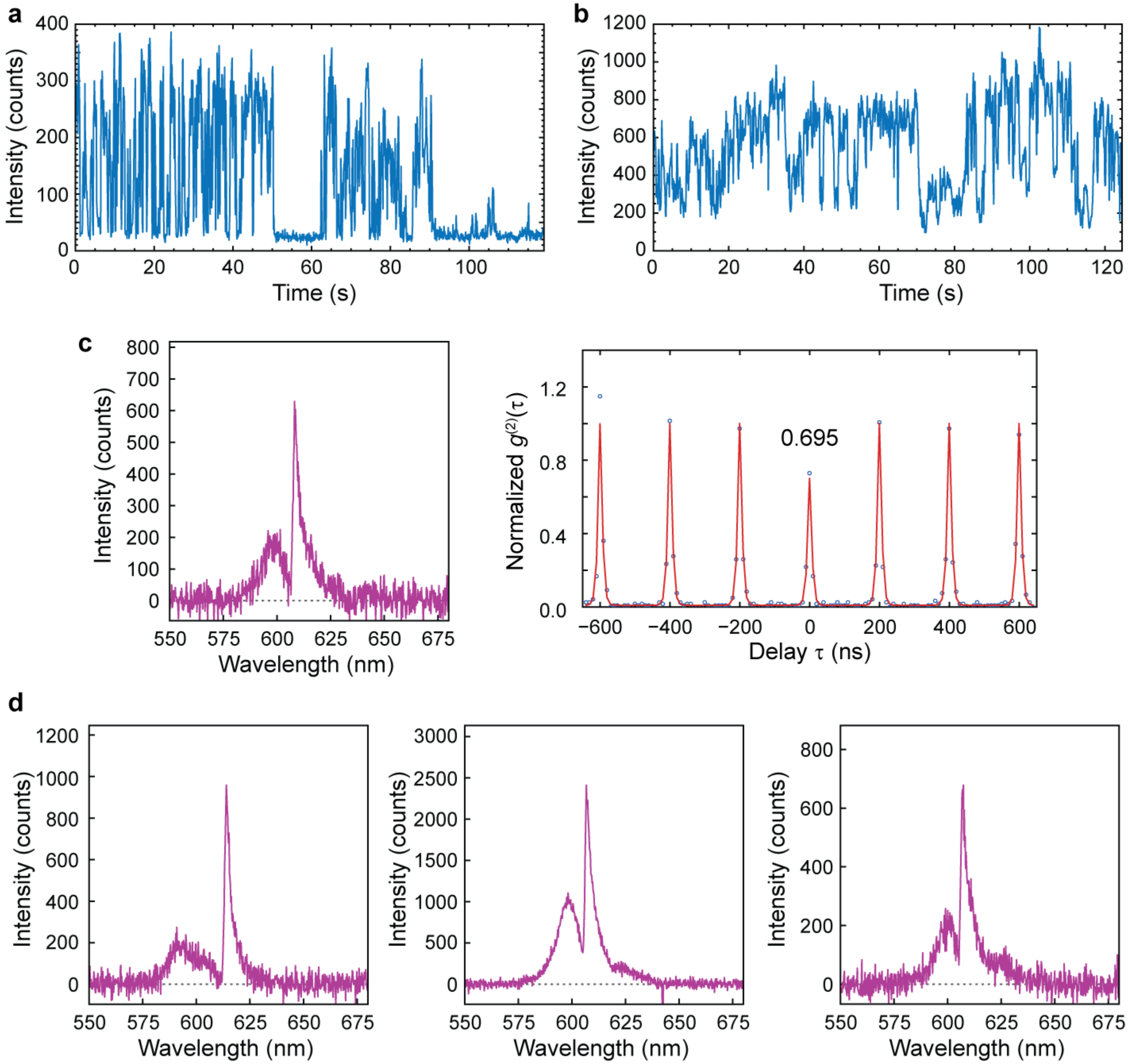


**Supplementary Fig. 49 | Additional optical characterizations of cavity-coupled QRs. a**, Intensity trace of cavity-coupled single-photon emission from the same device in Fig. 5f,g. **b**, Intensity trace of cavity-coupled QR-cluster emission (multiple QRs placed without DNA origami templating at the center of a $TiO_2$ bullseye cavity). **c**, PL spectrum and second-order correlation trace of the cavity-coupled QR-cluster emission from the same device in (b). The Fano line shape in the spectrum indicates cavity coupling, and $g^{(2)}(0) > 0.5$ suggests emission from multiple QRs at the cavity center. **d**, Additional spectra of cavity-coupled QR clusters showing Fano line shapes. QR cluster placement achieved by placing the same DNA-functionalized QRs on EBL-patterned $TiO_2$ bullseye chips without loading them onto origami templates following the protocol in Methods.

**Supplementary Table 1 | Sequences of staple strands modified with binding overhangs.** Binding overhang sequences are shown in red; the schematic below shows the staple positions.

| Origami | Staple | Sequence |
|---|---|---|
| Rhombic origami | 63-BS | TATATTTTGTCAATTTCCTGTAGCCAGCTCAATATCAACGCTTTTTTTTTTTTTTTTTAGAGAACCTGGG |
| | 68-BS | AATCCAAAAAACAGTTAACCAATAGGAAGAAGGGTTTGCTATTTTTTTTTTTTTTTTTAGAGAACCTGGG |
| | 119-BS | GAGAGTCTGGACCGACCAGAGGCATTTTCGACCAGTTACATCATATTTTTTTTTTTTTTTTAGAGAACCTGGG |
| | 120-BS | ACTAGCATAGTTAAACAAAAGGTAAAGTATCTTACAATCCTGTTTTTTTTTTTTTTTTAGAGAACCTGGG |
| | 121-BS | AGCCCCATCGCAAGACATGTTCAGCTAAAGATTAGTAGAACCTTTTTTTTTTTTTTTTAGAGAACCTGGG |
| Ladder-shaped origami | 115-BS (site1) | TTTAGCGATTTTCAGAAAACTTTTTCAACGTGTGACATATGGTTTTTTTTTTTTTTTTTTAGAGAACCTGGG |
| | 138-BS (site1) | TTTTTTGGCTGTCTTCGTAAAGGAAACATGAAAGTAGAATTAGGATAAGTTTTTTTTTTTTTTTTTAGAGAACCTGGG |
| | 139-BS (site1) | GCCGTTTTTAACCTCCTAACGAGCGTCTTTGAAAATTTAAATAATTTTTTTTTTTTTTTTAGAGAACCTGGG |
| | 103-BS (site2) | CTCATGGCTTGCCTAGAGGGTAGCTATTCAATAGGATCATCATATTTTTTTTTTTTTTTTAGAGAACCTGGG |
| | 107-BS (site2) | AGCGAAAGATTTTCAAAACAGAAATAAAGGATTATCAAGTTTTTTTTTTTTTTTTTTTAGAGAACCTGGG |
| | 130-BS (site2) | ATAACATCAAAATACCGCCAACAGAGATAGCGGAATTAACGCCATTTTTTTTTTTTTTTTAGAGAACCTGGG |

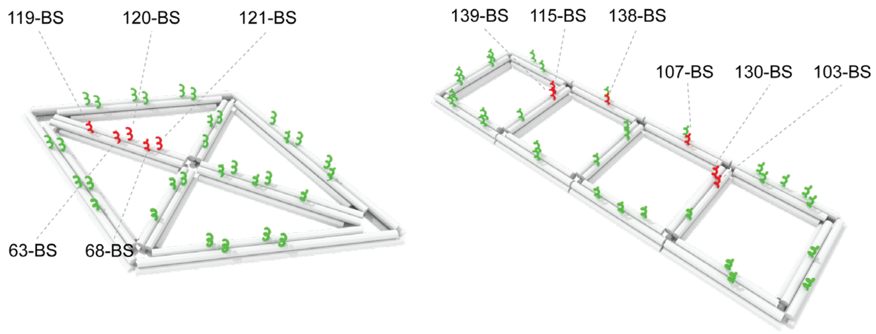


**Supplementary Table 2 | Fitted optical parameters for quantum rod coupled to bullseye cavity**

| Parameter | Description | Value |
|---|---|---|
| $\hbar\omega_d$ | Dipole resonance frequency | 2.11 eV |
| $\hbar\omega_c$ | Cavity resonance frequency | 2.08 eV |
| $2\hbar\Gamma_d$ | Dipole linewidth | 34.4 meV |
| $2\hbar\kappa$ | Cavity linewidth | 6.69 meV |
| $F_0$ | Coupling parameter | 0.51 |
| $\Delta\phi$ | Phase offset | 64° |
